# Overweight status drives early tumor microenvironment reprogramming in pancreatic ductal adenocarcinoma: a cell-type-resolved Bayesian hierarchical modeling and interactome analysis

**DOI:** 10.64898/2026.05.14.721695

**Authors:** Arun Viswanathan, Justin Seby, Kuzhuvelil B. Harikumar

**Affiliations:** Cancer Research Program, BRIC-Rajiv Gandhi Center for Biotechnology (BRIC-RGCB), Thiruvananthapuram, Kerala State, 695014, India; Manipal Academy of Higher Education (MAHE), Manipal, Karnataka State, 576104, India; SciLifeLab, Department of Oncology-Pathology, Karolinska Institutet, Tomtebodavägen, 23A, 17121, Solna, Sweden

**Keywords:** Pancreatic ductal adenocarcinoma, Obesity, Tumor microenvironment, Bayesian hierarchical modeling, Immune deconvolution, Cell-cell interactions

## Abstract

Obesity significantly increases the risk of pancreatic ductal adenocarcinoma, yet a comprehensive understanding of obesity-driven tumor microenvironment remodeling remains incomplete. We developed a cell-type-resolved transcriptomic model using bulk RNA sequencing data from the Clinical Proteomic Tumor Analysis Consortium cohort (n=140) stratified by body mass index. A custom functional gene signature database covering 65 immune and stromal cell types was manually constructed and validated through large-language model-assisted review. Cell-type-specific expression profiles were derived using BayesPrism deconvolution with matched single-cell RNA sequencing references, enabling high-resolution quantification of signature activity within each cell type. Body mass index-associated signatures were identified using a machine learning framework. Bulk pathway analysis showed deterioration of extracellular matrix homeostasis and primary immunodeficiency pathways with rising body mass index. Bayesian hierarchical modeling revealed cell-type-specific, non-linear dynamics: stromal populations in overweight individuals underwent coordinated extracellular matrix remodeling and inflammatory signaling that stabilized rather than intensified in obesity, while CD8-positive T cells showed dose-dependent activation progressing to chronic exhaustion. Spatial analysis showed that stromal-trapped CD8-positive T cells were positioned progressively closer to the cytokeratin-positive tumor boundary with rising body mass index. These findings establish overweight status as a critical tipping point in tumor microenvironment reprogramming, suggesting that early metabolic interventions may prevent irreversible microenvironmental deterioration.

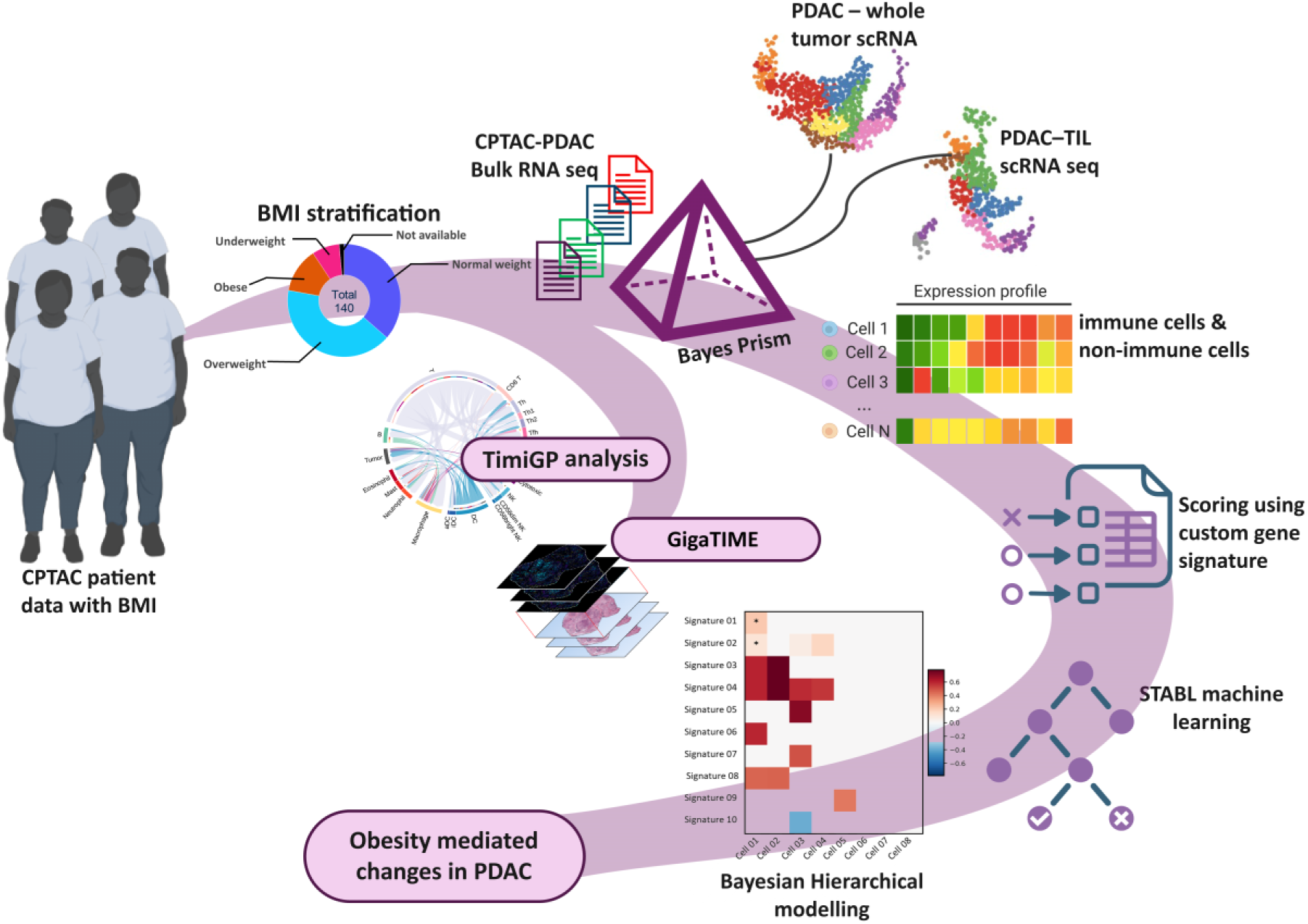

## INTRODUCTION

Globally, pancreatic ductal adenocarcinoma (PDAC) is among the most dangerous malignancies, distinguished by one of the lowest five-year overall survival rates, a status it has maintained for decades. ^1^ The malignancy presents significant challenges to diagnose, treat, and manage because of its high-functioning cancer-driving mutations, highly evolved tumor desmoplasia, unusual immune suppressive milieu, and its symptomatically silent nature (2–6).

Among the modifiable risk factors for PDAC, obesity has emerged as a significant driver for disease incidence, prognosis, and survival ^7–13^. Several epidemiological studies and meta-analyses have consistently linked Body Mass Index (BMI) with pancreatic cancer across different demographics ^14–21^. Mechanistically, obesity induces chronic systemic low-grade inflammation, altered adipokine signaling, and metabolic malfunctions that aid and abet malignancy ^22^. Within the tumor microenvironment (TME), these systemic changes translate into extracellular matrix (ECM) remodeling and immune cell dysfunction ^8,23^. Although the complex interplay of obesity in cancer has been widely recognized, many studies still concentrate on a binary understanding of obese and non-obese cancer dynamics ^24^. Such studies often overlook the intermediate OW phase, which might act as a critical tipping point where the earliest reprogramming of stroma and immune compartment is set in motion ^25^. Because current research often focuses on severe obesity, a gap remains in our broader understanding of how dynamic BMI levels shape the immune and stromal compartments in PDAC.

To address this complexity, we utilized transcriptomic data from the Clinical Proteomic Tumor Analysis Consortium (CPTAC-3) cohort to analyze 140 PDAC patients stratified by BMI. We developed a Bayesian hierarchical model for obesity-driven pancreatic cancer, supported by machine-learning-based screening of gene signatures. Our model framework allows for signature-level and cellular-level changes in BMI, both in categorical comparisons and in continuous PDAC modeling. To understand how obesity alters tissue architecture, we used gigaTIME, a multimodal AI that generates virtual multiplex immunofluorescence (mIF) from standard H&E slides ^26^.

Our findings challenge prevailing linear models of obesity-driven pancreatic cancer progression. Rather than a proportional intensification of gene signatures with increasing BMI, we identify OW status as a critical transition state in which stromal and immune reprogramming initiates and subsequently stabilizes or collapses in clinical obesity. This positions the OW state as a candidate window for metabolic intervention and argues for BMI-aware patient stratification in PDAC management.

## RESULTS

### The extracellular matrix exhibits distinct activation statuses among weight groups

We analyzed transcriptomic data 140 PDAC patients stratified by WHO BMI classification (N, n=51; OW, n=58; OB, n=18) using differential expression analysis, GSEA, and KEGG Pathview (Figure 1A)^27^. GSEA analysis revealed BMI-dependent enrichment patterns in ECM-related pathways (Figure 1B-E, Supplementary Figure 3D-H).

**Figure 1:**
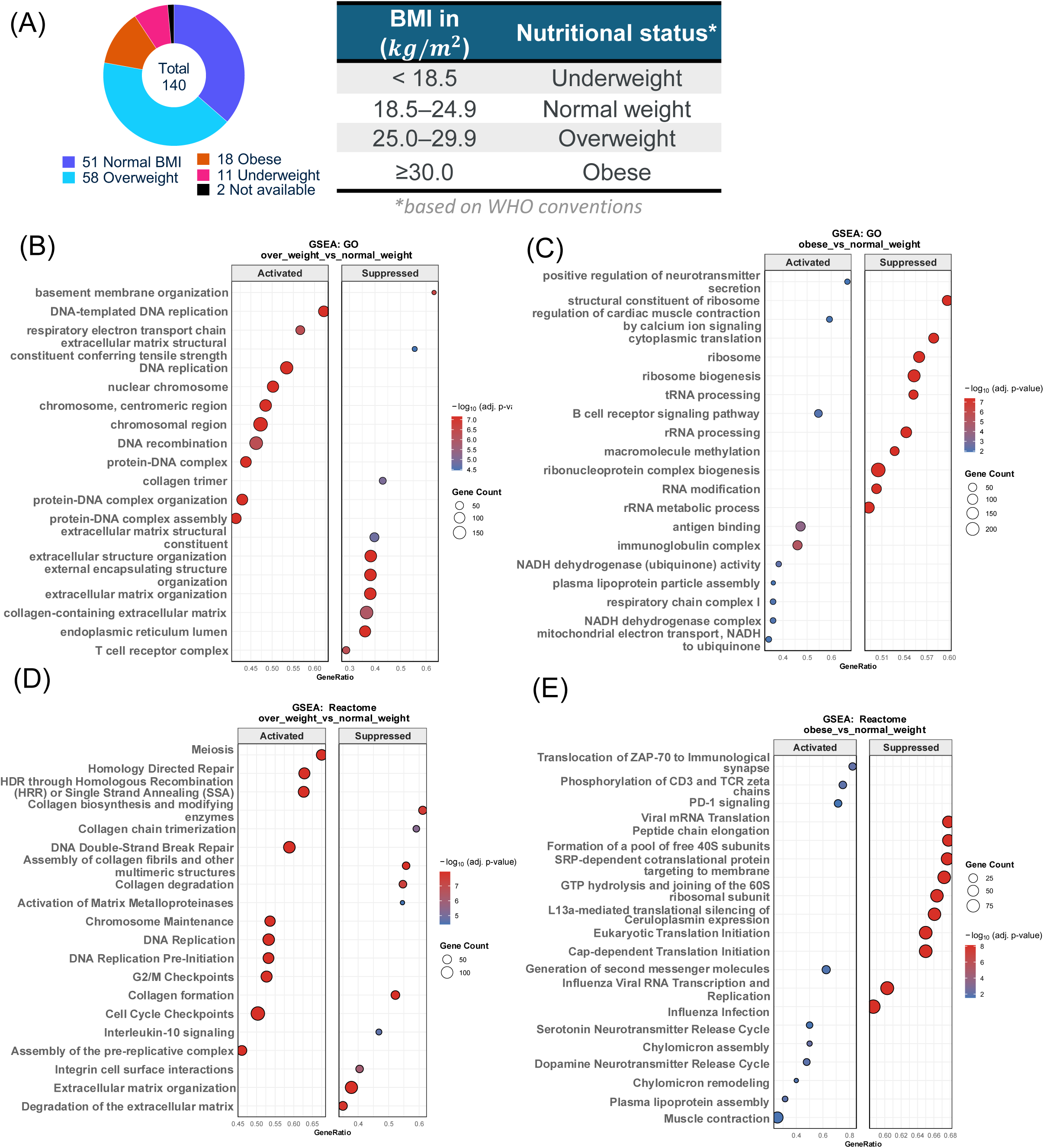
Transcriptomic profiles of pancreatic ductal adenocarcinoma (PDAC) stratified by BMI reveal differential immune and extracellular matrix pathways. (A) Distribution of 140 CPTAC pancreatic cancer patients across BMI categories per WHO classifications. (B-C) GSEA: GO bubble plots showing pathways differentially enriched between Overweight vs. Normal weight (B) and Obese vs. Normal weight (C) patients. (D-E) GSEA: Reactome pathways analysis comparing Overweight vs. Normal weight (D) and Obese vs. Normal weight (E) groups. Bubble size reflects the number of genes contributing to each term, and color indicates adjusted p-values. Note: In all GSEA plots, Activated indicates terms enriched in overweight/obese; Suppressed indicates terms enriched in normal weight patients. **Figure 1 Alt text:** A five-panel figure. Panel A shows a bar chart of 140 PDAC patients distributed across normal weight, overweight, and obese BMI categories. Panels B and C show bubble plots of Gene Ontology pathways enriched in overweight versus normal weight and obese versus normal weight comparisons respectively, with bubble size indicating gene count and color indicating adjusted p-value. Panels D and E show Reactome pathway bubble plots for the same two comparisons.

In the OW vs N comparison, N patients showed strong enrichment in ECM homeostasis pathways, including collagen synthesis, chain trimerization, degradation, and ECM organization, while OW patients exhibited broad structural ECM suppression (Figure 1B and 1D). Notably, no similar enrichment terms were identified in the OB versus N comparison (Figure 1C and E). In OB vs OW, obese patients showed enrichment of ECM terms, including organization and structural constituents (Supplementary Figure 3E). KEGG Pathwayview of the ECM-receptor interaction pathway (hsa04512) showed that OW patients exhibit broad downregulation of ECM components alongside selective upregulation of fatty acid transporter CD36, GPV, and specific integrin subunits (α7, α10, and β3), suggesting ECM modeling in OW coupled with lipid metabolic demands (Supplementary Figure 4A, 4B). OB patients showed specific upregulation of Thrombospondin (THBS), an established obesity biomarker ^28^, and partially re-acquired ECM/collagen organization signatures relative to OW individuals (Supplementary Figure 4B, 4C; Supplementary Figure 3E, 3H). This indicates that the ECM landscape has distinct remodeling modes based on BMI severity: homeostatic turnover in N, metabolic-coupled remodeling in OW, and a matricellular signaling phenotype in obesity. The latter aligns with the fibrosis-related matrisomal signatures in diet-induced obesity-driven PDAC ^29^.

### Progressive BMI-dependent TCR signaling dysfunction in PDAC

In GSEA, the T cell receptor complex (TCR) gene set was suppressed in OW patients compared with N patients (Figure 1B). KEGG Pathview analysis showed upregulation of proximal signaling receptors, receptors CD4/8, CD3ε, and CD3δ, and signaling kinases ZAP70 and IL-2-inducible T cell kinase (ITK) in high-BMI patients relative to N (Figure 2), aligning with OB vs N Reactome comparison, where ZAP-70 translocation to the immune synapse, CD3/TCR zeta chain phosphorylation pathways were enriched in OB patients (Figure 1E, 2). N patients, by contrast, showed higher expression of p21-activated kinase (PAK) and VAV, a guanine nucleotide exchange factor critical for actin remodeling during T cell activation ^30,31^. In OW vs N, alongside this disconnect in signaling execution, IL-2, IL-4, and IFNγ were elevated, while GM-CSF and TNFα were reduced in both the T cell receptor signaling pathway and the cytokine receptor interaction pathway (Supplementary Figure 5). This disconnect suggests that, while T cells in OW patients receive robust antigenic signals, they are unable to mount an effective immune response, resulting in abortive activation.

**Figure 2:**
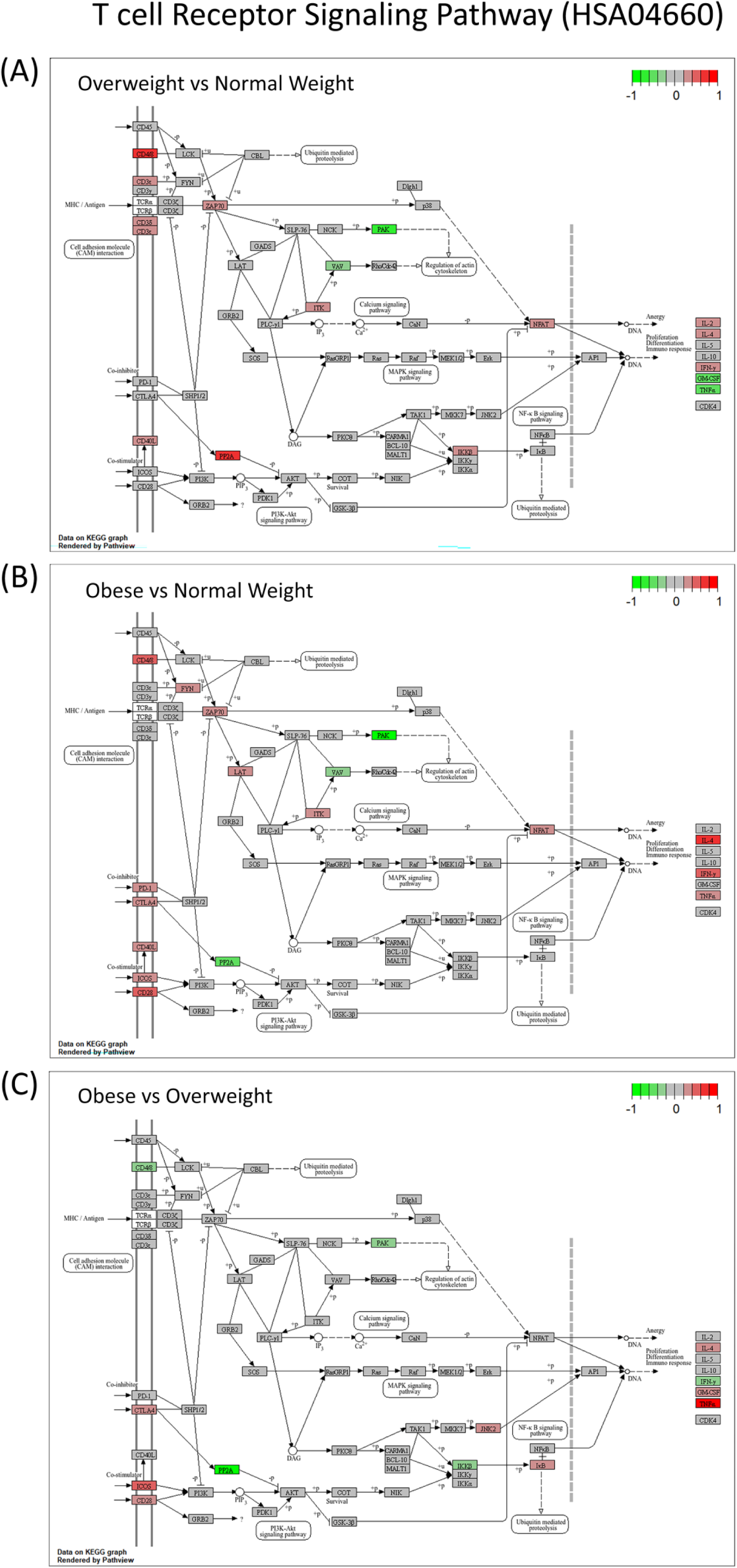
T Cell Receptor Signaling Pathway (HSA04660) expression in pancreatic cancer across BMI categories. KEGG pathway analysis showing differentially regulated genes in T cell receptor signaling between overweight vs. normal weight (top), obese vs. normal weight (middle), and obese vs. overweight (bottom) pancreatic cancer patients. Color scale indicates gene expression levels (green: downregulation [-1]; red: upregulation [+1]). **Figure 2 Alt text:** Three KEGG pathway diagrams of T cell receptor signaling arranged vertically, comparing overweight versus normal weight (top), obese versus normal weight (middle), and obese versus overweight (bottom) pancreatic cancer patients. Genes are color-coded green for downregulation and red for upregulation on a scale of minus one to plus one.

In obesity, contrasting N patients, proximal kinase engagement intensified alongside simultaneous upregulation of PD-1, CTLA-4, co-stimulatory molecules (CD40L, ICOS), and TNFα, indicating a chronic stimulation-exhaustion phenotype rather than functional silencing (Figure 2). The OB vs OW comparison revealed progressive deterioration of TCR signaling, with downregulation of PAK and PP2A in OB patients. Crucially, this functional deterioration in obesity tracks with the intensification of metabolic dysregulation. Metabolic dysregulation co-occurred with this functional deterioration in OB patients, with plasma lipoprotein assembly, NADH dehydrogenase activity, and chylomicron remodeling and assembly pathways enriched relative to N (Figure 1C, E). These bulk observations captured the trajectory of TCR dysfunction across BMI strata but could not resolve its cellular basis without deconvolution.

### BMI-associated alterations in primary immunodeficiency pathways

Pearson correlation analysis between BMI and CPTAC-3 RNA-seq gene expression in LinkedOmics (n=140) identified Primary Immunodeficiency (hsa05340) as the pathway most significantly associated with increasing BMI (Supplementary Figure 6). This pathway was detected in the pairwise OW vs N differential expression analysis but did not reach conventional significance thresholds (𝑎𝑑𝑗. 𝑝 𝑣𝑎𝑙𝑢𝑒 = 0.0195), demonstrating the added sensitivity of continuous BMI modeling over categorical comparisons. The enrichment does not suggest a lack of immune cells; instead, it reflects the dysfunctional, heightened activation state frequently seen in tumors associated with obesity ^32^.

Upon closer inspection of the pathway, OW patients showed a stoichiometric imbalance in the V(D)J recombination complex: RAG1 was upregulated, whereas its essential partner *RAG2* was downregulated, accompanied by upregulation of *CD3ε*, *CD3δ*, *RAG1*, *AIRE*, *CD8*, and *ZAP70* (Figure 3A) ^33^. In solid tumors, including PDAC, RAG expression likely denotes the presence of tertiary lymphoid structures; RAG1/RAG2 discordance, given that coordinated expression of both subunits is required for productive V(D)J recombination, thereby indicates defective tertiary lymphoid structure formation rather than active lymphoid maturation ^34^. However, *RAG2* was downregulated in OB patients, whereas *RAG1* overexpression was not detected in OB patients compared with N individuals alongside elevated CD8, ZAP70, and AIRE, indicating a transcriptionally active but developmentally impaired lymphoid state (Figure 3B). B-cell programs exhibited a bias towards terminal differentiation rather than de novo recruitment. OW patients demonstrated increased expression of components of class-switch recombination (*AID*, *CD40L*) and of activation and survival markers (*CD19*, *BAFFR*, *TACI*). λ5, a component of the pre-B cell receptor’s surrogate light chain, crucial for progression from pro-B to pre-B cell and allelic exclusion of immunoglobulin heavy chains, was decreased during early B cell maturation in OB patients, suggesting impaired early B-cell maturation (Figure 3B).

**Figure 3:**
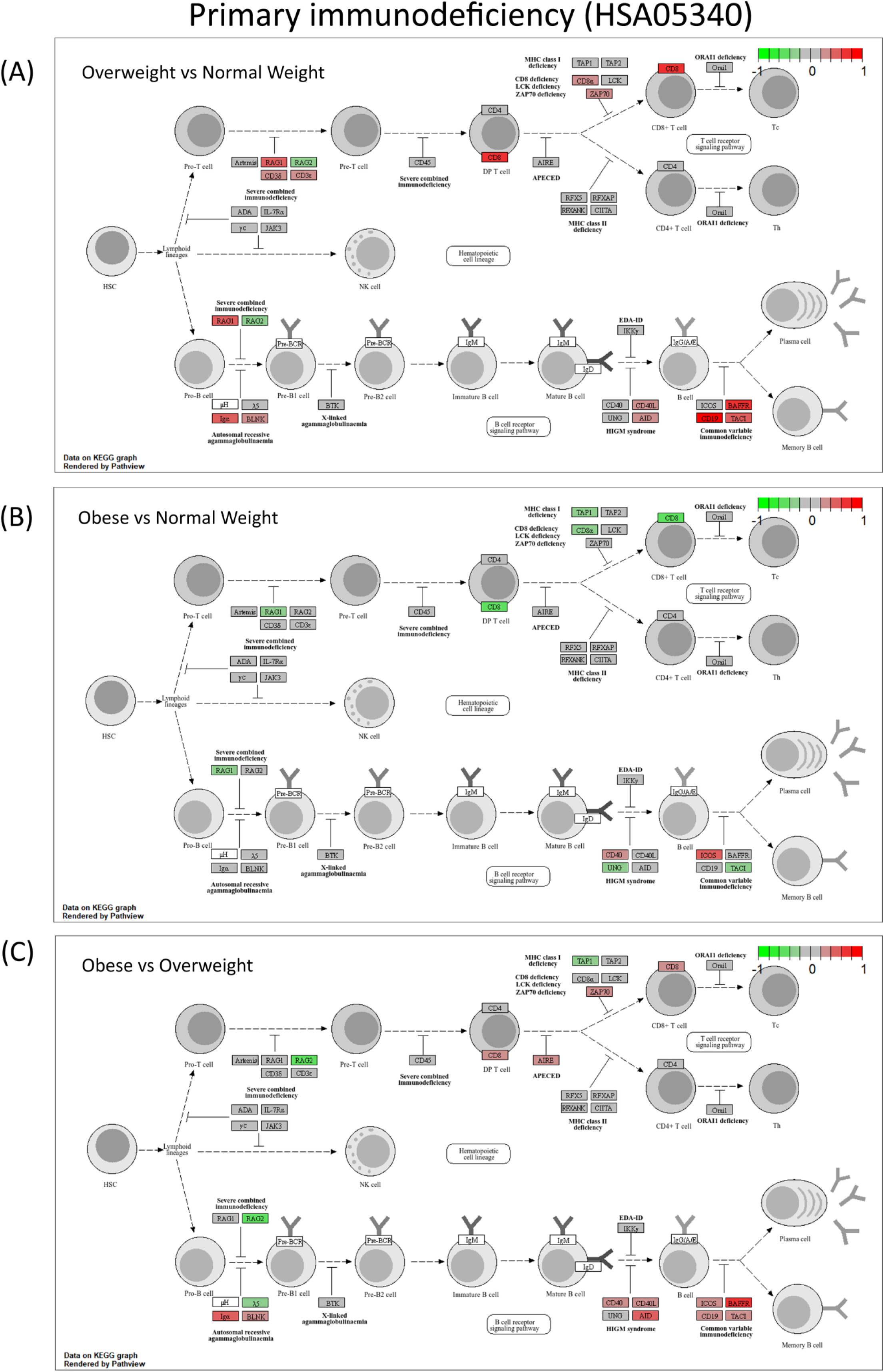
Primary Immunodeficiency (HSA05340) comparison across BMI groups in pancreatic cancer. KEGG pathway analysis showing differential gene expression in overweight vs. normal-weight (top), obese vs. normal-weight (middle), and obese vs. overweight (bottom) patients. Color scale indicates expression changes (green: downregulation [-1]; red: upregulation [+1]). **Figure 3 Alt text:** Three KEGG pathway diagrams of the primary immunodeficiency pathway arranged vertically, comparing overweight versus normal weight (top), obese versus normal weight (middle), and obese versus overweight (bottom) pancreatic cancer patients. Genes are color-coded green for downregulation and red for upregulation on a scale of minus one to plus one.

ssGSEA of ImmPort gene set (Supplementary Figure 7) confirmed that the OW condition represents a unique immunological peak rather than a progressive transition ^35^. Humoral immune response and adaptive memory pathways were highest in OW patients relative to N (Supplementary Figure 7A, C, D). While OW-specific upregulation of the pathway that negatively regulates T cell activation-induced cell death indicated active retention of antigen-experienced lymphocytes through suppressed apoptosis, a protective mechanism that subsequently declined in the OB group (Supplementary Figure 7E). This functional enrichment, along with the transcriptional evidence of class-switch recombination and possible atypical TLS formation, adds weight to the argument that the OW microenvironment actively facilitates the retention of antigen-experienced lymphocytes.

Analysis of the global Antigen Processing and Presentation pathway (KEGG: hsa04612) indicated that obesity induces a divergent regulation of antigen presentation machinery, rather than a uniform collapse (Supplementary Figure 8). MHC class II was upregulated in both OW and OB patients relative to N, consistent with B cell-driven changes in the tumor microenvironment (Supplementary Figure 8A, B), while MHC class I was suppressed in both high-BMI groups. OW patients have preserved *CD8* and *KIR* expression compared with N and OB patients, suggesting that they still possess a cytotoxic transcriptomic profile, though its functional profile cannot be inferred from this data. In OB patients, *TAP1* (Transporter associated with Antigen Processing) was specifically reduced relative to OW patients alongside lower CD8A expression, indicating concurrent impairment of intracellular antigen loading and cytotoxic T cell availability (Supplementary Figure 8C). The downregulation of UNG and TACI in OB compared to OW confirms a gradual decline in B cell activation ability. Collectively, these results reveal a non-linear immune development pattern: OW status leads to a dysfunctional yet transcriptionally active lymphoid structure, whereas clinical obesity promotes immune evasion by impairing antigen presentation. Understanding the cellular foundation of this trajectory necessitated deconvolution.

### Cell-type–specific functional reprogramming across BMI in PDAC

Bulk analysis revealed BMI-dependent transcriptional changes that conventional pathway-level methods could not assign to specific cell populations. We deconvoluted the bulk RNA-seq data using BayesPrism with matched single-cell references, finding no significant compositional changes across BMI groups except in naïve B cells and CD8 T memory cells (Supplementary Figure 9D-E, Supplementary Figures 10 and 11). Followed by the custom signature’s z-score calculation and screening of the BMI-associated signature using the *Stabl* ML algorithm (Supplementary Figure 7A-C). This strategy addressed the statistical challenge of comparing high-dimensional features across a relatively small, BMI-stratified cohort.

Full convergence diagnostics are reported in Supplementary Table 2. Three comparisons presented challenges and are handled as follows: OW vs N in the immune coarse compartment failed convergence and is treated as exploratory throughout; OW vs N in the non-immune compartment showed marginal convergence, with OB vs N used as the primary comparison; and the continuous immune fine BMI slope showed partial convergence, with conclusions drawn only from cell types that met criteria.

### Overweight-driven non-immune reprogramming precedes attenuation in obesity

The non-immune compartment is defined by major cell types, including fibroblasts and tumor cells. We observed a distinct shift in non-immune cell signatures across BMI comparisons, with most changes pointing towards increased stromal activation and metabolic changes in the epithelial population. Fibroblast accounted for the largest share of credible signatures (HDI excluding zero), with changes in the positive direction (Figure 4A, B, C, and G). Given the Marginal convergence status for fibroblasts in OW vs N, these results are corroborated by the OB vs N comparison, which converged cleanly (𝑅^ = 1.003, ESS > 1,200) and showed a near-identical directional pattern. In the OW vs N comparison, fibroblasts show feature-level upregulation of ECM remodeling, fibrosis, inflammatory, and metabolic signaling pathways, an activation that was sustained and extended in OB vs N. The pooled cell-type posterior distribution was positive in both comparisons, with the OB vs N estimate reaching primary convergence status, providing additional support for the fibroblast signal independent of the marginal OW vs N result (Figures 4E, 4F). Fibroblasts showed the most credible upregulated signatures in OB vs N (Figure 4I); however, the effect sizes of these signatures are minimal, suggesting the changes are low magnitude but widespread. (Figure 4G, H). No fibroblast or broader stromal signature reached credibility in the OB vs OW comparison, which returned only 9 credible features in total, distributed across tumor epithelial, normal ductal, mast, and neural populations (Figure 4C). The observed trend suggests that, rather than being a constant linear response to rising BMI, stromal remodeling is an early event triggered during the initial phase of weight gain.

**Figure 4:**
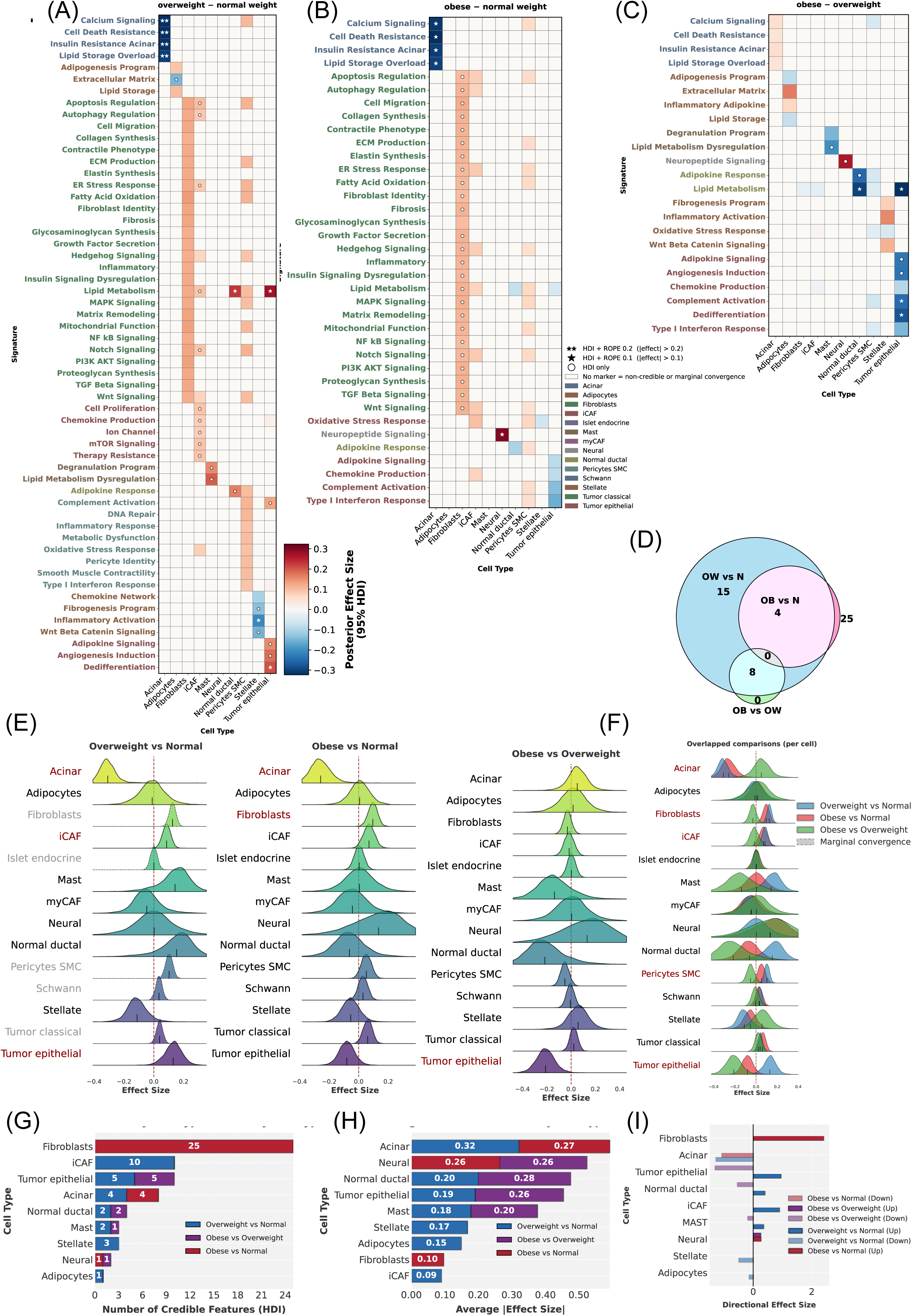
Bayesian hierarchical modeling of non-immune cells in PDAC. Heatmap visualization of posterior mean effect sizes for functional gene signatures across BMI comparisons: (A) Overweight vs. Normal, (B) Obese vs. Normal, and (C) Obese vs. Overweight. Rows represent unique functional signatures, color-coded by their associated cell type. Statistical credibility is denoted by markers: Double Star (★★) denotes strong practical significance (95% HDI excludes 0 and >95% of the posterior distribution falls outside a ROPE of ±0.2); Single Star (★) denotes moderate practical significance (95% HDI excludes 0 and >95% of the posterior distribution falls outside a ROPE of ±0.1); White Circle (○) denotes statistical credibility only (95% HDI excludes 0). (D-E) Cellular-Level Posterior Distributions. (D) Ridge plots displaying the posterior distributions of pooled cell-type-level effects, separated by comparison. (E) Overlaid ridge plots demonstrating the shift in cellular state distributions across all three comparisons. (F) Stacked bar plot quantifying the number of credible features (𝐻𝐷𝐼 ≠ 0) per cell type, stratified by comparison. (G) Stacked bar plot showing the average magnitude of effect sizes per cell type, highlighting compartments with the most substantial transcriptional shifts. (H) Directionality of effect size changes. The count of significantly upregulated (positive) and downregulated (negative) features across comparisons. (I) Venn diagram illustrating the overlap of credible functional signatures shared between the three comparisons. **Figure 4 Alt text:** A nine-panel figure showing Bayesian hierarchical modeling results for non-immune cells. Panels A, B, and C are heatmaps of posterior mean effect sizes for overweight versus normal, obese versus normal, and obese versus overweight comparisons respectively, with rows representing functional signatures color-coded by cell type and statistical credibility marked by stars and circles. Panel D shows ridge plots of pooled cell-type posterior distributions separated by comparison. Panel E shows overlaid ridge plots across all three comparisons. Panel F is a stacked bar plot of credible feature counts per cell type. Panel G is a stacked bar plot of average effect size magnitude per cell type. Panel H shows directional bar plots of upregulated and downregulated features. Panel I is a three-way Venn diagram of credible signatures shared across comparisons.

All four acinar signatures in OW vs N and OB vs N exceeded the ROPE=0.1 practical significance threshold, making them the strongest and most practically meaningful effects in the non-immune compartment. Acinar cells exhibited decreased calcium signaling, increased resistance to cell death, insulin resistance, and enhanced lipid storage signatures. Acinar cells showed a negative pooled posterior effect size in both high-BMI vs N comparisons; in contrast, OB v OW showed a density centered near zero (Figure 4E and F), consistent with a suppression established at OW that does not intensify further with additional weight gain.

Tumor epithelial cells (neither classical nor basal-like) showed metabolic and survival adaptations during OW transition, with upregulation of lipid metabolism, complement activation, angiogenesis signaling, and adipokine signaling (Figure 4B). The dedifferentiation signature, indicative of epithelial plasticity, was elevated in OW relative to N but decreased in OB patients (Figure 4B, C), suggesting that the OW tumor microenvironment promotes epithelial plasticity, whereas the obese microenvironment may select for a more static phenotype. Tumor classical cells showed a small positive pooled posterior across comparisons. The intermediate epithelial signature activity was elevated in OW vs N, then flipped to negative in OB vs N and OB vs OW, with the OB vs N estimate near zero (Figure 4E, F). A non-linear trajectory in which the effect peaks at moderate adiposity and attenuates in clinical obesity.

Mast cells show increased degranulation and alterations in lipid metabolism in the OW vs N comparison, which decline in OB to OW (Figure 4A, C). Neural cells showed increased neuropeptide signaling in OB compared to N and OW, indicating that neural activation is specific to obesity rather than the earlier OW transition. Neural and acinar cells showed the greatest change in effect size, followed by normal ductal cells (Figure 4H). iCAF signatures showed credibility only in OW vs N comparisons and disappeared in higher-BMI comparisons, reinforcing the OW-specific nature of inflammatory CAF activation.

Overall, OW patients exhibited large, distinct changes, with OB patients displaying a subset of these altered signatures, along with a few unique signatures of their own (Figure 4D). No distinct differences were observed between OB and OW patients, except for the reversal of effects established in the OW non-immune milieu, suggesting that the most cellular reprogramming occurs during the transition from N to OW, establishing a window of metabolic plasticity that closes as obesity becomes chronic.

### Non-linear immune remodeling across BMI reveals lineage-specific activation and collapse

The immune course compartment captures broad lineage-level populations, including B cells, CD4+ and CD8+ T cells, NK cells, and monocytes. BMI. OB vs N constitutes the primary comparison for this compartment; OW vs N did not meet convergence thresholds and is treated as exploratory throughout, with cell-type posterior means reported as directional signals only (Supplementary Table 2).

In OB vs N, B cells exhibit a strong increase in stress-related signatures (Cellular stress, ER stress, Oxidative stress, DNA repair, autophagy regulation, and Hypoxia response signature) alongside extensive activation of core B cell functions, including antibody machinery, antigen presentation, and signaling upregulation of metabolic programs including lipid processing, glycolysis, and mitochondrial activity (Figure 5A and B). The upregulation of both stress and metabolic signatures is consistent with metabolic reprogramming to sustain chronic inflammatory demand ^36^. The pooled cell-type posterior mean is above zero, indicating a broad and non-selective activation state at the signature level (Supplementary Figure 12 A, B). The OW vs N posterior mean was directionally consistent, suggesting this activation may begin before clinical obesity, though this cannot be confirmed from an exploratory comparison.

**Figure 5:**
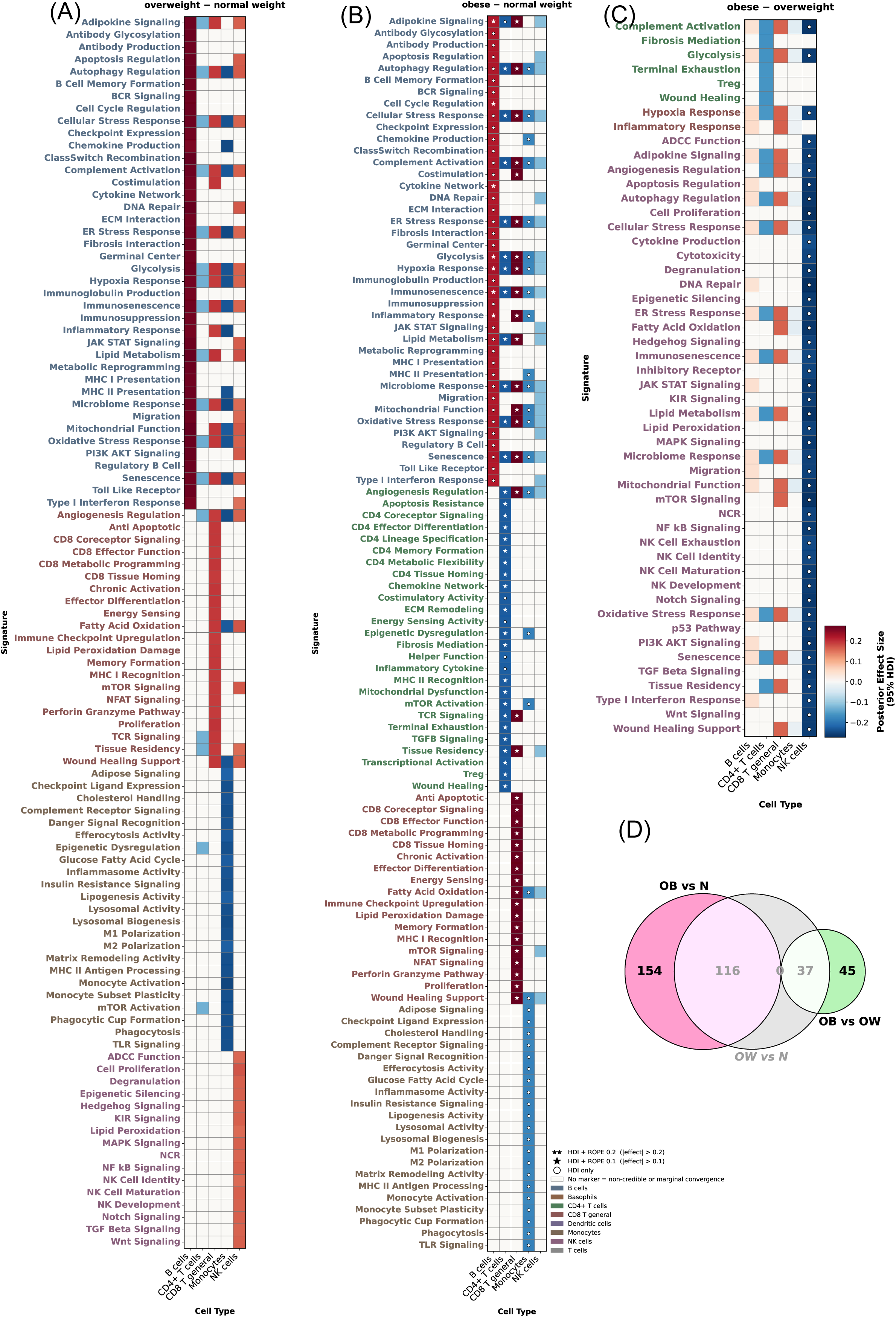
(A–C) Bayesian Evaluation of immune course compartment in PDAC. Heatmap visualization of posterior mean effect sizes for functional gene signatures across major immune compartments (B cells, CD4+/CD8+ T cells, NK cells, Monocytes, and Dendritic cells). Panels display transcriptional signatures in (A) Overweight vs. Normal, (B) Obese vs. Normal, and (C) Obese vs. Overweight comparisons. Rows represent unique functional signatures, color-coded by their associated cell type, and columns denote broad immune cell lineages. Statistical credibility and practical significance are indicated by markers: Double Star (★★) denotes strong practical significance (95% HDI excludes 0 and >95% of the posterior distribution falls outside a ROPE of ±0.2); Single Star (★) denotes moderate practical significance (95% HDI excludes 0 and >95% of the posterior distribution falls outside a ROPE of ±0.1); White Circle (○) denotes statistical credibility only (95% HDI excludes 0). (D) Venn diagram illustrating the intersection of credible functional signatures shared across the three comparisons. **Figure 5 Alt text:** A four-panel figure showing Bayesian evaluation of the immune coarse compartment. Panels A, B, and C are heatmaps of posterior mean effect sizes for overweight versus normal, obese versus normal, and obese versus overweight comparisons across B cells, CD4-positive and CD8-positive T cells, NK cells, monocytes, and dendritic cells, with statistical credibility indicated by stars and circles. Panel D is a three-way Venn diagram of credible signatures shared across the three comparisons.

CD4+ T cells showed no credible feature-level differences in OW vs N. In OB vs N, all 38 signatures were credibly downregulated, with posterior means in narrow range (Figure 5B, 5C, Supplementary Figure 12 A, B). The suppression was non-selective across siz functionally distinct categories: core helper functions including MHC II recognition, costimulatory activity and transcriptional activations; signaling networks including TCR, TGFβ, chemokine, and mTOR; lineage and differentiation programs including effector differentiation, lineage specification, tissue homing, and coreceptor signaling; metabolic signatures including glycolysis, lipid metabolism, energy sensing, and metabolic flexibility; and cellular stress responses; and subtype programs, including Treg signatures, memory formation, and terminal exhaustion. The equivalence of suppression across all programs indicates total functional collapse rather than a phenotypic shift. The pooled cell-type posterior shifted to negative in OB vs N, consistent with an obesity-threshold effect on CD4+ T cell transcriptional activity (Supplementary Figure 12 A, B).

CD8+ T cells showed credible upregulation across all measured signatures in OB vs N (Figure 5B). Upregulated programs include effector function, TCR signaling, coreceptor and MHC I signatures, metabolic reprogramming including glycolysis, fatty acid oxidation, mitochondrial function, and energy sensing, chronic activation and immune checkpoint upregulation, and senescence and immunosenescence programs accompanied by elevated ER, cellular, and oxidative stress responses. The parallel rise in global activity of CD8+ T cells and senescence programs suggests that these cells are being driven towards a terminally differentiated phenotype under chronic TME stress. The pooled cell-type posterior mean in OB vs N was +0.233, with the OW vs N posterior mean pointing in the same direction (+0.111) but not meeting convergence criteria; this directional consistency is noted as a signal but not treated as confirmatory evidence of progressive intensification (Supplementary Figure 12 A, B).

In OB vs N, monocytes showed credible downregulation across all signatures with posterior means of −0.149 to −0.160 (Figure 5B). The uniformity of this suppression across biologically divergent programs, including TLR signaling, phagocytosis, M1 and M2 polarization, glycolysis, efferocytosis, and complement activation, argues against a phenotypic shift toward any specific monocyte lineage. Critically, even stress response programs, such as cellular stress, ER stress, and oxidative stress, are downregulated, suggesting that obese monocytes have lost the capacity to mount adaptive responses to the hostile TME rather than simply redirecting their activity. The simultaneous collapse of both polarization axes (M1 and M2) further supports this interpretation. These cells have not been reprogrammed toward a tolerogenic or pro-tumorigenic phenotype; they have effectively ceased to function across all measurable axes. The OW vs N cell-type pooled posterior cell-type mean was similarly negative (−0.140), but the onset timing of monocyte suppression could not be established from the current analysis (Supplementary Figure 12A, B).

In OB vs N, NK cells showed no credible feature-level differences, with the pooled cell-type posterior mean near zero (−0.098), leaving NK transcriptional programs in obese patients indistinguishable from those in N patients (Figure 5B, Supplementary Figure 12 A, B). The more informative signal, though preliminary, comes from OB vs OW, where 45 signatures were downregulated relative to the overweight state (posterior means: −0.181 to −0.204), across programs as functionally distinct as cytotoxicity, degranulation, mTOR signaling, mitochondrial function, KIR and NCR receptor signaling, and NK cell identity and maturation. As with monocytes, this uniformity argues against selective pathway suppression and points toward a global transcriptional deflation between the overweight and obese states. Critically, NK exhaustion, senescence, and immunosenescence signatures were among the downregulated features, indicating that the OB vs OW decline does not reflect classical exhaustion, in which effector function falls while exhaustion markers rise, but rather a simultaneous collapse of both functional and stress-adaptation programs. The OW vs N cell-type posterior mean was directionally positive, near zero at OB vs N, and credibly lower at OB vs OW, consistent with transient NK activation in the overweight state that does not persist into obesity, though this trajectory remains unconfirmed in a converged OW vs N model (Supplementary Figure 12A).

### Fine immune resolution reveals BMI-dependent threshold and reversal effects

Immune fine compartment resolves immune populations into granular subtypes, including naïve B cells, CD4+ T cell subsets (naïve, follicular helper, regulatory, and Th1), CD8+ T cell subsets (memory and exhausted, dendritic cell subtypes (plasmocytic and monocytic), classical and non-classical monocytes, TAM, γδT cells, MAIT cells, and basophils. Both OW vs N and OB vs N met convergence criteria in this compartment (Supplementary Table 2), lending credibility to directional comparisons across both transitions.

Naive B cells showed credible upregulation of all 18 measured signatures in both OW vs N (posterior means: 0.238 to 0.263) and OB vs N (posterior means: 0.245 to 0.273), with no credible OB vs OWdifferences, and represented one of the largest pooled cell-type effects in the compartment (cell-type posterior means: +0.252 and +0.260 respectively; Figures 6A, 6B, 6G). Naive B cells exhibit significant elevation of metabolic reprogramming, including OXPHOS, glycolysis, fatty acid oxidation, and lipid metabolism; stress and adaptive responses, including autophagy, oxidative stress, hypoxia, and p53 pathway; signaling programs including NF-κB, mTOR, MAPK, Notch, TGF-β, Hedgehog, and Type I interferon; and naive B cell identity and activation signatures. The near-identical effect sizes across OW vs N and OB vs N, and the absence of any credible OB vs OW differences, indicate that naive B cell activation is fully established at the OW stage and does not escalate further with additional weight gain.

**Figure 6:**
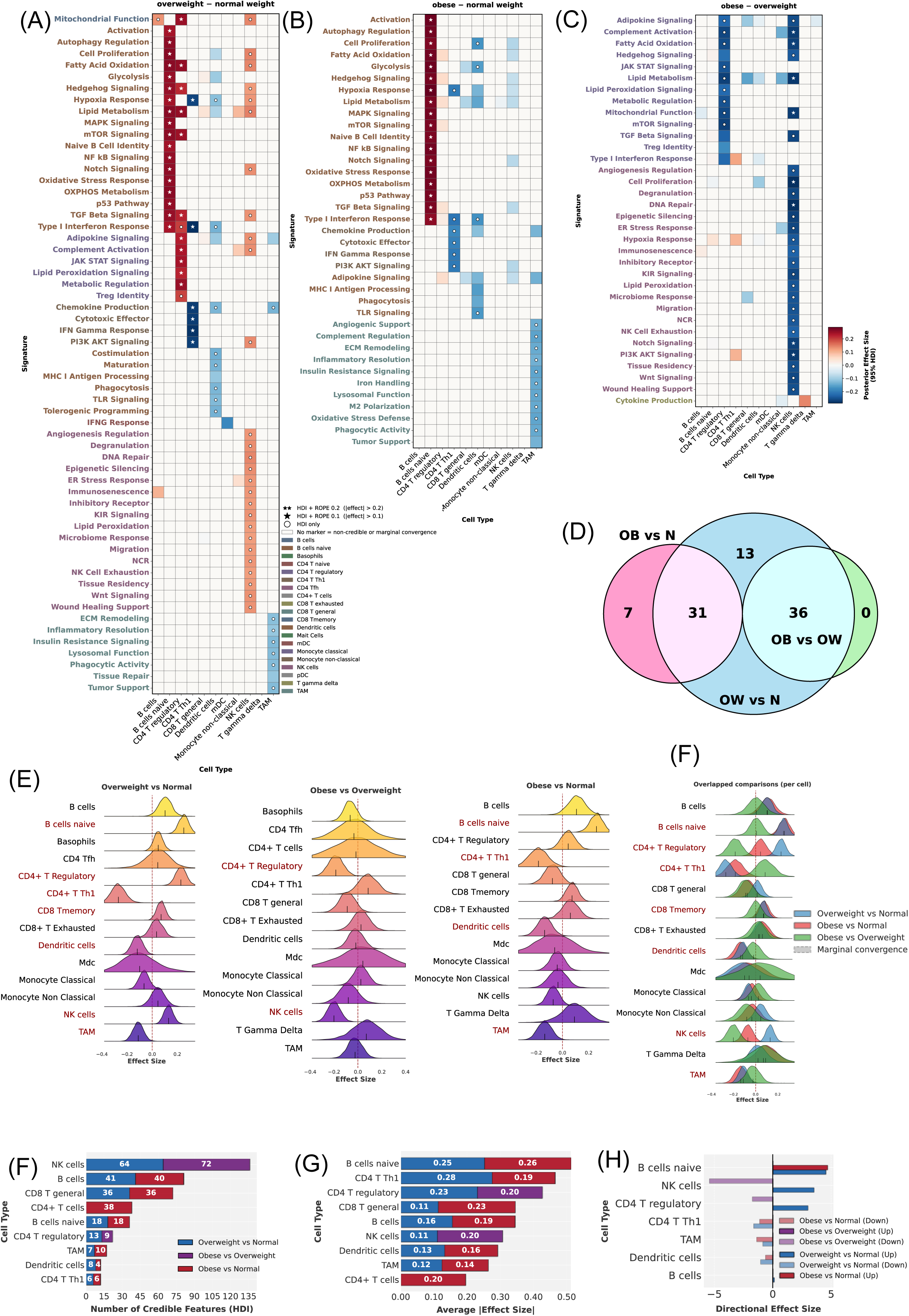
Bayesian Evaluation of the immune fine compartment in PDAC reveals Distinct Immune Dysregulation Patterns. **(A-C)** Landscape of State-Specific Immune Remodeling. Heatmap visualization of posterior mean effect sizes for functional gene signatures across granular immune cell states (e.g., CD8+ Tex/Tem, Treg, CD4+ Treg/Th1, TAM). Panels display the transcriptional shifts in **(A)** Overweight vs. Normal, **(B)** Obese vs. Normal, and **(c)** Obese vs. Overweight comparisons. Rows represent unique functional signatures, color-coded by their associated cell type, and columns denote specific immune cell states. Statistical credibility and practical significance are indicated by markers: Double Star (★★) denotes strong practical significance (95% HDI excludes 0 and > 95% of the posterior distribution falls outside a ROPE of ±0.2); Single Star (★) denotes moderate practical significance (95% HDI excludes 0 and > 95% of the posterior distribution falls outside a ROPE of ±0.1); White Circle (○) denotes statistical credibility only (95% HDI excludes 0). **(D)** Venn diagram illustrating the overlap of credible functional signatures shared across the three comparisons. **(E-F)** Cellular-Level Posterior Distributions in the immune fine compartment. **(E)** Ridge plots displaying the posterior distributions of pooled cell-type-level effects, separated by comparison. **(F)** Overlaid ridge plots demonstrating the shift in cellular state distributions across all three comparisons. **(G)** Stacked bar plot quantifying the number of credible features (𝐻𝐷𝐼≠0) per cell type, stratified by comparison. **(H)** Stacked bar plot showing the average magnitude of effect sizes per cell type, highlighting compartments with the most substantial transcriptional shifts. **(I)** Directionality of effect size changes. The count of significantly upregulated (positive) and downregulated (negative) features across comparisons. **Figure 6 Alt text:** A nine-panel figure showing Bayesian evaluation of the immune fine compartment. Panels A, B, and C are heatmaps of posterior mean effect sizes across granular immune cell states including CD8-positive exhausted and effector memory T cells, regulatory T cells, CD4-positive Th1 cells, and tumor-associated macrophages for overweight versus normal, obese versus normal, and obese versus overweight comparisons respectively. Panel D is a three-way Venn diagram of shared credible signatures. Panels E and F show ridge plots of pooled cell-type posterior distributions separated and overlaid across comparisons. Panels G, H, and I are stacked bar plots showing credible feature counts, average effect size magnitude, and directionality of changes per cell type respectively.

CD4+ Th1 cells showed credible downregulation of six signatures in both OW vs N (posterior means: −0.264 to −0.288) and OB vs N (posterior means: −0.177 to −0.199), with no credible differences in OB vs OW (Figures 6A, 6B). Suppressed programs include Type I interferon response, IFN-γ response, chemokine production, cytotoxic effector function, PI3K-AKT signaling, and hypoxia response, indicating a broad collapse of Th1 effector capacity. Effect sizes were larger in OW vs N than in OB vs N across all six signatures, and the pooled cell-type posterior mean shifted from −0.269 in OW to −0.183 in OB (Figures 6E, 6F). Th1 suppression is therefore most pronounced at the OW transition and partially attenuates as BMI increases further, suggesting a threshold effect rather than progressive changes.

CD4+ Treg cells showed a distinct two-phase response. In OW vs N, cell signatures were credibly upregulated (posterior means: 0.194 to 0.254; pooled posterior: +0.229), driven by a coordinated elevation of metabolic programming, including lipid metabolism, fatty acid oxidation, mitochondrial function, mTOR signaling, and metabolic regulation along with adipokine and JAK-STAT signaling, and upregulation of Treg identity and TGF-β signaling (Figures 6A, B, E, F). This indicates that OW Tregs are both metabolically primed and transcriptionally reinforced in their regulatory identity. In OB vs N, no feature-level signatures reached credibility, and the pooled posterior declined to near zero (+0.044), indicating attenuation of the OW-established state. In OB vs OW, credible signatures were reversed, comprising exclusively the metabolic and nutrient-sensing programs, including lipid metabolism, mTOR, fatty acid oxidation, mitochondrial function, metabolic regulation, JAK-STAT, adipokine, and lipid peroxidation signaling (Figure 6C). The other signatures also showed reductions, including Treg identity, TGF-β signaling, Hedgehog signaling, and Type I interferon response; however, they did not meet the credibility criteria. This selective metabolic reversal, without credible identity reversal, suggests that OW-established regulatory capacity is not transcriptionally dismantled in obesity.

NK cells showed the most interpretively rich pattern in the compartment. In OW vs N, all signatures were credibly upregulated (posterior means: 0.120-0.146) across metabolic programs, including lipid metabolism, fatty acid oxidation, and mitochondrial function; effector machinery, including degranulation and KIR and NCR receptor signaling; and DNA repair and epigenetic silencing (Figure 6A). Crucially, immunosenescence and NK exhaustion signatures were among the credibly upregulated features, and inhibitory receptor signaling rose in parallel with effector programs, indicating OW NK cells are under sustained pressure rather than in a state of clean activation. In OB vs N, the pooled cell-type posterior shifted from +0.131 to −0.070, and no feature reached credibility, indicating complete dissolution of the OW-activated state in clinical obesity. The OB vs OW comparison showed credible downregulation across all signatures (posterior means: −0.182 to −0.224), with exhaustion, senescence, and immunosenescence signatures among the downregulated features. This pattern indicates a simultaneous collapse of both functional and stress-adaptation programs, rather than classical exhaustion, in which effector function falls while exhaustion markers rise. Because both OW vs N and OB vs N converged in this compartment, the directional trajectory of OW-specific activation followed by collapse in obesity carries stronger evidential weight than the immune coarse NK findings.

In OW vs N, dendritic cells showed credible suppression spanning both activating and tolerogenic programs: phagocytosis, TLR signaling, maturation, costimulation, chemokine production, Type I interferon response, hypoxia response, and tolerogenic programming (Figure 6A). The simultaneous suppression of both axes indicates OW dendritic cells are not shifted toward tolerance but are broadly impaired across all functional programs. In OB vs N, the nature of DC impairment changed qualitatively: programs broadly suppressed in OW were no longer credibly different from N in OB, while interferon signaling deficits deepened, and glycolysis and cell proliferation emerged as new credible deficits (Figure 6B). OW produces generalized DC paralysis; obesity narrows this to a specific and deepening deficit in innate sensing and interferon signaling, accompanied by newly acquired metabolic insufficiency.

TAM showed a pattern of progressive dysfunction that expands in both depth and scope with rising BMI. In OW vs N, signatures were suppressed, reflecting reductions in phagocytic activity, chemokine production, insulin resistance, ECM remodeling capacity, lysosomal function, inflammatory resolution, and the tumor-support program (Figure 6A). As BMI rose into the obese range, these programs deepened further in effect size, and additional signatures reached levels of credibility not observed at OW, including M2 polarization, angiogenic support, complement regulation, iron handling, and oxidative stress defense (Figure 6B). This confirms that the additional programs reaching credibility in OB vs N were not newly suppressed from OW to OB they were already directionally reduced at OW but required the larger BMI contrast to cross the credibility threshold.

Across immune cell lineages, OW correlates with a net elevation of immune programs: naive B cells, NK cells, and Treg cells exhibit directional activation, as do CD8+ T cells, whereas Th1 effectors are inhibited, and dendritic cells and TAMs show broad functional impairment. In OB vs N, plausible alterations are approximately evenly divided between upregulation and downregulation, indicative of incoherence rather than a gradual transition from OW (Figure 6D, 6H). The comparison between OB and OW shows that only credible features align, with lineages stimulated at OW collapsing at OB. This suggests the immune response is a transient, metabolically driven activation, not a persistent state impossible for obese tumors to sustain. The obese microenvironment isn’t just an enlarged overweight or subdued baseline; it’s qualitatively different, with early activation reversed and surveillance restructured across lineages.

### Continuous BMI modeling reveals dose-dependent and previously masked signature dynamics

The continuous BMI model examines how signature activity scales as a dose-dependent function of standardized BMI, complementing the categorical comparisons and identifying programs that respond progressively (Figure 7A-E).

**Figure 7:**
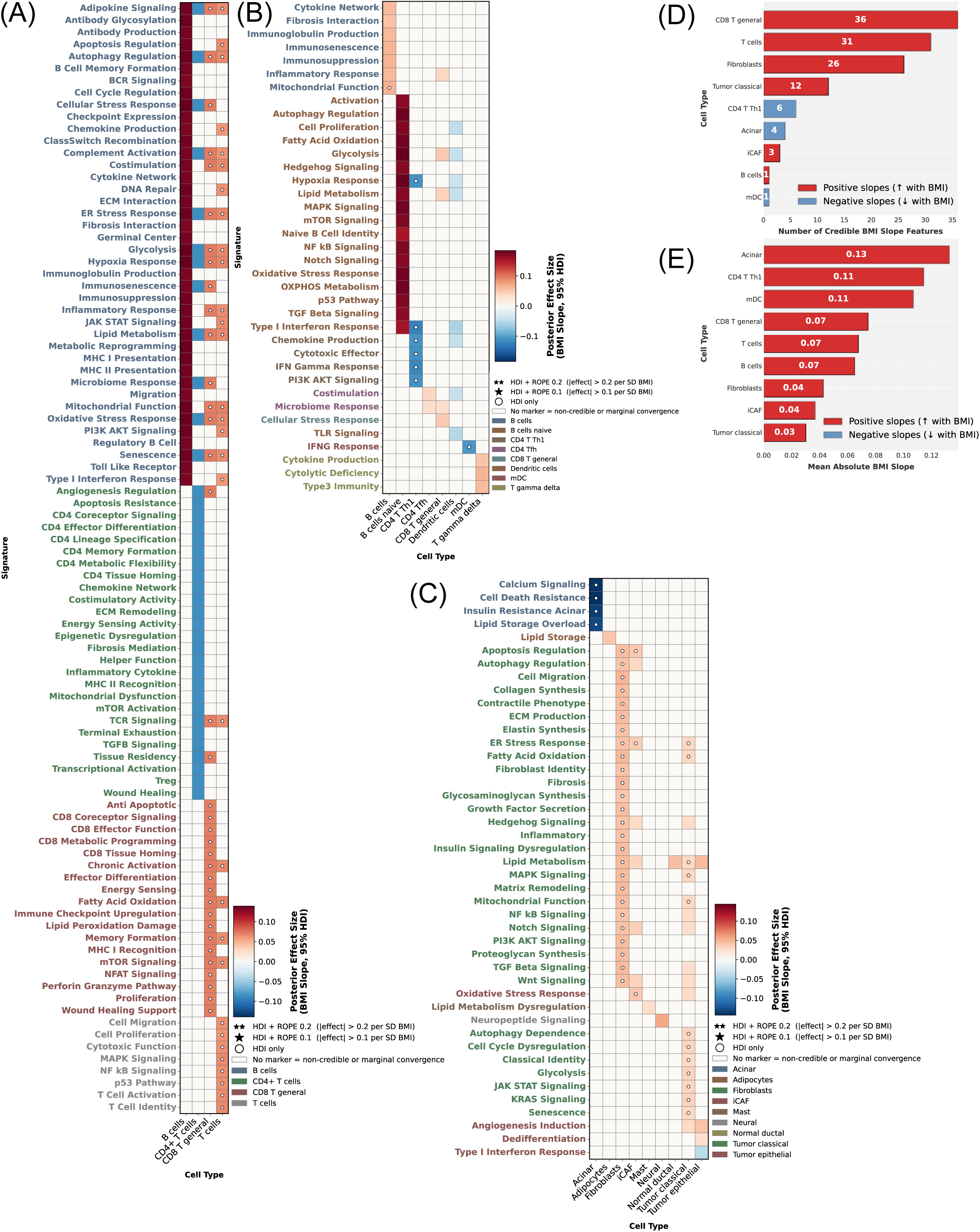
Continuous Bayesian Linear Modeling Reveals BMI-Dependent Dose-Response Trajectories in the PDAC Microenvironment. Heatmap visualization of posterior mean BMI slope coefficients (𝛽_𝐵𝑀𝐼_) for functional gene signatures across **(A)** Non-Immune, **(B)** Immune Coarse, and **(C)** Immune Fine compartments. The color scale represents the rate of change in signature expression per unit increase in BMI (ΔExpression/ΔBMI), identifying linear relationships. Rows represent functional signatures, color-coded by their associated cell type, and columns are cell types. Statistical credibility and practical significance of the slope are indicated by markers: Double Star (★★) denotes strong practical significance (95% HDI excludes 0 and > 95% of the posterior distribution falls outside a ROPE of ±0.02); Single Star (★) denotes moderate practical significance (95% HDI excludes 0 and > 95% of the posterior distribution falls outside a ROPE of ±0.01); White Circle (○) denotes statistical credibility only (95% HDI excludes 0). **(D)** Bar plot showing the number of credible BMI-associated features (HDI ≠ 0) per cell type. **(E)** Bar plot showing the average magnitude of the BMI slope 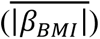, highlighting cell types with the most sensitive transcriptional response to metabolic shifts. **Figure 7 Alt text:** A five-panel figure showing continuous Bayesian linear modeling results. Panels A, B, and C are heatmaps of posterior mean BMI slope coefficients for the non-immune, immune coarse, and immune fine compartments, respectively, with rows representing functional signatures, color-coded by cell type, and statistical credibility indicated by stars and circles. Panel D is a bar plot of credible BMI-associated feature counts per cell type. Panel E is a bar plot of the average BMI slope magnitude per cell type.

In the non-immune compartment, the continuous model largely confirmed the categorical findings (Figure 7C). Fibroblasts showed credible positive BMI slopes across all signatures, and acinar cells showed credible negative slopes across all four signatures, reproducing the categorical suppression pattern. iCAF contributed three credible positive slopes, which include apoptosis regulation, ER stress, and oxidative stress, providing dose-dependent support for the OW-associated activation signal, though narrower in mean effect size than the categorical model.

The most consequential non-immune finding was the emergence of Tumor Classical as a credibly responding population, absent from categorical comparison. Twelve signatures scaled credibly with rising BMI, spanning oncogenic signaling programs (KRAS, JAK-STAT, and MAPK), metabolic reprogramming (glycolysis, fatty acid oxidation, mitochondrial function, and lipid metabolism), and stress and identity programs (ER stress, autophagy dependence, cell cycle dysregulation, senescence, and classical tumor identity). The KRAS signature increase was most pronounced at severe obesity (Supplementary Figure 13E-H), and the gemcitabine resistance signature showed a non-significant positive trend with BMI (Supplementary Figure 13A-D). These findings indicate that the oncogenic and metabolic burden in classical tumor cells accumulates continuously with adiposity, along a gradual, dose-dependent trajectory that the categorical model cannot resolve.

In the immune compartments, the continuous model confirmed the dose-dependent nature of CD8+ T cell activation, with all signatures showing credible positive slopes in the coarse compartment. The broad T cell population showed a comparable pattern with signatures showing credible positive slopes, including metabolic, effector, and stress programs. CD4+ Th1 suppression was also confirmed as dose-dependent, with all suppressed signatures in the categorical model returning credible negative slopes in the continuous fine model, including Type I interferon response, IFN-gamma response, chemokine production, cytotoxic effector function, PI3K-AKT signaling, and hypoxia response. Monocyte-derived dendritic cells contributed a single credible negative slope for IFN-gamma response, consistent with the broader interferon signaling reduction identified in the categorical dendritic cell analysis. These ongoing dynamics show that dose-dependent progressive change and obesity threshold effects occur together across different cell types: activation of CD8+ T cells, suppression of Th1, and increase in tumor oncogenic burden all gradually rise with BMI. Meanwhile, monocyte collapse and the specific activation of stromal and innate immune cells in the OW represent distinct transitions that continuous models cannot further enhance.

### Obesity rewires tumor immune cell–cell communication networks

Cell-cell interaction analysis was conducted using TimiGP, with three immune signature sets (Bindea, Newman LM22, and Zheng) confined to N and OW cohorts, as the OB group lacked sufficient sample size for reliable network inference ^37,38^. TimiGP computes a favorability score for each cell type based on its interaction network, where positive scores indicate anti-tumor functions and favorable prognosis, and negative scores indicate pro-tumor functions and poor prognosis. This scoring reflects the immunological assumption that the balance of pro- and anti-tumor cells dictates whether the tumor immune microenvironment is effective or suppressive, ultimately affecting prognosis.

In N patients, the Bindea signature revealed a T cell-centric interactome, with T cells accounting for the top five interactions and carrying the highest favorability score, followed by DC (Figure 8C, G). T cells maintained active contacts with CD8+ T cells, memory T cells, macrophages, and tumor cells, suggesting a coordinated adaptive immune surveillance in which the tumor remains (Figure 8G and Supplementary Figure 14A, B, C). Supplementary T cell contacts included several immunological compartments, including eosinophils (Enrichment ratio, ER = 2.13), neutrophils (ER = 2.21), helper T cells (ER = 2.23), follicular helper T cells (ER = 2.08), B cells (ER = 2.31), and mast cells (ER = 2.37), further showing the elaborative adaptive-innate crosstalk in N patients (Supplementary figure 14D-G, J, and K). Dendritic cells exhibited contacts with both tumor cells (ER = 2.46) and CD8+ T cells (ER = 2.28). Moreover, T cells interacted with tumor cells, indicating active antigen presentation and antigen-specific actions (Supplementary figures 14H, I, and L), suggesting that the tumor remains “visible” to the adaptive immune system. In OW patients, this architecture collapsed entirely. Tumor cells effectively switched partners, disengaging from adaptive immune interactions and establishing dominant interactions with innate effectors. Tumor-mast cell interactions became prominent, and tumor-eosinophil interactions emerged as top-enriched contacts (Figure 8B, I, J). Neutrophils showed a more pronounced interaction in OW, shifting the interactome towards a pro-inflammatory state. The most prominent interaction was between eosinophils and neutrophils (ER=1.96), characterized by SMPD3/SLC25A37, ABHD2/FPR1, and SMPD3/S100A12 interactions associated with lipid metabolism and inflammatory signaling (Supplementary Figure 15A). The second-ranked interaction between mast cell and neutrophil (ER=1.98) exhibited similar metabolic indicators through MAOB/CD93, PPM1H/FPR1, and MAOB/DYSF interaction pairs (Supplementary Figure 15B). T cell involvement decreased substantially among the top cell-cell interactions, mast cell favorability scores rose to become the dominant positive values in the OW network, reflecting the fundamental shift in immune interaction topology.

**Figure 8:**
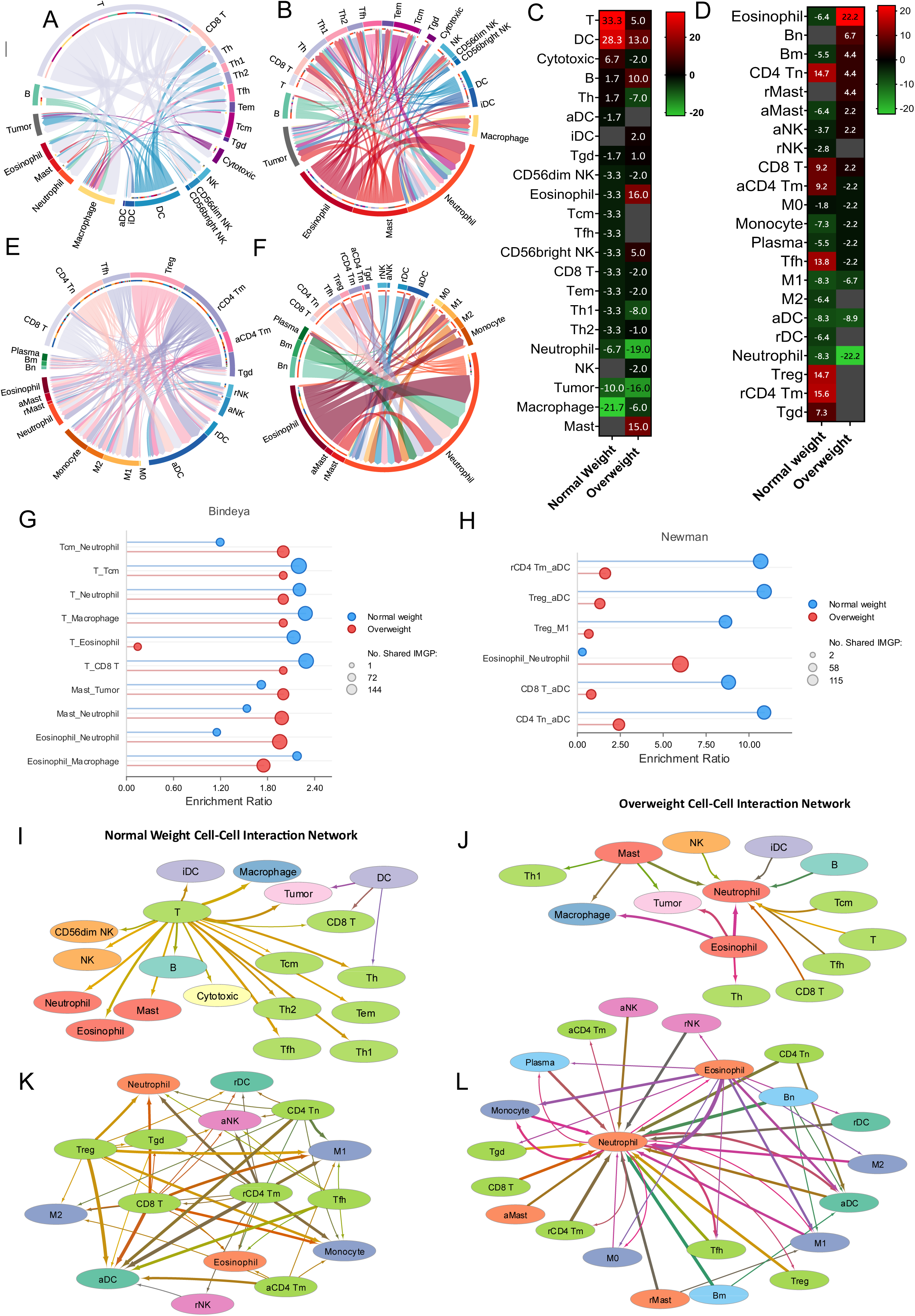
TimiGP analysis using Bindea et al. and Newman et al. LM22 signature reveals shifts from T cell-centric immune cell interactions in normal-weight patients to eosinophil/neutrophil-enriched landscapes in overweight patients, along with prognostic associations. Chord diagrams illustrating the global cell-cell interaction networks in Bindea signature **(A)** Normal Weight and **(B)** Overweight cohorts; in Newmann signature **(E)** Normal Weight and **(F)** Overweight cohorts. Interactions were inferred using TimiGeneNetwork based on the corresponding immune signature, where links represent significant cross-talk between cell types. Heatmap of the TimiGP Favorability Score (TimiFS), depicting the prognostic association of individual immune cell types: Bindea signature **(C),** Newman signature **(D)**. Higher scores indicate a stronger association with favorable outcomes, while lower scores suggest a poorer prognosis. Dot plot displaying the top 5 cell-cell interactions ranked by the group enrichment ratio. Ratios were calculated using Immune Marker Gene Pairs (IMGPs) to highlight interactions that are disproportionately enriched in specific BMI groups: Bindea signature **(G),** Newman signature **(H)**. **(I-L)** Network topology visualizations generated in Cytoscape from TimiGP analysis, representing the spatial organization and connectivity of the immune microenvironment in Normal Weight and Overweight patients; Bindea signature: Normal weight **(I),** overweight **(J)**, Newman signature: Normal weight **(K),** overweight **(L)**. **Figure 8 Alt text:** A twelve-panel figure showing TimiGP cell-cell interaction analysis using Bindea and Newman LM22 immune signatures. Panels A and B are chord diagrams of global immune interaction networks in normal-weight and overweight patients using the Bindea signature. Panels E and F show the same for the Newman signature. Panels C and D are heatmaps of TimiGP favorability scores for the Bindea and Newman signatures, respectively, with warmer colors indicating favorable prognosis associations. Panels G and H are dot plots of the top five cell-cell interactions ranked by enrichment ratio for the Bindea and Newman signatures, respectively. Panels I, J, K, and L are Cytoscape network topology visualizations for normal-weight and overweight patients using the Bindea and Newman signatures, respectively.

The Newman LM22 signature validated these results with enhanced precision (Figure 8D, E, F, H, K I). In N, activated dendritic cells (aDC) emerged as a pivotal interaction hub, with Treg-aDC showing the largest enrichment (ER=10.89), followed by rCD4 Tm-aDC (ER=10.68) and CD4 Tn-aDC (ER=10.88) (Figure 8H). All six aDC contacts prominently expressed the co-stimulatory marker CD86, and five also expressed IL-12B, indicative of Th1 polarization; notable interacting genes were CD3E-CD86, UBASH3A-IL12B, and CD40LG-PDCD1LG2. M1 macrophages, linked to anti-tumor immunity, were also significantly represented (Treg-M1: ER=8.63; rCD4+ Tm-M1: ER=8.05). In the N weight group, almost all T cell lineages have a positive favorability score. In OW patients, the whole aDC-T cell axis was absent, with no aDC-T cell interactions reaching significance (Figure 8G). A single interaction reached significance: eosinophil-neutrophil (ER=6.01), characterized by stress response markers (SMPD3-HSPA6), inflammatory amplification signals (SMPD3-TREM1), and TNF signaling markers (SMPD3-TNFAIP6) (Supplementary Figure 17). The consistent presence of SMPD3 — sphingomyelin phosphodiesterase 3 — across OW interaction profiles mechanistically links sphingolipid-driven lipid metabolic dysregulation to the restructured immune landscape. CD8+ T cell favorability declined from +9.2 to +2.2, and T cell lineage favorability scores flipped from positive in N to near-zero or negative in OW. The network topology shift is visually apparent in the transition from a balanced multi-lineage N network to a neutrophil- and eosinophil-dominated OW structure (Figure 8K, L).

To further understand T cell functional states, we utilized the Zheng signature, which enables more granular analysis of T cell subsets, focusing on activation, exhaustion, and metabolic states. The signature identified 1,560 interactions, of which 58 were significant in N and 7 in OW, confirming the pronounced network collapse previously observed in the Bindea and Newman signatures. In N patients, Interferon-stimulated gene (ISG)-positive T cells accounted for the top interactions (Figure 9D) with the CD8+GZMK+ early Tem interaction with CD4+ISG+ Th (ER=2.13) showed the highest enrichment, demonstrating active type I interferon responses involving interacting gene pairs such as SH2D1A-PLSCR1, CCR4-IRF7, and SH2D1A-STAT (Supplementary Figure 18A). ISG+CD8+ T cells were the main hub, interacting with CD8+TCF7+ Tex, CD8+Tc17, CD4+CCL5+ Tm, and CD4+TNFRSF9+ Treg (ER=2.04) (Figure 9D, Supplementary Figure 18 E-G, and L). These interactions involve ISG markers such as IRF7, STAT1, IFIT1, IFIT3, and CMPK2, suggesting coordinated interferon-driven immune activation in the T cell compartment. Notably, CD8+TCF7+ Tex cells were identified in N interactions (ER = 2.07 for Tex_CD8+ISG+ CD8+ T and Tex_CD4+ISG+ Th). TCF7 (transcription factor 7) is a master regulator of stem-like exhausted T cells that maintain proliferative capacity and respond to checkpoint blockade therapy ^39^. These cells exhibited a progenitor-exhausted phenotype, characterized by interactions involving CCR7/OAS3, CD40LG-SP100, and CCR4-IRF7 gene pairs. Additionally, CD8+GZMK+ early Tem cells were observed in cell-cell interactions, characterized by granzyme K expression and markers indicative of early effector memory differentiation rather than terminal differentiation (Figure 9D and E).

**Figure 9:**
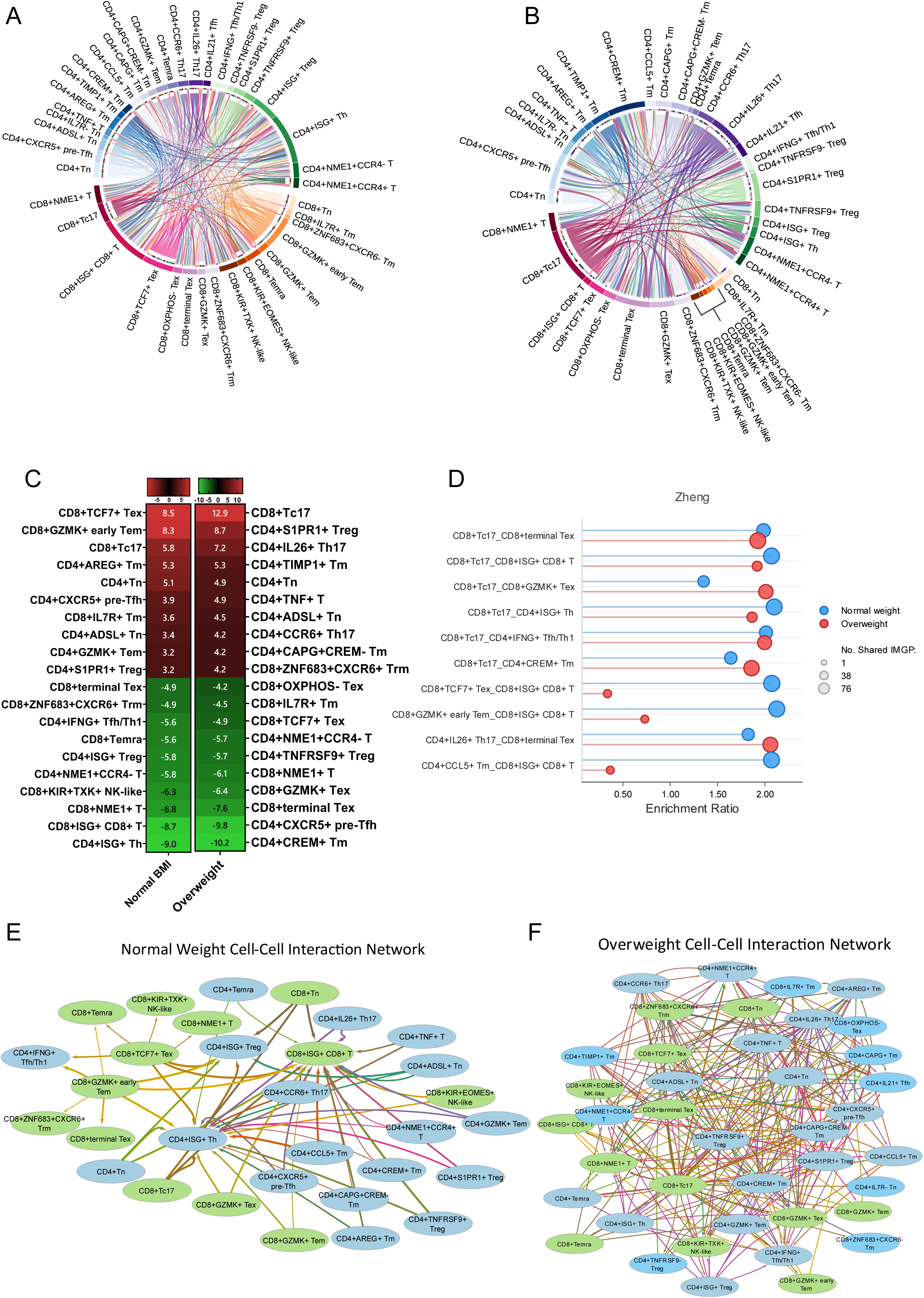
TimiGP analysis using the Zheng et al. signature provides high-resolution characterization of diverse T cell populations in normal-weight and overweight PDAC patients. (A-B) Chord diagrams illustrating the global cell-cell interaction networks in (A) Normal Weight and (B) Overweight cohorts. Interactions were inferred using TimiGeneNetwork based on the Zheng et al. immune signature, where links represent significant cross-talk between cell types. **(C)** Heatmap of the TimiGP Favorability Score (TimiFS), depicting the prognostic association of individual immune cell types. Higher scores indicate a stronger association with favorable outcomes, while lower scores suggest a poorer prognosis. **(D)** Dot plot displaying the top 5 cell-cell interactions ranked by the group enrichment ratio. Ratios were calculated using Immune Marker Gene Pairs (IMGPs) to highlight interactions that are disproportionately enriched in specific BMI groups. **(E-F)** Network topology visualizations generated in Cytoscape from TimiGP analysis, representing the spatial organization and connectivity of the immune microenvironment in **(E)** Normal Weight and **(F)** Overweight patients. **Figure 9 Alt text:** A six-panel figure showing TimiGP cell-cell interaction analysis using the Zheng T cell signature in normal weight and overweight PDAC patients. Panels A and B are chord diagrams of global T cell interaction networks in normal-weight and overweight cohorts, respectively. Panel C is a heatmap of TimiGP favorability scores with warmer colors indicating favorable prognosis associations. Panel D is a dot plot of the top five cell-cell interactions ranked by enrichment ratio. Panels E and F are Cytoscape network topology visualizations for normal-weight and overweight patients, respectively.

In OW patients, ISG+ T cells were absent from significant interactions, either losing their interferon-responsive phenotype or becoming non-prominent in the analysis (Figure 9D). CD8+ Tc17 cells dominated the network, appearing in the top interactions, along with CD4+ IL-26+ Th17 cells, with CD4+IL26+ Th17_CD8+ terminal T cells (ER=2.06) as the primary interaction, characterized by IL-17 receptor signaling and terminal exhaustion transcription factors (TOX2, RBPJ), and CD8+Tc17 × CD8+terminal TEx (ER=1.93) as the second-ranked interaction with comparable marker patterns (Figure 9D). The most consequential finding from the favorability data was the reversal of CD8+TCF7+ Tex from the most favorable cell type in N to unfavorable in OW, directly indicating loss of the checkpoint-responsive progenitor pool at the OW transition. Notably, CD8+ISG+ T cells, which dominated N cell-cell interactions, were absent in OW, either losing their interferon-responsive phenotype or becoming less prominent in the analysis. Instead of CD8+TCF7+ Tex cells, which were found in N interactions, OW had CD8+terminal Tex and CD8+GZMK+ Tex, both of which are later-stage exhausted populations without stem-like features. CD4+CREM+ Tm cells became the most unfavorable cell type in OW, expressing the cyclic AMP response element modulator associated with T cell anergy, while CD4+CXCR5+ pre-Tfh cells, modestly favorable in N, became deeply unfavorable in OW. CD8+OXPHOS-negative Tex cells were additionally prominent (CD8+Tc17 × CD8+OXPHOS-Tex, ER=1.93), indicating mitochondrial failure under lipotoxic stress and forced reliance on glycolysis for any residual effector functions ^40,41^. OW interactions were characterized by IL-17 pathway markers (IL17RE, IL23R, CA2), terminal exhaustion transcription factors (TOX2, RBPJ, ETV1), and glycolytic enzymes (ENO1, GPI, PKM, PFN1), in contrast to the interferon response genes (IRF7, STAT1, IFIT family), TCR signaling molecules (CD40LG, LTB), and early activation markers (SH2D) that defined N interactions (Supplementary Figure 19). CD8+ terminal Tex favorability declined from -4.9 in N to -7.6 in OW, indicating a deepening prognostic penalty for terminally exhausted populations in the OW microenvironment (Figure 9C). The differences between the groups were also evident in the interaction networks of cells within each group (Figure 8I and J).

These three independent signatures demonstrate that overweight status reduces immune interaction complexity and fundamentally restructures the tumor immune microenvironment, replacing coordinated adaptive responses centered on T cells, dendritic cells, and interferon signaling with a neutrophil-dominated inflammatory network driven by sphingolipid metabolic dysregulation, characterized by terminal T cell exhaustion, and marked by the loss of checkpoint-responsive TCF7+ progenitor populations.

### Virtual multiplex immunofluorescence identifies BMI-dependent spatial displacement of stromal-trapped CD8+ T cells in PDAC

Transcriptomic profiling and Bayesian modeling revealed functional changes in the TME, but cannot determine how these changes manifest in tissue space. To examine spatial changes, we applied gigaTIME, a multimodal AI that generates virtual multiplex immunofluorescence images from standard H&E slides, producing images for 9 markers spanning tumor epithelial, immune, and stromal compartments (Supplementary Figure 21). Across all markers, no significant changes in expression levels were found between BMI groups (Supplementary figures 20A and 19B), consistent with the absence of compositional changes in BayesPrism deconvolution and indicating that BMI-associated dysfunction is functional rather than compositional in nature.

Analysis of spatial matrices revealed a distinct pattern. Stromal-trapped CD8+ T cells, defined as CD8+ pixels with centroids falling within Transgelin-positive (Tg+) stromal regions, were located significantly closer to the CK+ tumor boundary (p = 0.033) in OB patients than in N patients (Figure 10, supplementary figure 20C). Continuous BMI analysis confirmed this relationship, with stromal CD8-to-CK+ tumor boundary distance declining progressively with rising adiposity (Supplementary Figure 20D). This geometric compression was accompanied by a reduction in the overall stromal area fraction (Tg+ pixels/total pixels) with increasing BMI (Supplementary Figure 20E), whereas overall Tg+ density showed no significant change (Supplementary Figure 20B). Together, these observations indicate that the Tg+ stromal compartment itself contracted with rising BMI rather than CD8+ T cells actively advancing toward the tumor, a finding consistent with the Bayesian model’s identification of fibroblast ECM remodeling as an early, low-amplitude event established at the OW transition.

**Figure 10:**
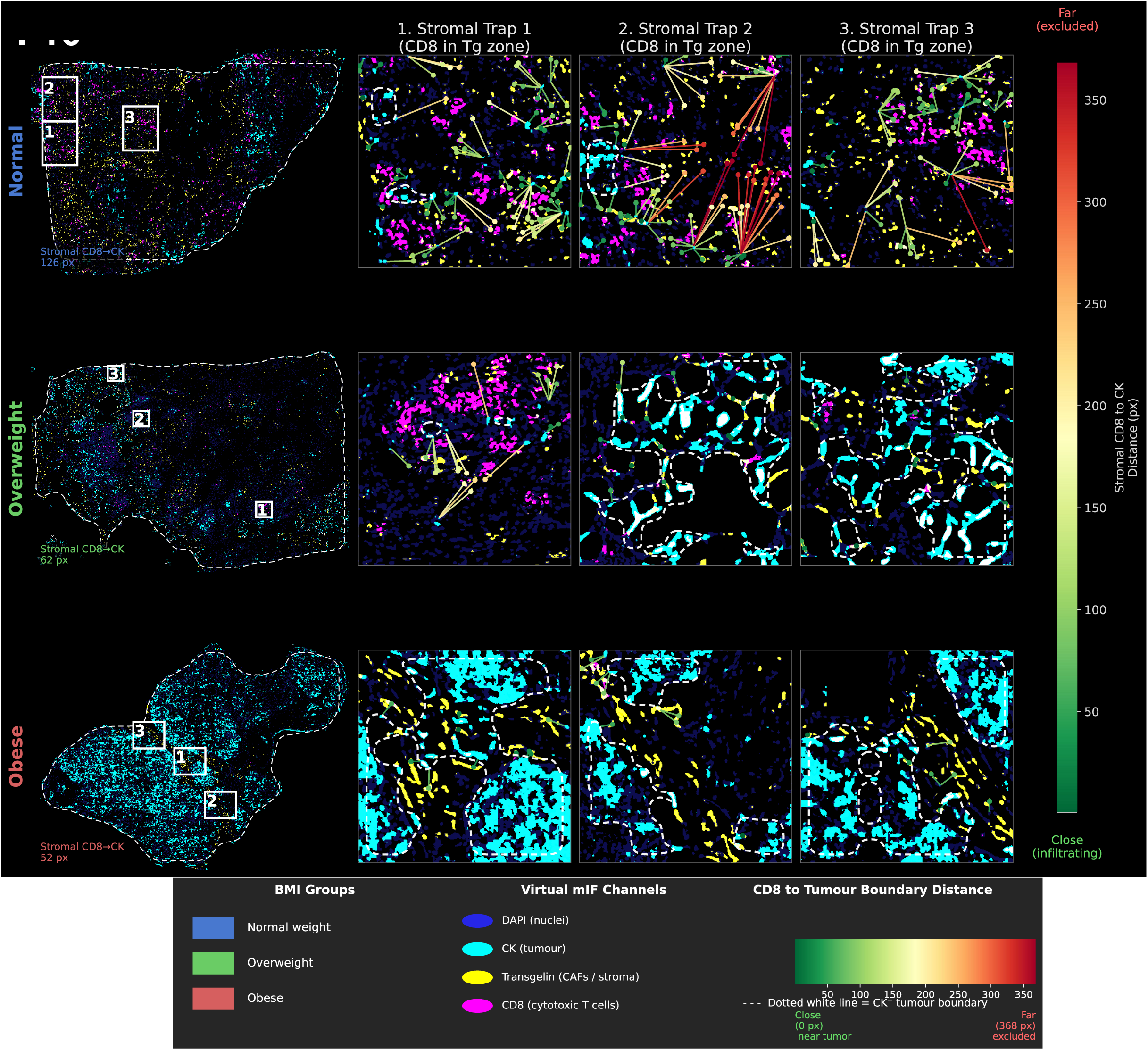
Stromal-trapped CD8+ T cells are positioned closer to the tumor boundary with increasing BMI. Each row represents one BMI group. Left panels show the whole-section composite image with DAPI (blue-violet), CK (cyan), Transgelin (Tg, yellow), and CD8 (magenta), with the tissue boundary indicated by a dashed white outline derived from DAPI. Numbered boxes indicate the three zoomed regions shown to the right of each macro panel. In each zoomed region, lines connect stromal-trapped CD8+ T cell centroids (dots) to the nearest CK+ tumor pixel; line and dot color indicate distance (red, far from tumor boundary; green, close to tumor boundary). The dotted white line in each zoomed panel traces the collective CK+ tumor boundary derived from the CK mask. **Figure 10 Alt text:** A multiplex immunofluorescence image panel organized in three rows, one per BMI group (normal weight, overweight, and obese). In each row, the leftmost panel shows a whole-section composite image with four fluorescent markers: DAPI in blue-violet marking all nuclei, cytokeratin in cyan marking tumor epithelial cells, Transgelin in yellow marking stromal regions, and CD8 in magenta marking cytotoxic T cells. A dashed white outline indicates the tissue boundary. Three numbered boxes in each whole-section panel indicate zoomed regions shown to the right. In each zoomed panel, dots represent stromal-trapped CD8-positive T cell centroids and lines connect each centroid to the nearest cytokeratin-positive tumor pixel. Dot and line color indicate distance from the tumor boundary, with red indicating far and green indicating close. A dotted white line traces the collective cytokeratin-positive tumor boundary. Across the three rows, the stromal-trapped CD8-positive T cell centroids shift progressively closer to the tumor boundary from normal weight to obese patients, indicating geometric compression of immune cells within a contracting stromal compartment with rising body mass index.

Spatial co-localization between CD34+ vasculature and Tg+ fibroblasts increased with BMI (Supplementary figure 20G), suggesting a progressive association of vasculature with the fibroblast compartment at higher BMI. The spatial distance between stromal-trapped CD8+ T cells and the nearest PD-L1+ pixels also increased with BMI (Supplementary figure 19F), indicating that the stromal-confined T cells become more spatially separated from PD-L1 signal as BMI rises. Taken together, transcriptionally defined functional collapse of T cells in obesity is accompanied by physical narrowing of the stromal immune exclusion zone, with residual T cells geometrically compressed closer to the tum or boundary within a shrinking stromal compartment rather than achieving productive infiltration or functional activation.

## DISCUSSION

Obesity is recognized as a modifiable risk factor for PDAC incidence and therapeutic response. However, a comprehensive understanding of the effect of obesity on pancreatic cancer and its cellular manifestation is still largely unclear. Our integrative analysis reveals that BMI does not exert linear, dose-dependent effects on the PDAC microenvironment. Instead, OW status consistently emerges as a critical inflection point at which stromal and immune reprogramming is initiated, with obesity consolidating rather than intensifying these changes across most cell types. An integrated landscape of this remodeling is visualized in Supplementary Figure 22.

At the bulk level, normal-weight patients displayed balanced ECM homeostasis, while OW patients showed broad suppression of structural ECM components, accompanied by CD36 and specific integrin subunits, indicating metabolic coupling of the stromal landscape. OB patients showed collagen organization signatures relative to OW, and the OB vs N bulk comparison showed no ECM enrichment terms, suggesting that OW, rather than OB, represents the point of greatest structural change. The Bayesian model resolved the cellular basis: fibroblast ECM remodeling and a fibrosis signature were credible across both high-BMI comparisons but showed no credible differences between OB and OW, and had minimal effect sizes. This pattern of credible but widespread low-amplitude changes established in OW is a recurring pattern of non-immune reprogramming. Acinar cells showed the strongest practically meaningful effects in the compartment, with suppression of calcium signaling, resistance to cell death, insulin resistance, and lipid storage established at OW and not intensifying further. The observation that stromal remodeling is initiated during early weight gain has direct implications for intervention timing: therapies targeting ECM stiffness or stromal metabolism may be most effective before clinical obesity is established.

The enrichment of primary immunodeficiency pathways with rising BMI reflects dysfunctional immune overactivation rather than conventional immune deficiency. The RAG1/RAG2 stoichiometric discordance in OW patients carries specific prognostic implications: tertiary lymphoid structures are emerging predictors of checkpoint therapy responsiveness in solid tumors, and impaired V(D)J recombination signals compromised TLS maturation at the earliest stage of weight gain ^42–44^. The OW immunological peak, confirmed by humoral and adaptive memory pathway enrichment, anti-apoptotic retention of antigen-experienced lymphocytes, and naive B cell activation reaching a ceiling at OW in the Bayesian model, represents an immune state that is hyperactivated yet functionally ineffective, consistent across bulk and cell-type-resolved analyses. The obesity-specific TAP1 reduction adds a mechanistic layer to antigen presentation failure in OB that is distinct from the broader MHC class I suppression shared by both high-BMI groups: antigen loading capacity itself is compromised in obesity, not merely presentation machinery expression, providing a specific basis for cytotoxic T cell recognition failure that is unlikely to be reversed by interventions targeting downstream signaling^45–47^.

Continuous modeling captures a progressive increase in oncogenic programs that accumulate incrementally with BMI. Although not significant, the gemcitabine resistance signature also showed an increase with BMI, which could partially explain reduced chemotherapy efficacy in OB patients ^8^. In the overweight state, intermediate tumor epithelial cells show upregulation of plasticity and metabolic programs that subsequently reverse as BMI rises into the obese range, indicating closure of an epithelial plasticity window rather than progressive oncogenic intensification. In contrast, the classical tumor subpopulation shows dose-dependent accumulation of oncogenic and metabolic programs with rising BMI in the continuous model, including KRAS signaling, glycolysis, and mitochondrial function, representing a distinct trajectory from the non-linear epithelial plasticity pattern. More notably, dedifferentiation signatures with stem-like markers (SOX2, NANOG, POU5F1, KLF4) were augmented in OW patients while diminishing in OB individuals. This suggests that tumor epithelial cells do not follow a single trajectory but instead adapt through distinct TME changes in obesity-driven PDAC. While specific markers like POU5F1 (OCT4) exhibited a significant positive correlation with BMI, indicating a dose-dependent upregulation, which is correlated with an adverse outcome in gastric cancer ^48^. However, most dedifferentiation markers showed weak trends, whereas NANOG, MYC, and CD44 showed negative correlations (*see the Streamlit app for gene-specific changes*). This mixed pattern at the gene level, combined with an overall signature that declines categorically in obesity, suggests heterogeneous regulation of plasticity programs. Moreover, this pattern likely reflects the closure of an epithelial plasticity window under chronic metabolic and inflammatory stress in obesogenic TME. The dedifferentiation capacity of tumor cells is typically associated with metastatic potential and therapeutic resistance in PDAC ^49^. Chronic metabolic constraints might limit certain forms of plasticity while promoting others, as tumor cells adapt to nutrient-poor conditions by selecting metabolically flexible subpopulations and suppressing less-adaptive states ^49–52^. The protective association for some markers may indicate that the transient plasticity during early weight gain might represent an adaptive response rather than a purely pro-tumor mechanism.

The CD4+ threshold collapse specifically in obesity, without OW involvement, identifies clinical obesity as producing a qualitatively more severe T cell immunological state than OW, despite OW being the primary reprogramming inflection point, a distinction invisible to binary obese/non-obese classifications. The partial attenuation of Th1 in OB relative to OW argues that Th1 suppression reaches a plateau at OW rather than worsening continuously, consistent with the broader cross-compartment pattern of OW-established states that do not further intensify in clinical obesity. The progressive CD8+ T cell activation trajectory confirmed by continuous BMI modeling scaling with every unit of adiposity rather than crossing a discrete threshold indicates that there is no safe BMI increment above normal weight at which CD8+ functional compromise stabilizes, consistent with the dose-dependent cytotoxic, metabolically stressed, and exhausted trajectory reported in obesity-driven immune remodeling ^53^. Spatially, the structural confinement of residual CD8+ T cells within a shrinking stromal compartment, with increasing separation from PD-L1+ pixels, indicates that even transcriptionally active CD8+ T cells in obesity are geometrically precluded from productive tumor engagement and from accessing checkpoint signals, a spatial barrier that transcriptional analysis alone cannot detect.

NK cell biphasic behavior, with exhaustion programs co-upregulated alongside activation at OW, indicates that functional collapse in obesity is programmed during the overweight transition rather than triggered by it. The trajectory toward dysfunction is established before the cells fail ^54^. That broad B-cell stress and metabolic activation in obesity reflect the energetic cost of sustaining chronic inflammatory demand rather than productive immune function, which explains how immune activity can be elevated while immune effectiveness declines 36. The monocyte global transcriptional shutdown, confirmed as a pure threshold effect by the continuous model, indicates that obese PDAC monocytes have entirely lost adaptive capacity, not merely redirected it, leaving the innate myeloid compartment unable to respond to microenvironmental stress. These findings collectively indicate that the window for immune intervention is established at OW and narrows rapidly.

The Treg metabolic collapse in obesity without credible identity reversal is therapeutically consequential: if Tregs retain immunosuppressive transcriptional identity while losing metabolic fitness, checkpoint therapy responses may remain suppressed even when metabolic interventions partially restore immune fitness. The TAM dysfunction programs that become statistically credible in OB vs N, but show no credible OB vs OW differences, indicate that these programs were directionally impaired from OW onward and cross statistical detectability only in the larger BMI contrast; the obese TAM is not progressively more dysfunctional than the OW TAM; its dysfunction is more statistically certain. The qualitative OW-to-OB shift in DC impairment from generalized paralysis to a specific deepening of interferon signaling deficits suggests that type I interferon-based therapeutic approaches may face a more specific and potentially addressable deficit in obese patients than in OW patients.

In N patients, the T cell-aDC-centric interactome anchored by ISG+ and TCF7+ stem-like exhausted T cell interactions reflects an immune surveillance system retaining checkpoint blockade responsiveness ^55,56^. In OW, the CD8+ T cell prognostic privilege is substantially weakened^57^, and the interactome shifts to eosinophil-neutrophil dominance driven by SMPD3-mediated sphingolipid dysregulation, linking lipid metabolic stress to inflammatory network restructuring and desmoplasia elevation ^58^. This mechanistic connection suggests that lipid-targeting interventions may confer benefits in immune communication beyond their direct metabolic effects.

At fine resolution, OW patients showed the dominance of type 17 immunity and terminal exhaustion. CD8+ ISG+ cells, dominant in N networks, were reduced in the interactome of the OW group. Instead of CD8+ TCF7+ stem-like exhausted cells, the OW microenvironment displayed CD8+ terminal-exhausted populations lacking stem-like features. In PD1 checkpoint therapy, TCF+ T cells can be revitalized to generate tumor-specific intratumoral CD8+ T cells (59,60). A reduction in TCF7+ interaction in the OW interactome suggests a lack of PD1 checkpoint responses ^60^. Most critically, CD8+OXPHOS-negative exhausted cell interactions appeared, indicating a fundamental metabolic failure. In the lipotoxic environment of obesity, these metabolically stressed T cells lack mitochondrial respiration and are unable to produce sufficient ATP for effector functions, forcing a reliance on glycolysis for effector functions ^40,41^.

This restructuring indicates that obesity not only alters immune cell states but also rewires the cellular communication topology in PDAC. The shift from coordinated adaptive immunity centered on dendritic cell-T cell interactions and interferon signaling to neutrophil-dominated networks represents a qualitative organizational change. Despite widespread activation signatures in individual cell types, the loss of coordinated interactions indicates functional isolation. Immune cells may remain transcriptionally active yet unable to engage in the coordinated communication required for effective anti-tumor immunity. This dissociation between activation and coordination may explain why obesity-associated inflammation does not translate to enhanced tumor control ^61,62^.

## CONCLUSION

In summary, our multi-resolution analysis demonstrates that obesity reshapes the PDAC microenvironment through non-linear, cell-type-specific mechanisms that operate at transcriptional, cellular, and cell network levels. Overweight status consistently emerges as a critical inflection point where stromal and immune reprogramming occurs. Non-immune compartments undergo coordinated low-amplitude remodeling that stabilizes or reverses in obesity rather than intensifying linearly. Tumor epithelial cells exhibit dynamic plasticity windows that close with increasing BMI, while simultaneously acquiring oncogenic and therapy-resistance programs. Immune compartments display heterogeneous, lineage-specific trajectories across BMI strata. Immune activation does not translate to effective anti-tumor immunity, as obesity rewires coordinated adaptive networks into fragmented, inflammation-dominant ones. The identification of Overweight as a dominant transition state suggests that early metabolic interventions may prevent subsequent functional collapse. The emergence of BMI-associated chemotherapy resistance signatures indicates potential clinical utility for BMI-aware treatment stratification.

Several limitations warrant consideration. The obese cohort (n=18) is the smallest group, necessitating validation in larger balanced cohorts. The obese group could not be included in the TimiGP network inference. The gigaTIME analysis is subject to inherent uncertainty due to the absence of ground-truth mIF for CPTAC patients. Analysis is restricted to transcriptional signatures and cannot assess protein-level changes or post-translational modifications. BMI is an imperfect proxy for adiposity, not accounting for body composition or metabolic health status^24,63,64^. Refined classifications, such as metabolically healthy versus unhealthy obesity, may reveal heterogeneity that our model does not capture. Future studies should employ longitudinal cohorts to track microenvironmental changes during weight interventions, larger, balanced cohorts for interaction modeling, and functional validation of chemotherapy resistance and exhaustion signatures in preclinical models. Integration of proteomics and metabolomics will be essential to connect transcriptional programs to functional cellular behavior in the PDAC microenvironment.

## METHODS

### Patient Cohort and Differential Expression Analysis

Transcriptomic data from patients with pancreatic ductal adenocarcinoma (PDAC) were obtained from the Clinical Proteomic Tumor Analysis Consortium (CPTAC-3) (𝑛 = 140) ^65^. Patients were divided into three groups based on World Health Organization BMI classifications: Normal weight (𝑁, 18.5 − 24.9 𝑘𝑔/𝑚^2^, 𝑛 = 51), Overweight (𝑂𝑊, 25.0 − 29.9 𝑘𝑔/𝑚^2^, 𝑛 = 58), and Obese (𝑂𝐵, ≥ 30.0 𝑘𝑔/𝑚^2^, 𝑛 = 18). Raw RNA-seq STAR count matrices were processed using DESeq2 for differential expression analysis (DEA). Functional enrichment was done using the clusterProfiler and Gene Set Enrichment Analysis (GSEA) tools across Gene Ontology (GO), KEGG, and Reactome databases. To complement group-wise comparisons, we performed a continuous correlation analysis of BMI and gene expression using the LinkedOmics platform ^66^. Detailed quality control and normalization procedures are described in the Supplementary Methods.

### Construction of Single-Cell References and Deconvolution

To achieve granularity in the bulk transcriptomic data, we constructed two dataset-specific single-cell RNA sequencing (scRNA-seq) references from publicly available GEO datasets (GEO: GSE242230 and GSE235452) ^67,68^.

#### Tumor and Stroma

Malignant epithelial cells were distinguished from normal ductal cells using *CopyKat* to infer chromosomal copy number variations (CNV) ^69^. Malignant cells were further subtyped into *Basal-like*, Classical, or *tumor epithelial* phenotypes based on established molecular signatures. Stromal populations (fibroblasts, endothelial cells) were annotated using module scoring based on markers. For detailed procedures, see Supplementary Methods.

#### Immune Landscape

We used an immune cell annotation strategy by deriving consensus from readily available lymphocyte annotation databases. CD45+ cells were annotated using *SingleR* ^70^ against the Monaco, DICE, Blueprint, and Novershtern, followed by fine-grained subtyping to identify specific states, including exhausted CD8+ T cells and Tumor-Associated Macrophages (TAMs). TAMs were distinguished from circulating monocytes using a multi-marker scoring of lineage markers and the exclusion of dendritic/neutrophil markers. Bulk RNA-seq data were deconvoluted using *BayesPrism*, a method designed to infer cell type fractions and gene expression profiles from bulk RNA-seq data. For detailed procedures, see Supplementary Methods.

### Immune Dysfunction Signatures and Feature Selection

We curated a comprehensive database of 2,143 immune dysfunction signatures across 65 cell types, capturing functional states such as T-cell exhaustion, metabolic reprogramming, and senescence. LLM-assisted signature refinement using the Google Gemini API (gemini-2.5-flash) was performed as described in the Supplementary Methods. Signature scores were computed using a standardized Z-score approach with winsorization and minimum gene thresholds. To identify BMI-associated cellular features within this high-dimensional dataset, we utilized *Stabl*, a machine learning framework ^71^. Stabl was applied in two modes: a categorical mode (logistic regression) to discriminate between BMI groups, and a continuous mode (linear regression) to identify linearly associated BMI signatures. For detailed procedures, see Supplementary Methods.

### Bayesian Hierarchical Modeling

To quantify the magnitude and uncertainty of BMI-associated cellular changes, we developed a Bayesian hierarchical modeling framework. We implemented two distinct models:

1. **Categorical Model:** Captures non-linear associations across N, OW, and OB groups.
2. **Continuous Model:** Estimates the linear slope of signature changes per standardized unit of BMI.

The models used zero-centered normal priors for effect sizes and half-normal priors for feature-level variability to prevent overfitting; full prior specifications per compartment are provided in the Supplementary Methods. Posterior distributions were approximated using the No-U-Turn Sampler (NUTS). A change in signature is statistically credible if the 95% HDI of the posterior distribution strictly excludes zero and is annotated with “ ∘ ”. Since each patient contributes multiple signature effects, we added a patient-level random intercept to both models using non-centered parameterization. This separates within-patient correlation from BMI-associated effects and prevents pseudoreplication from inflating posterior certainty. To assess the practical significance, we defined a ROPE interval on the standardized scale. A signature is classified as practically significant only if the entire 95% HDI excluded this ROPE interval and was annotated with “★”. For detailed procedures, see Supplementary Methods.

### Cell-Cell Communication and Interactome Analysis

Cell-cell interactions were inferred using the TimiGP framework ^37^. We analyzed cell-cell interaction networks in the N and OW cohorts using immune signatures derived from established cancer immune interaction datasets (Bindea, Newman LM22, and Zheng et al.). The prognostic relevance was assessed using Cox proportional hazards regression. The top 5% of gene pairs ranked by raw p-value (p < 0.01) were selected, and enriched cell-cell interactions were identified using Benjamini-Hochberg adjusted p < 0.05. For detailed procedures, see Supplementary Methods.

### Virtual multiplex immunofluorescence using gigaTIME

To evaluate the physical manifestations of increasing BMI, we used gigaTIME, a NestedUNet cross-translator model trained on a pan-cancer dataset (n=14,256; 24 cancer types), including pancreatic cancer, to generate virtual mIFs from H&E whole-slide images ^26^. Whole slide images of the CPTAC-PDA cohort were obtained from the NCI Imaging Data Commons (IDC) using the *idc-index (V23)* Python package. Full resolution series from multiple DICOM series corresponding to each patient we identified and used for downstream analysis. To identify tissue-containing regions, the whole slide was scanned with non-overlapping 512×512-pixel tiles. Tiles with mean pixel intensities below 220 were retained as tissue. Disconnected tissue fragments were identified by connected component analysis on the resulting tiles to build a coordinate map. Tiles below the minimum size (≥50 tiles) and tissue density (≥30) thresholds were excluded from downstream analysis. Among the selected sections, the best tumor-representative sections were identified by estimating cytokeratin (CK) density using gigaTIME on 30 randomly selected tiles per section. Sections below the CK threshold were discarded for further analysis. A full virtual mIF inference was then performed on the selected section using a 256×256-pixel window with a 128-pixel stride (50% overlap) to generate virtual mIF images for each selected image. Spatial metrics were computed from these canvases using the mask produced for each marker, and BMI association was assessed using Spearman and Pearson correlation; categorical group comparison used Kruskal-Wallis with Benjamini-Hochberg correction. For detailed procedures, see Supplementary Methods.

## Data Availability

The data that support the findings of this study are available within the paper and its Supplementary Information files. The interactive model explorer, TimiGP interactome analysis, and custom signature database can be accessed via our web app at https://obese-pdac-model.streamlit.app/ and also archived at Zenodo [https://doi.org/10.5281/zenodo.19386459]^72^. Data used to construct the scRNA reference for deconvolution are publicly available in GEO under GSE242230 and GSE235452. CPTAC data is accessible through TCGAbiolinks at TCGA.

## Code availability

All custom computer code used for this paper, including signature curation, single-cell reference construction, Stabl ML, deconvolution, Bayesian models, TimiGP, and gigaTIME, is available in the GitHub code repository [https://github.com/arunviswanathan91/obese-model] and archived at Zenodo [https://doi.org/10.5281/zenodo.19386461].

## Author contributions

**AV:** Planning, Coding, Signature Building, Model building, Analysis, and Writing; **JS:** Conceptualized the machine learning pipeline, led code review and correction, GitHub repository architecture, and maintenance; **KBH**: Conceived and supervised the study, secured funding, and critically reviewed and revised the manuscript. All authors read and approved the final version of the manuscript.

## Acknowledgments

Not applicable

## Abbreviations

PDAC: Pancreatic ductal adenocarcinoma
TME: Tumor microenvironment
ECM: Extracellular matrix
BMI: Body mass index
N: Normal weight
OW: Overweight
OB: Obese
CNV: Copy number variation
TLS: Tertiary lymphoid structures
TAM: Tumor-associated macrophage
CAF: Cancer-associated fibroblast
myCAF: Myofibroblastic cancer-associated fibroblast
iCAF: Inflammatory cancer-associated fibroblast
apCAF: Antigen-presenting cancer-associated fibroblast
NK: Natural killer cells
DC: Dendritic cell
aDC: Activated dendritic cell
Treg: Regulatory T cells
Th1: T helper 1 cells
Tfh: Follicular helper T cells
Tex: Exhausted T cells
Tem: Effector memory T cells
Tc17: Type 17 CD8-positive T cells
MAIT: Mucosal-associated invariant T cells
TCR: T cell receptor
MHC: Major histocompatibility complex
TAP1: Transporter associated with antigen processing
PD-1: Programmed cell death protein 1
PD-L1: Programmed death-ligand 1
CTLA-4: Cytotoxic T-lymphocyte-associated protein 4
KIR: Killer immunoglobulin-like receptor
NCR: Natural cytotoxicity receptor
ISG: Interferon-stimulated gene
SMPD3: Sphingomyelin phosphodiesterase 3
TCF7: Transcription factor 7
OXPHOS: Oxidative phosphorylation
RAG1/RAG2: Recombination activating gene 1 and 2
AIRE: Autoimmune regulator
ZAP70: Zeta-chain-associated protein kinase 70
ITK: IL-2-inducible T cell kinase
PAK: p21-activated kinase
AID: Activation-induced cytidine deaminase
BAFFR: B cell activating factor receptor
TACI: Transmembrane activator and calcium modulator and cyclophilin ligand interactor
IFN-γ: Interferon gamma
TNFα: Tumor necrosis factor alpha
TGF-β: Transforming growth factor beta
GM-CSF: Granulocyte-macrophage colony-stimulating factor
IL: Interleukin
KRAS: Kirsten rat sarcoma viral proto-oncogene
JAK-STAT: Janus kinase-signal transducer and activator of transcription
MAPK: Mitogen-activated protein kinase
mTOR: Mechanistic target of rapamycin
NF-κB: Nuclear factor kappa B
PI3K-AKT: Phosphoinositide 3-kinase-AKT
CREM: Cyclic AMP response element modulator
scRNA-seq: Single-cell RNA sequencing
ssGSEA: Single-sample gene set enrichment analysis
GSEA: Gene Set Enrichment Analysis
DEA: Differential expression analysis
VST: Variance stabilizing transformation
RSEM: RNA-Seq by expectation-maximization
UMI: Unique molecular identifier
HDI: Highest density interval
ROPE: Region of practical equivalence
NUTS: No-U-Turn Sampler
ESS: Effective sample size
E-BFMI: Expected Bayesian fraction of missing information
HMC: Hamiltonian Monte Carlo
SNR: Signal-to-noise ratio
FDR: False discovery rate
HR: Hazard ratio
LLM: Large language model
STABL: Stability selection algorithm
LASSO: Least absolute shrinkage and selection operator
TimiFS: TimiGP favorability score
ER: Enrichment ratio
IMGP: Immune marker gene pair
WSI: Whole-slide image
DICOM: Digital imaging and communications in medicine
IDC: NCI Imaging Data Commons
mIF: Multiplex immunofluorescence
CPTAC: Clinical Proteomic Tumor Analysis Consortium
GEO: Gene Expression Omnibus
GO: Gene Ontology
KEGG: Kyoto Encyclopedia of Genes and Genomes
WHO: World Health Organization
CK: Cytokeratin
Tg: Transgelin
DAPI: 4’,6-diamidino-2-phenylindole

