## Supplementary Figures for "Overweight status drives early tumor microenvironment reprogramming in pancreatic ductal adenocarcinoma: a cell-type-resolved Bayesian hierarchical modeling and interactome analysis"

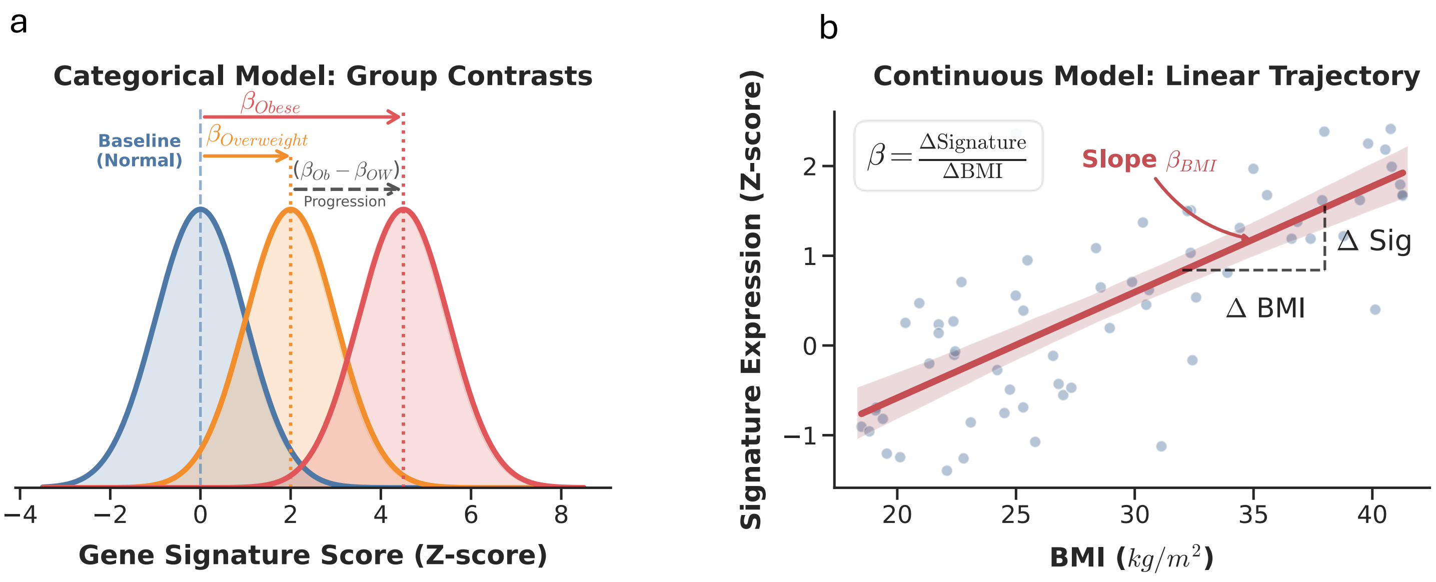


Supplementary Figure 1: Schematic representation of Bayesian hierarchical modeling strategies. a Structure of the Categorical Model. Posterior distributions represent the estimated population-level mean expression (Z-score) for Normal (baseline; blue), Overweight (orange), and Obese (red) cohorts. Arrows indicate the estimated regression coefficients ($\beta$), which represent the magnitude of the effect relative to the Normal baseline. The dashed grey arrow denotes the calculated contrast ($\beta_{Obese} - \beta_{Overweight}$), from overweight to obesity. b Structure of the Continuous Model. A linear regression framework models the trajectory of gene signature expression as a function of BMI ($kg/m^2$). The slope coefficient ($\beta_{BMI}$) quantifies the rate of change in signature expression per unit increase in BMI (ΔSignature/ΔBMI), serving as a metric for the strength and directionality of the continuous association. Shaded regions in both panels represent the uncertainty (95% High Density Interval) associated with the parameter estimates.

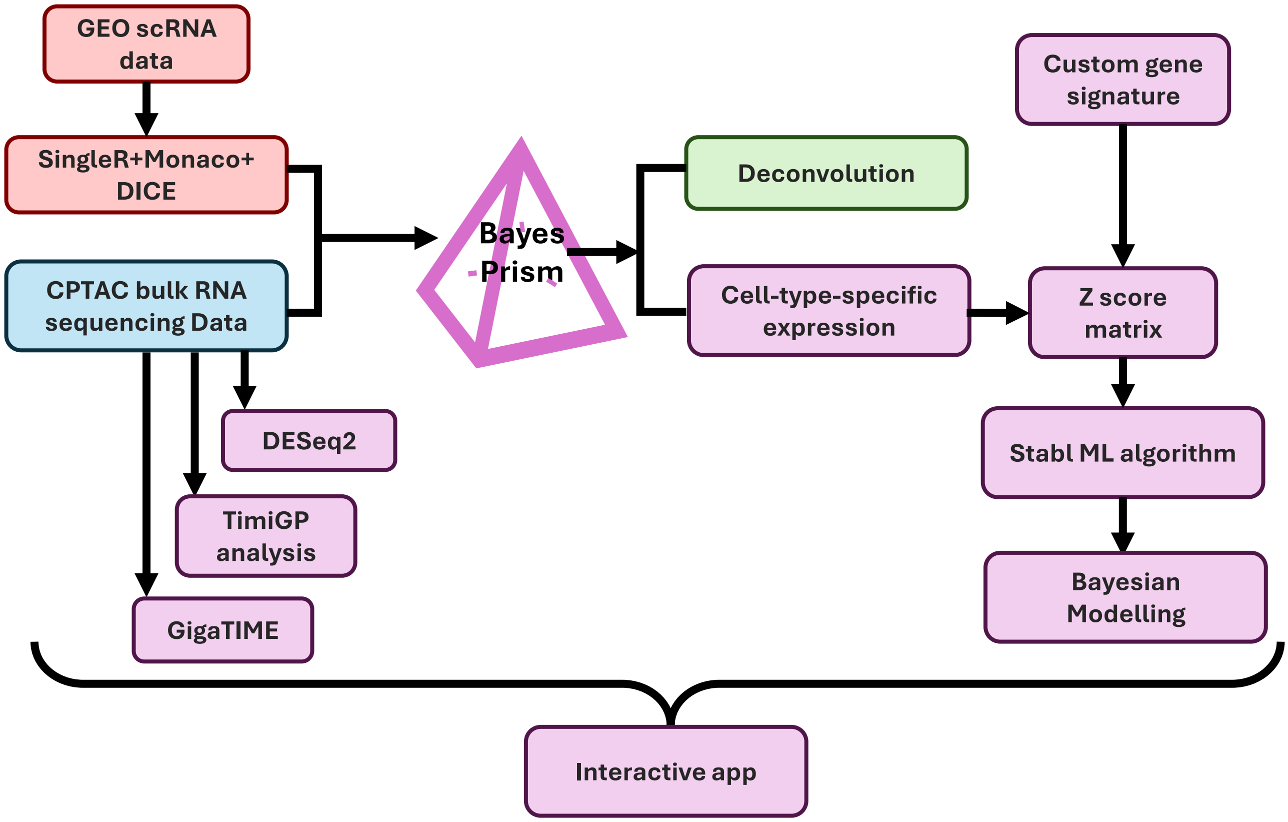


Supplementary Figure 2: Computational Framework for BMI-Stratified PDAC Microenvironmental Analysis: This schematic illustrates the multi-resolution analytical workflow used to dissect the impact of body mass index (BMI) on the pancreatic ductal adenocarcinoma (PDAC) tumor microenvironment.

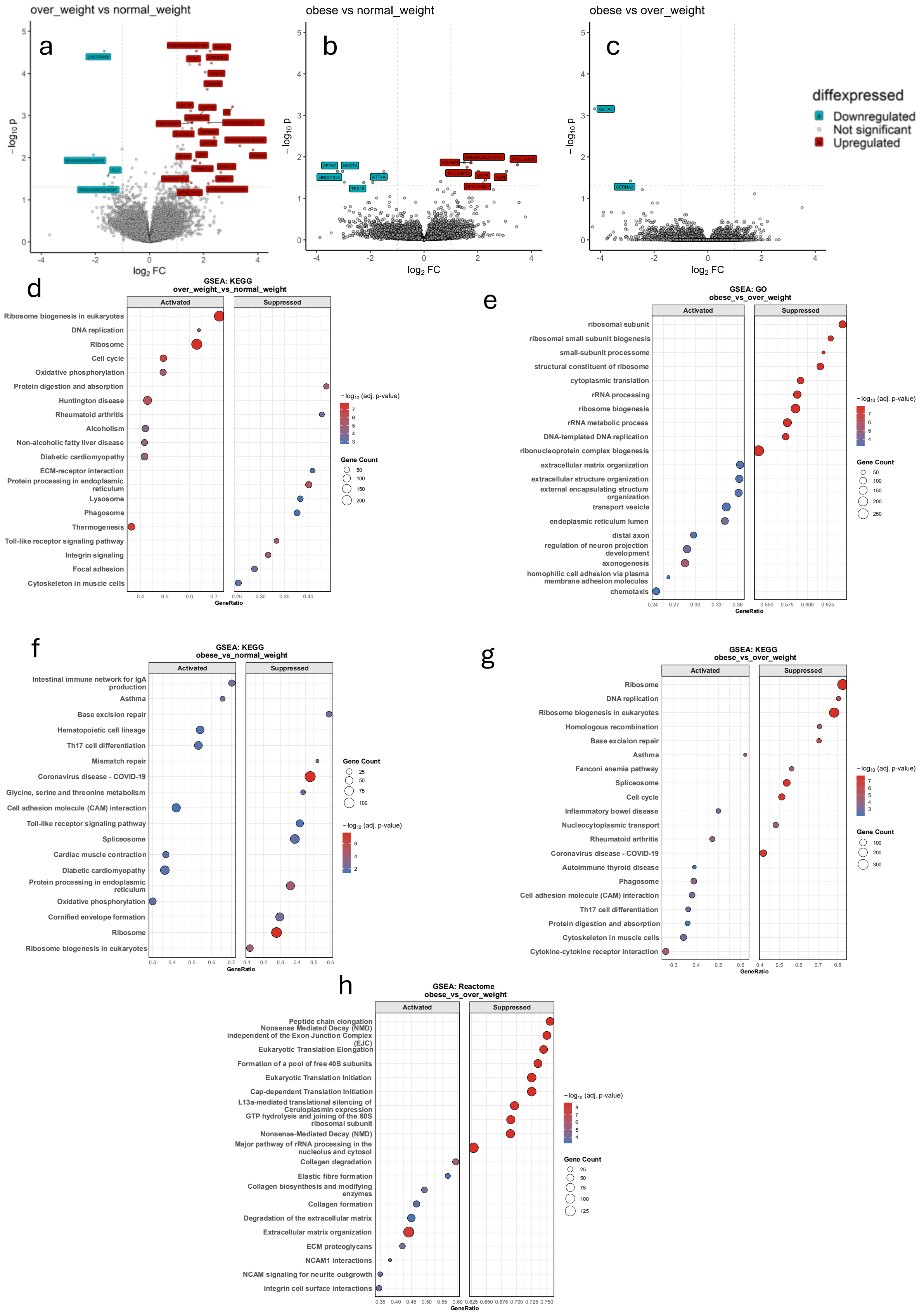


Supplementary Figure 3: Differentially expressed genes in the BMI group comparison using DESeq2. Overweight vs Normal weight a, Obese vs Normal weight b, and Obese vs Overweight c. Only a few significant changes in gene expression with the majority of changes in the overweight vs normal weight comparison. p.adj. ≤ 0.05, log(FC) ≤ 1.5. (D-H) GSEA bubble plots: GO bubble plots showing pathways differentially enriched in KEGG between Overweight vs. Normal weight d, obese vs Overweight g, and Obese vs. Normal weight f patients. GO analysis comparing obese vs overweight e and Reactome analysis obese vs. overweight h groups. Bubble size reflects the number of genes contributing to each term, and color indicates adjusted p-values. Note: In all GSEA plots, *Activated* indicates terms enriched in overweight/obese; *Suppressed* indicates terms enriched in normal weight patients.

### ECM-receptor Interaction Pathway (HSA04512)

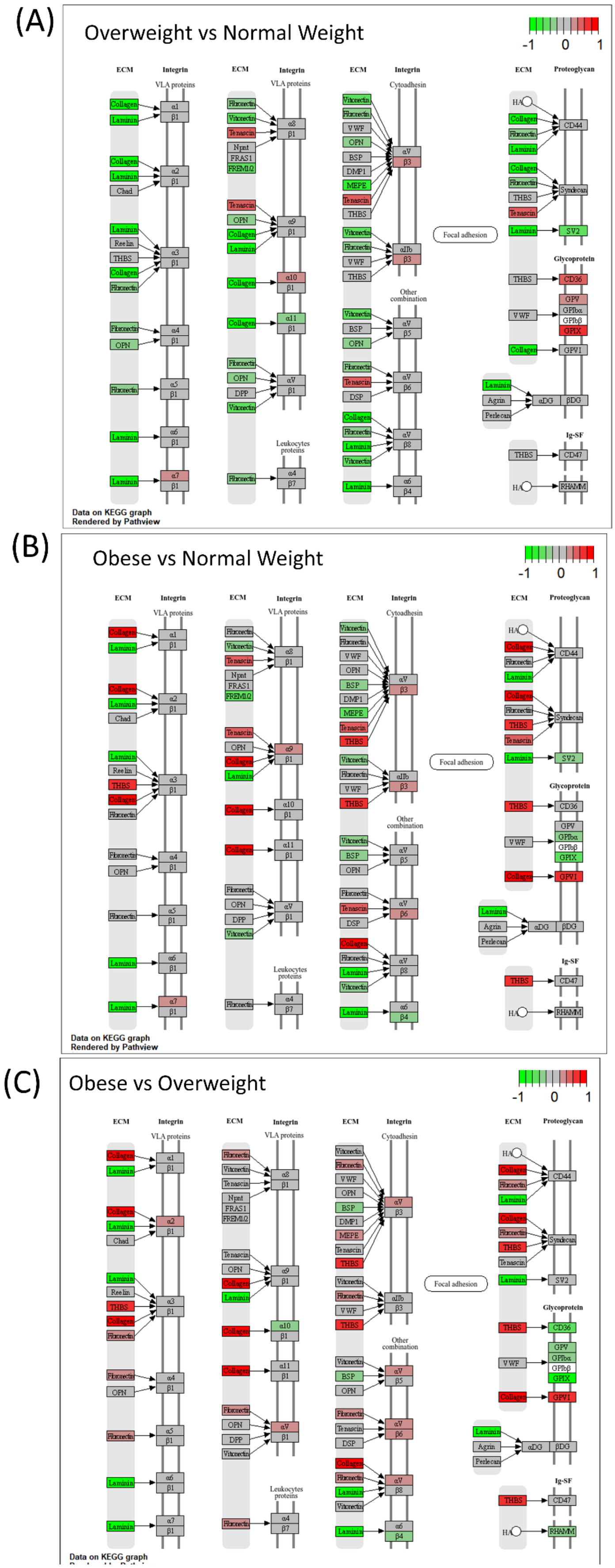


**Supplementary Figure 4:** ECM-receptor interaction pathway (HSA04512) expression in pancreatic cancer across BMI categories. KEGG pathview analysis showing differentially regulated genes in the extracellular matrix-receptor interaction pathway between overweight vs. normal weight a and obese vs normal weight b. And obese vs. Overweight c pancreatic cancer patients. Color scale indicates gene expression levels (Green: downregulation [-1]; red: upregulation [+1] in the comparison group relative to the reference group).

### Cytokine-cytokine receptor interaction pathway (HSA04060)

(A) Overweight vs Normal Weight

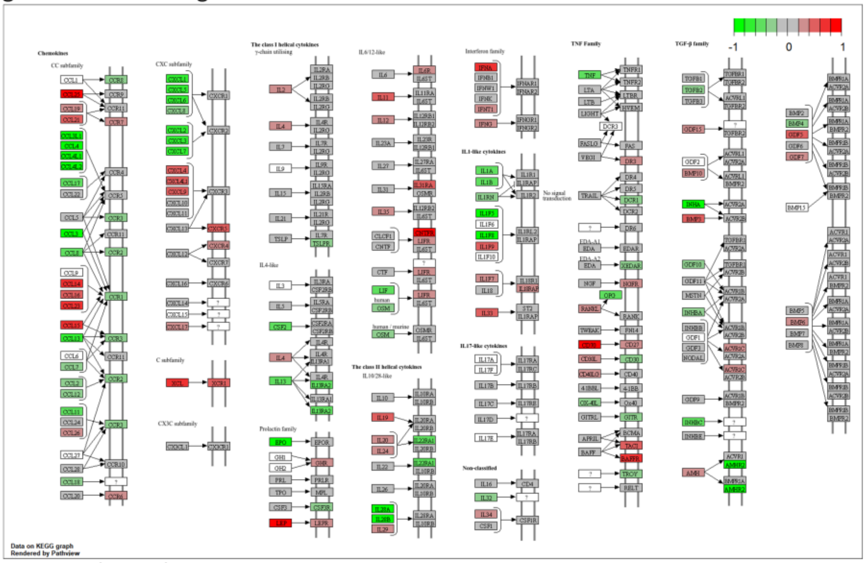


(B) Obese vs Normal Weight

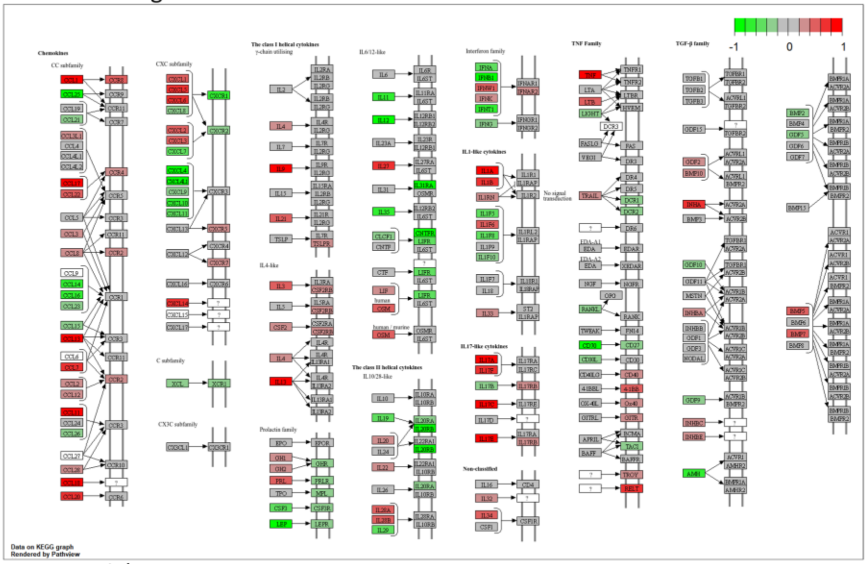


(C) Obese vs Overweight

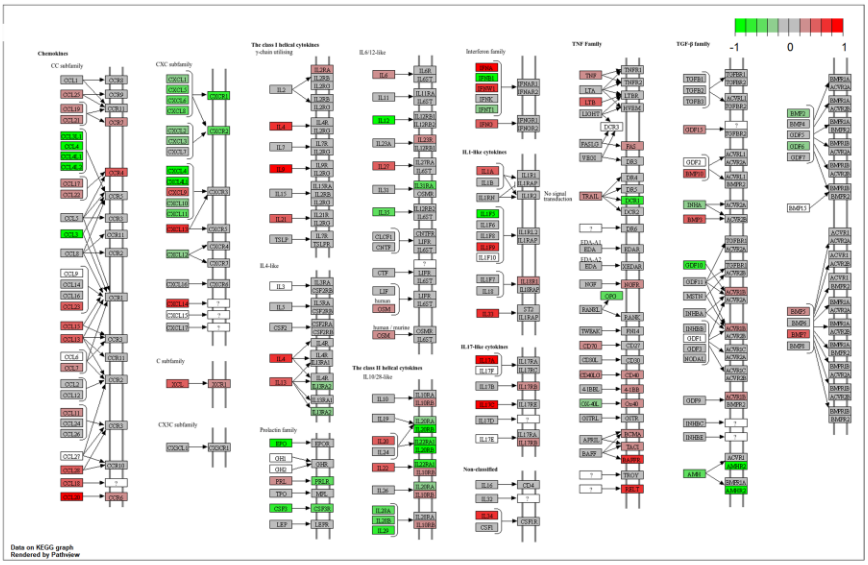


**Supplementary** Figure 5: Cytokine-cytokine receptor interaction pathway (HSA04060) comparison across BMI groups in pancreatic cancer. KEGG pathview analysis showing differentially regulated genes in the extracellular matrix-receptor interaction pathway between overweight vs. normal weight a and obese vs normal weight b. And obese vs. Overweight c pancreatic cancer patients. Color scale indicates gene expression levels (Green: downregulation [-1]; red: upregulation [+1] in the comparison group relative to the reference group).

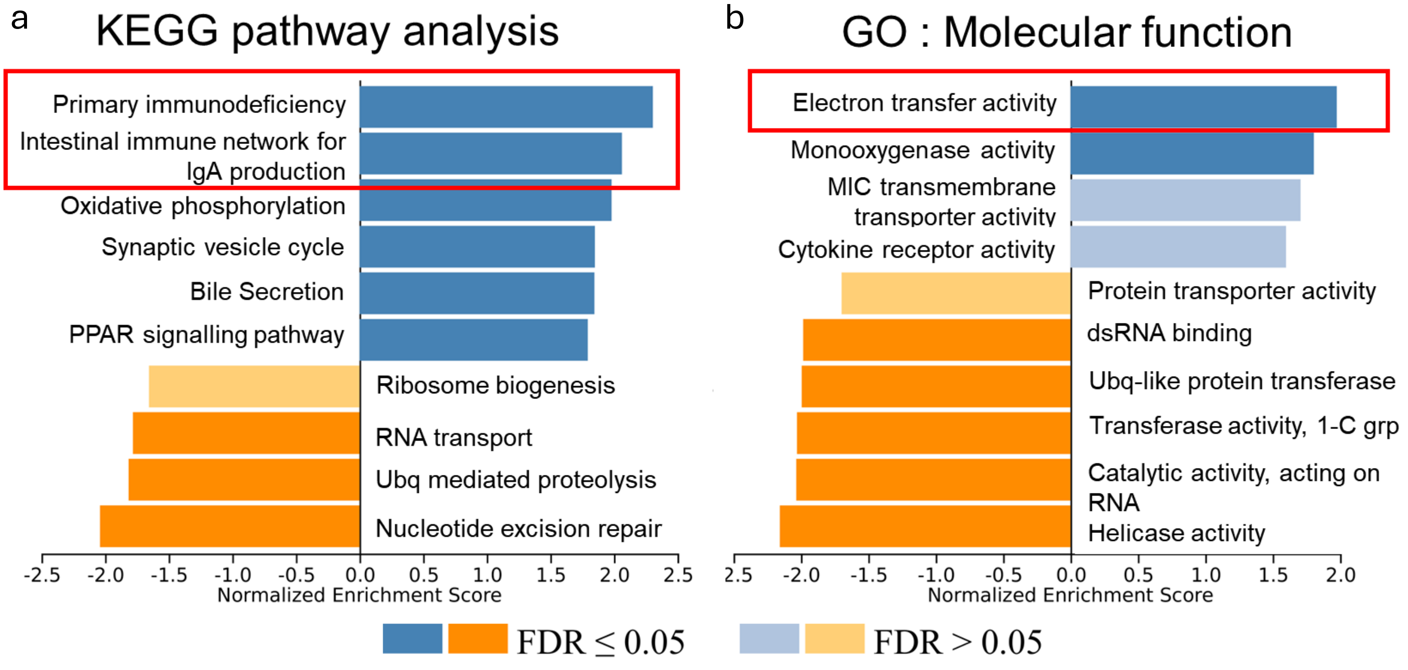


Supplementary Figure 6: Figure showing KEGG pathway analysis a and GO enrichment: Molecular function b results of BMI correlated genes in the CPTAC-3/PDAC dataset generated using LinkedOmicsKB.

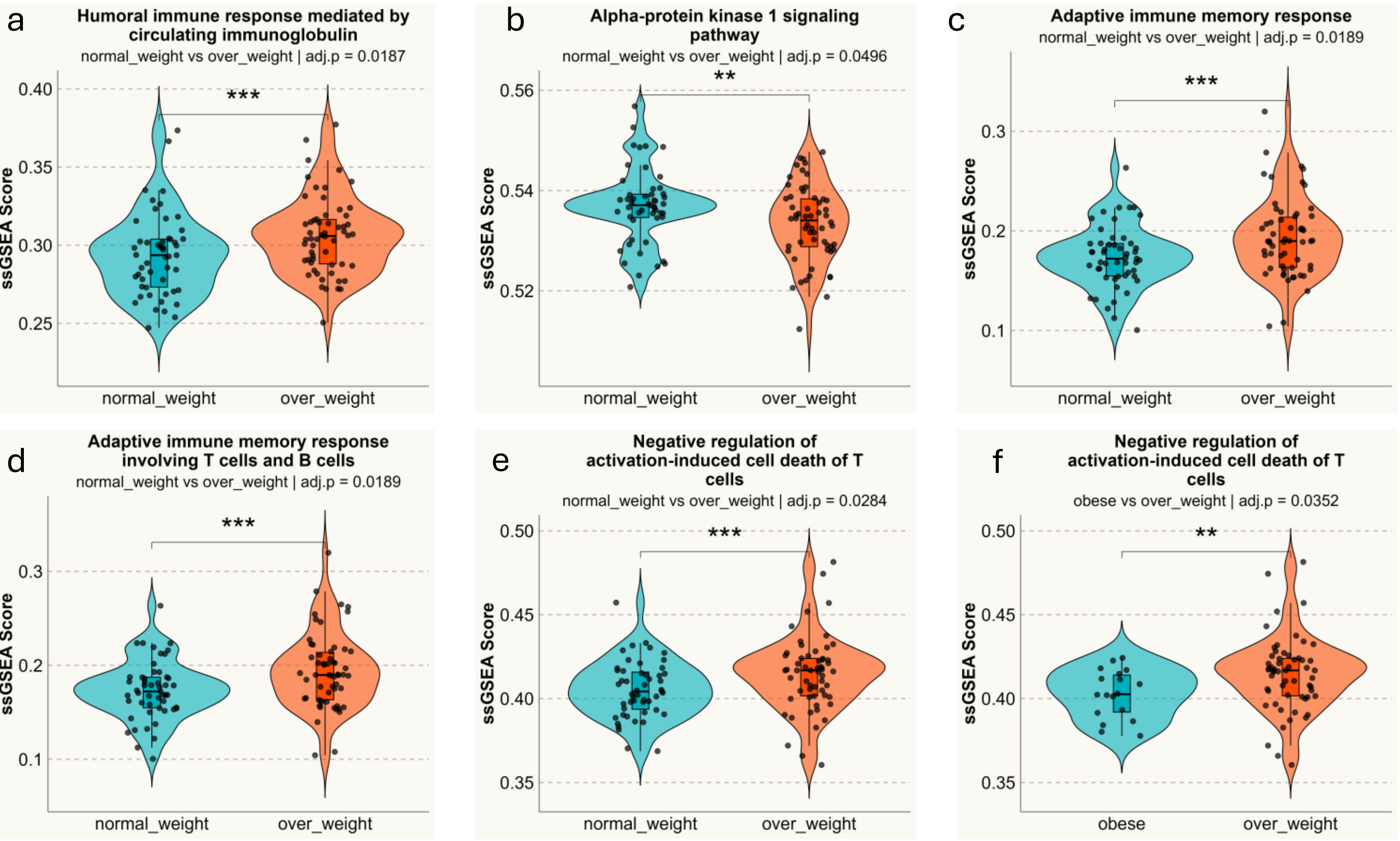


Supplementary Figure 7: Immune pathway activity across BMI groups in pancreatic cancer. Single-sample Gene Set Enrichment Analysis (ssGSEA) of immune-related pathways from the ImmPort database comparing normal weight vs. overweight patients (a-e) and obese vs. overweight patients **f**. All pathways show significantly higher activity in higher BMI groups (Wilcoxon rank-sum test): humoral immune response (p=0.0187), alpha-protein kinase 1 signaling (p=0.0496), adaptive immune memory responses (p=0.0189), T and B cell memory (p=0.0189), and negative regulation of T cell activation-induced death (p=0.0284, p=0.0352). Statistical significance: ****p<0.001, ***p<0.01, **p<0.05, *p<0.1, ns=not significant.

Antigen Processing And Presentation (HSA04612)

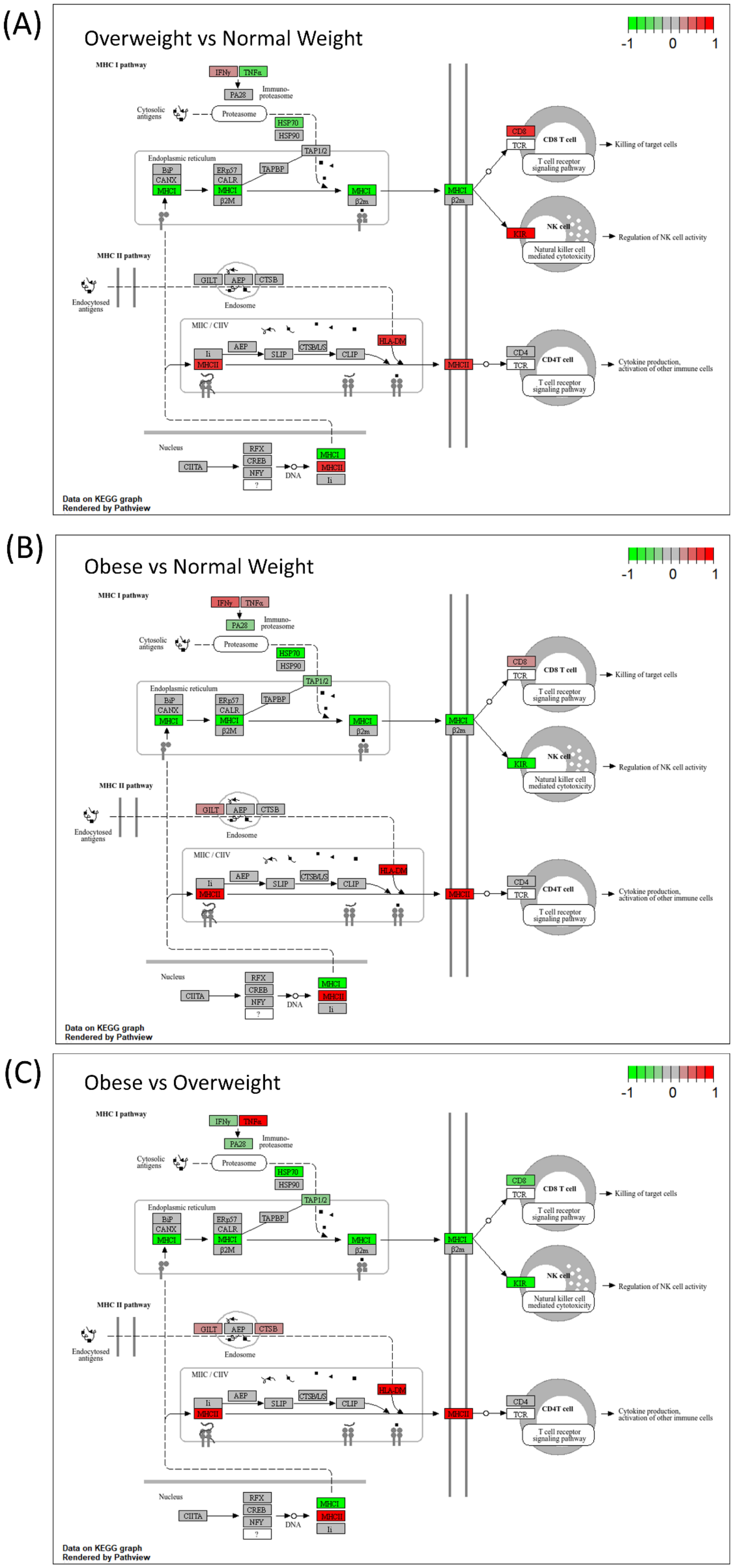


Supplementary Figure 8: Antigen Processing and Presentation Pathway (HSA04612) comparison across BMI groups in pancreatic cancer. KEGG pathview analysis showing differentially regulated genes in the extracellular matrix-receptor interaction pathway between overweight vs. normal weight a and obese vs normal weight b. And obese vs. Overweight c pancreatic cancer patients. Color scale indicates gene expression levels (Green: downregulation [-1]; red: upregulation [+1] in the comparison group relative to the reference group).

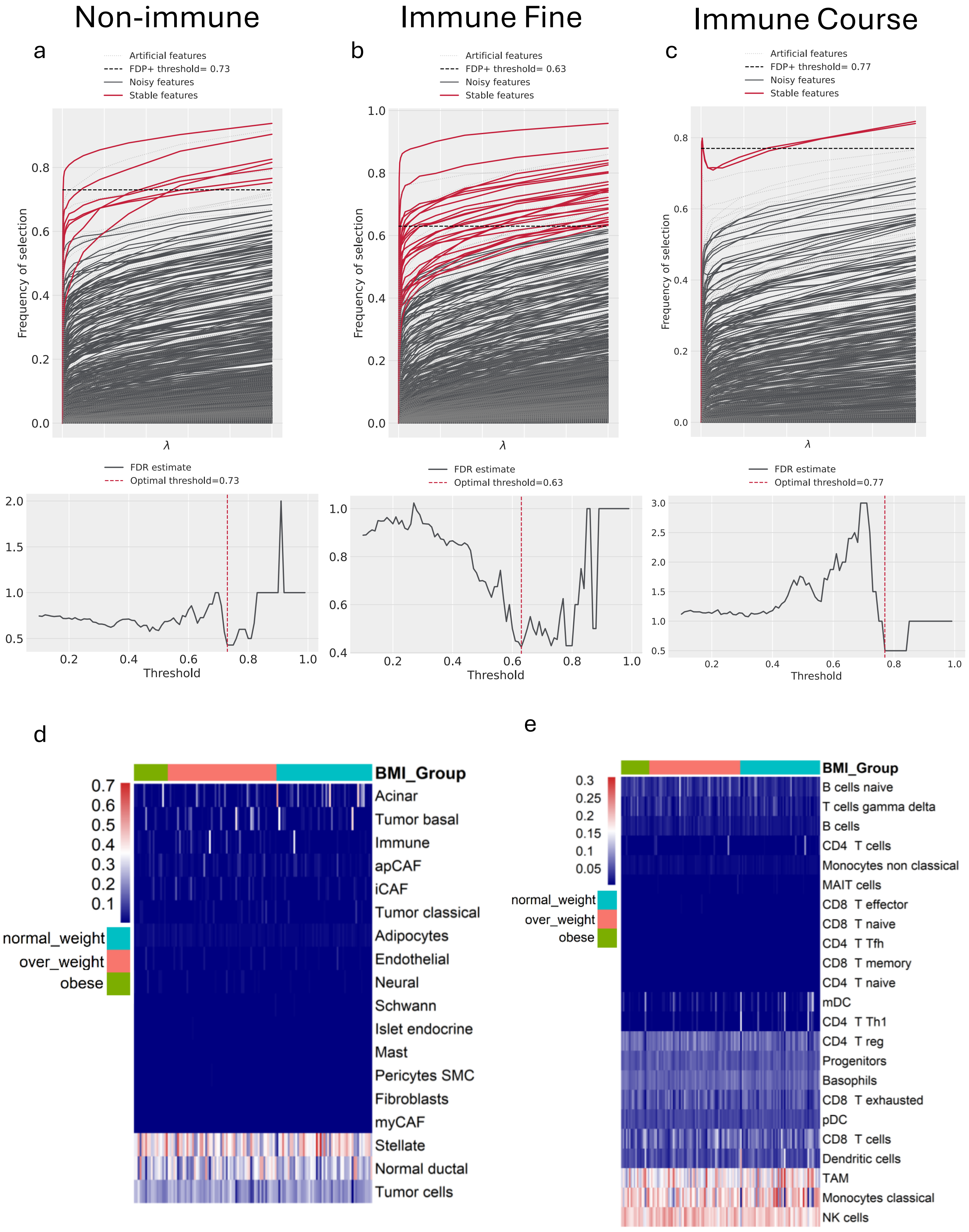


Supplementary Figure 9: (a-c) STABL selected paths showing feature selection frequency across regularization parameter (λ). Below each curve graphs shows FDP estimates as a function of stability threshold. (d & e) heatmap showing cellular proportions after BayesPrism Deconvolution in CPTAC data per patient

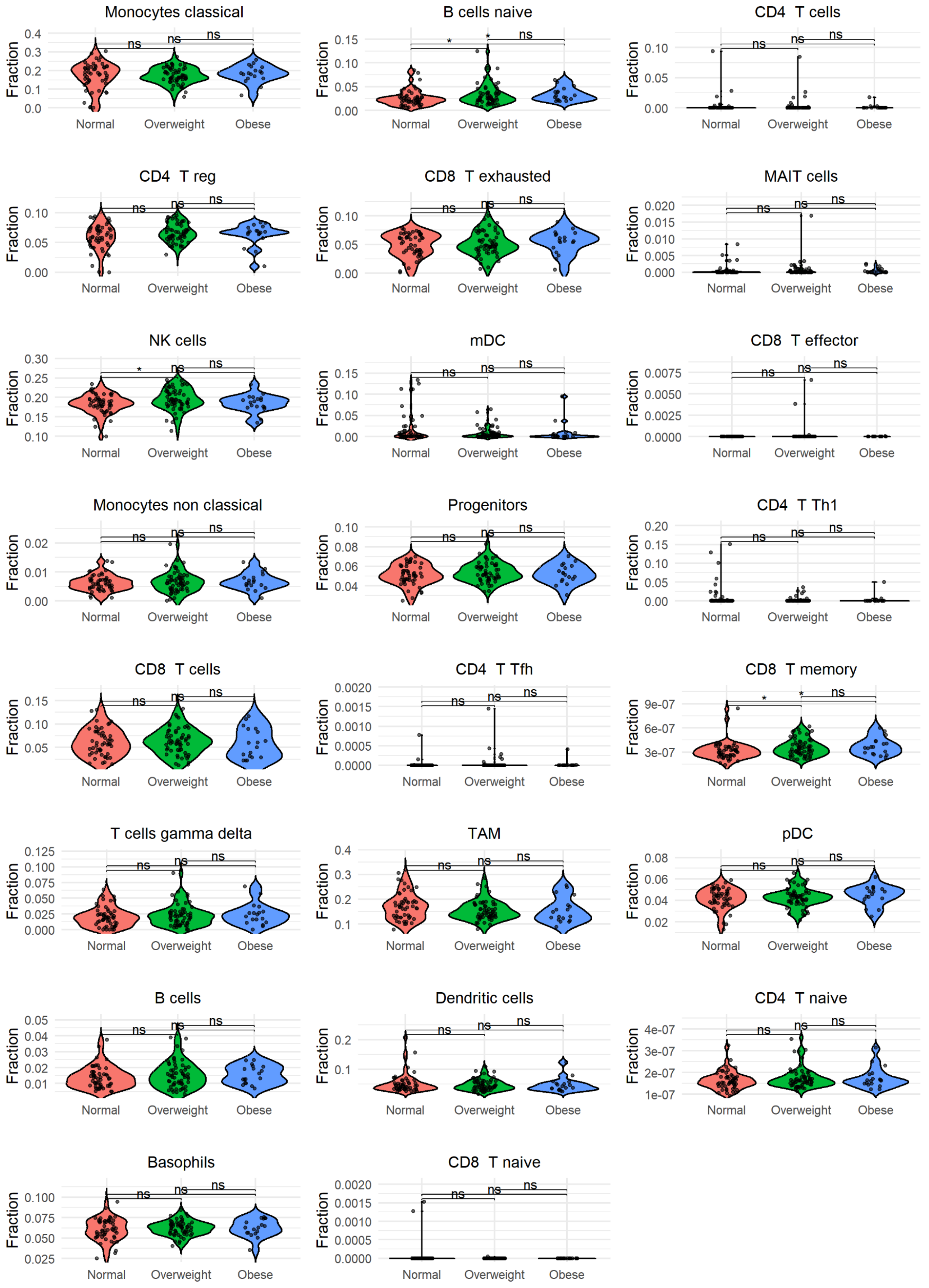


Supplementary Figure 10: Figure showing changes in theta fractions of the immune population, BayesPrism deconvolution. Except for B cell naïve and CD8+ T cell memory cells, no change in theta fractions was observed. Statistical comparisons between BMI groups (Normal Weight, Overweight, Obese) were performed using pairwise Wilcoxon rank-sum tests. Significance levels are denoted as: $*\ P < 0.05$, $**\ P < 0.01$, $***\ P < 0.001$; ns, not significant

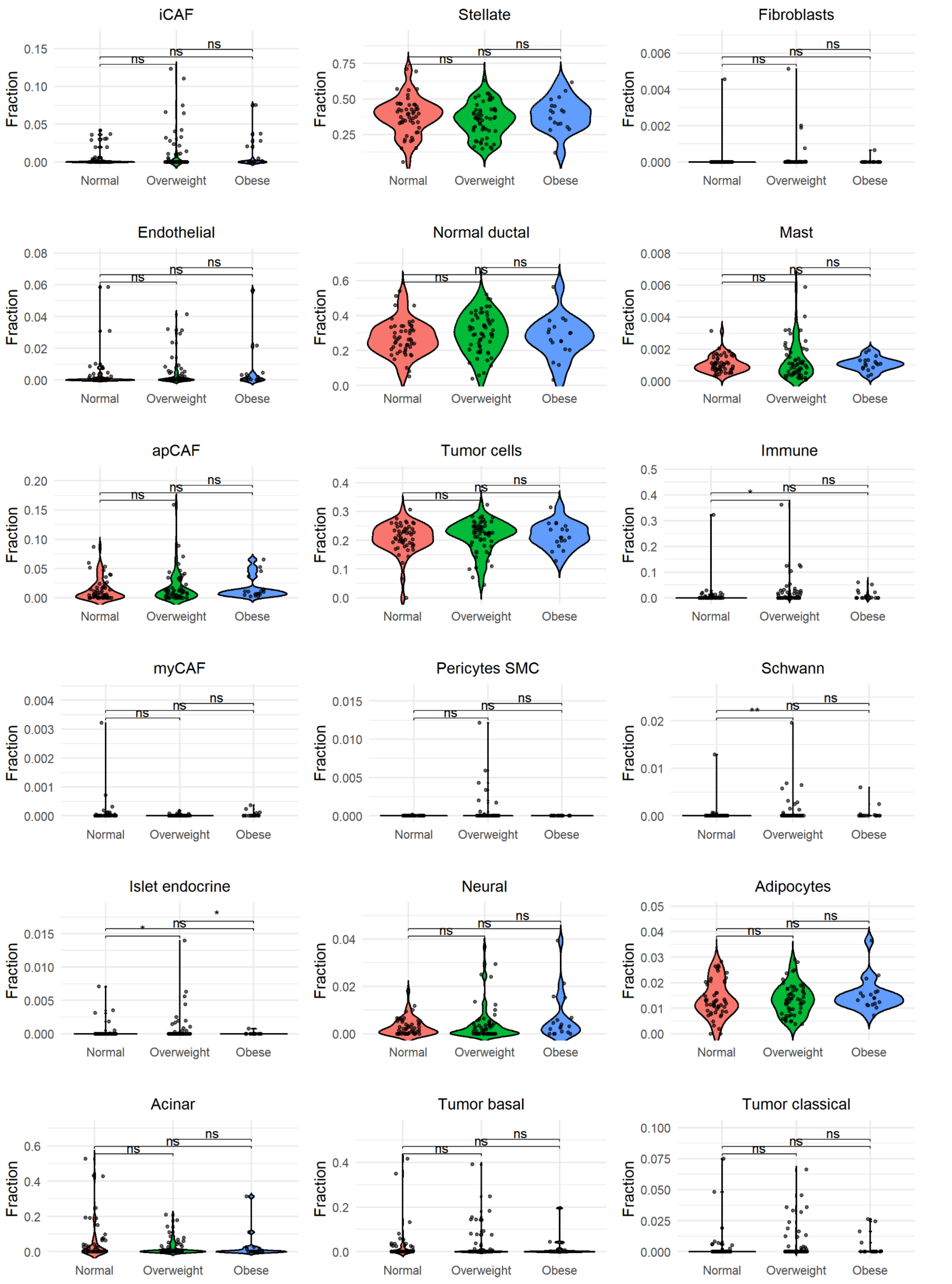


Supplementary Figure 11: Figure showing changes in theta fractions of the non-immune population, BayesPrism deconvolution. Except for the Islet endocrine and Schwann cells, no change in theta fractions was observed. Statistical comparisons between BMI groups (Normal Weight, Overweight, Obese) were performed using pairwise Wilcoxon rank-sum tests. Significance levels are denoted as: $* P < 0.05$, $** P < 0.01$, $*** P < 0.001$; ns, not significant

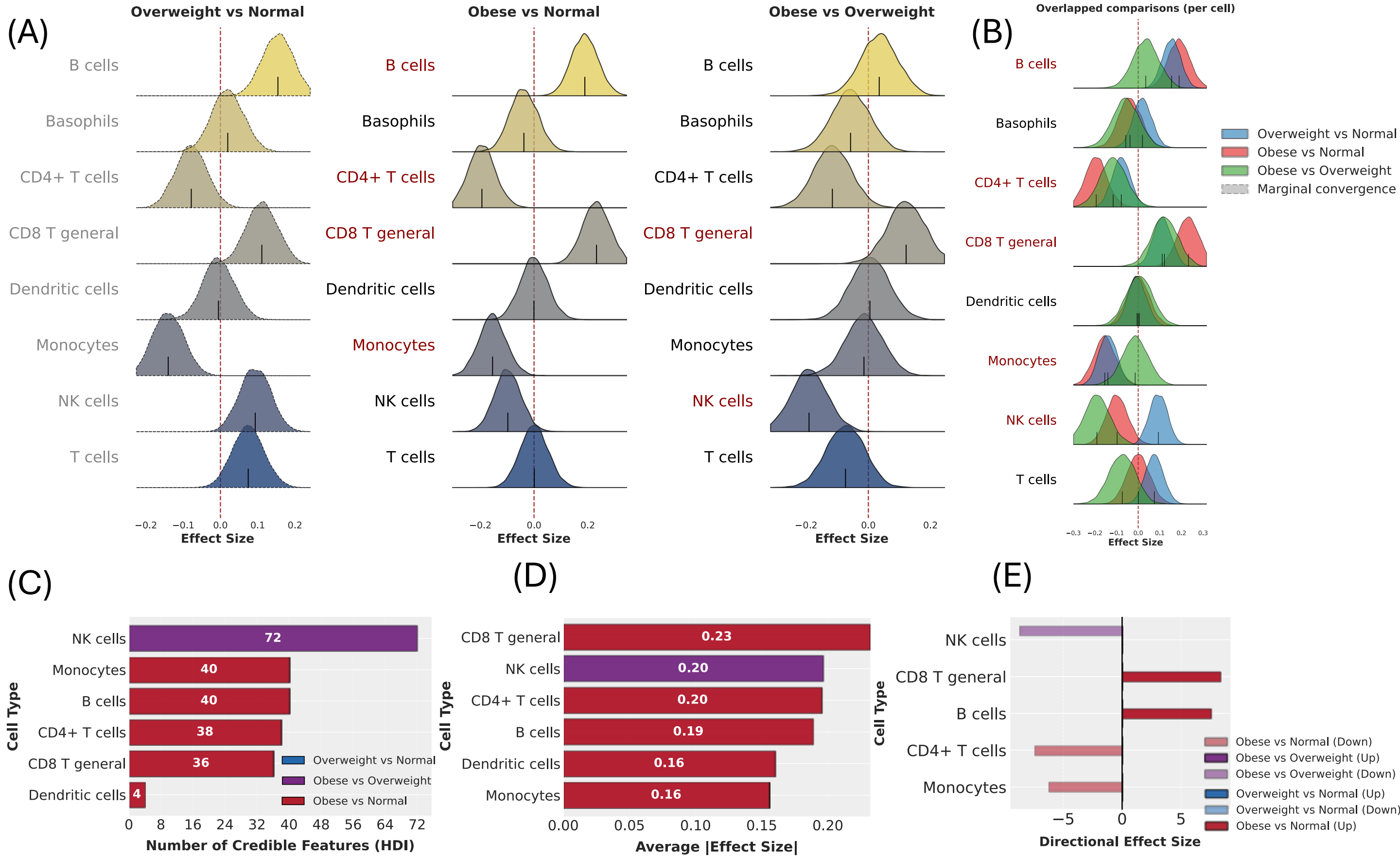


Supplementary Figure 12 : (A–B) Cellular-Level Posterior Distributions in the immune course compartment with broad immune classification. (A) Ridge plots displaying the posterior distributions of pooled cell-type-level effects, separated by comparison. (B) Overlaid ridge plots demonstrating the shift in cellular state distributions across all three comparisons. (C) Stacked bar plot quantifying the number of credible features ($HDI \neq 0$) per cell type, stratified by comparison. (D) Stacked bar plot showing the average magnitude of effect sizes per cell type, highlighting compartments with the most substantial transcriptional shifts. (E) Directionality of effect size changes. The count of significantly upregulated (positive) and downregulated (negative) features across comparisons.

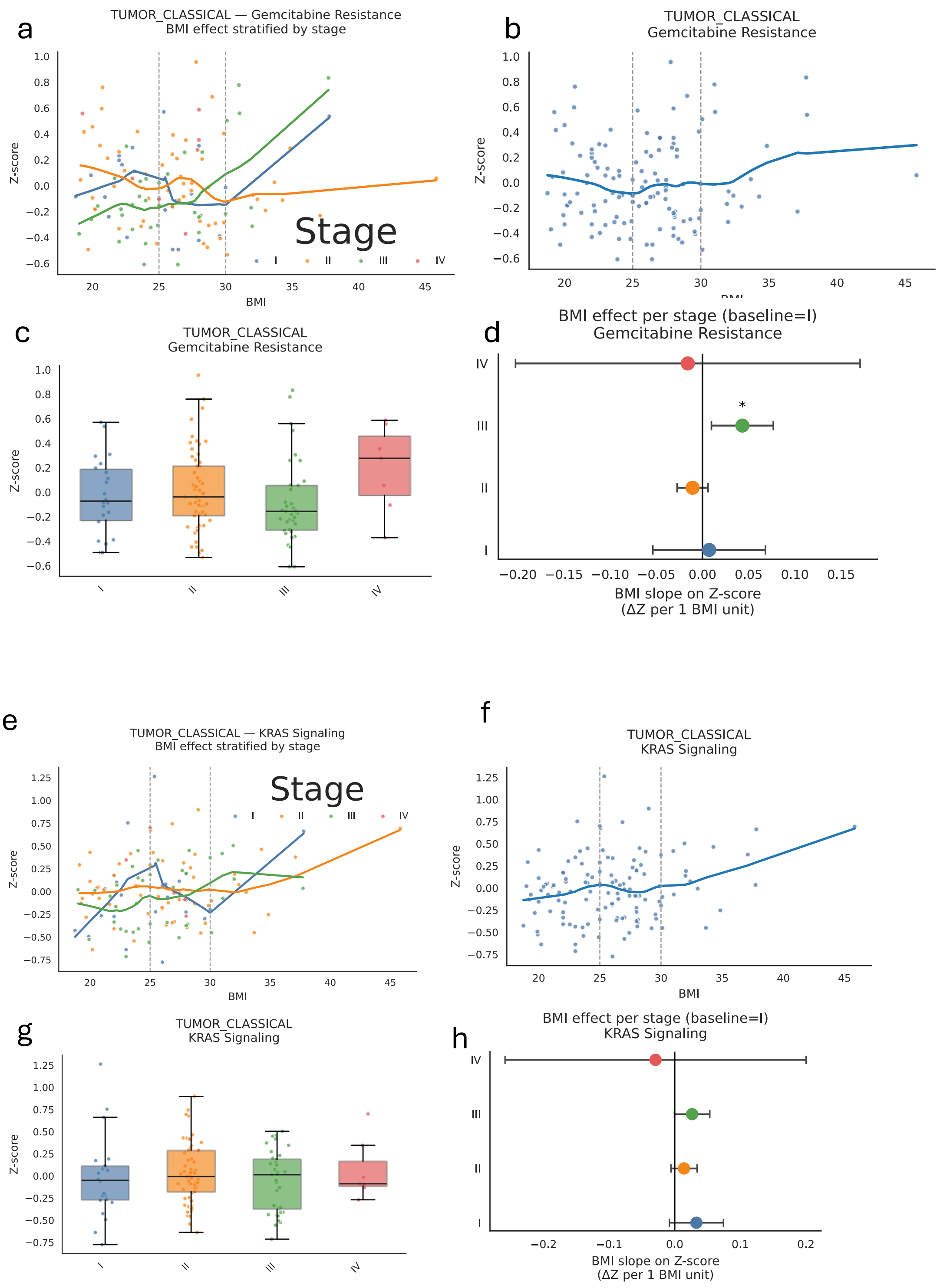


Supplementary Figure 13: Figure showing changes in signature as a continuous function of BMI Tumor classical cells showing changes in Gemcitabine resistance signature and KRAS signaling signature (a,e) stage based changes of signature with BMI change, (b,f) average z score changes of signature with BMI (c,g) Stage based plot showing changes in the signature (d,h) BMI slope changes per one unit BMI change.

### Normal Weight: Bindea

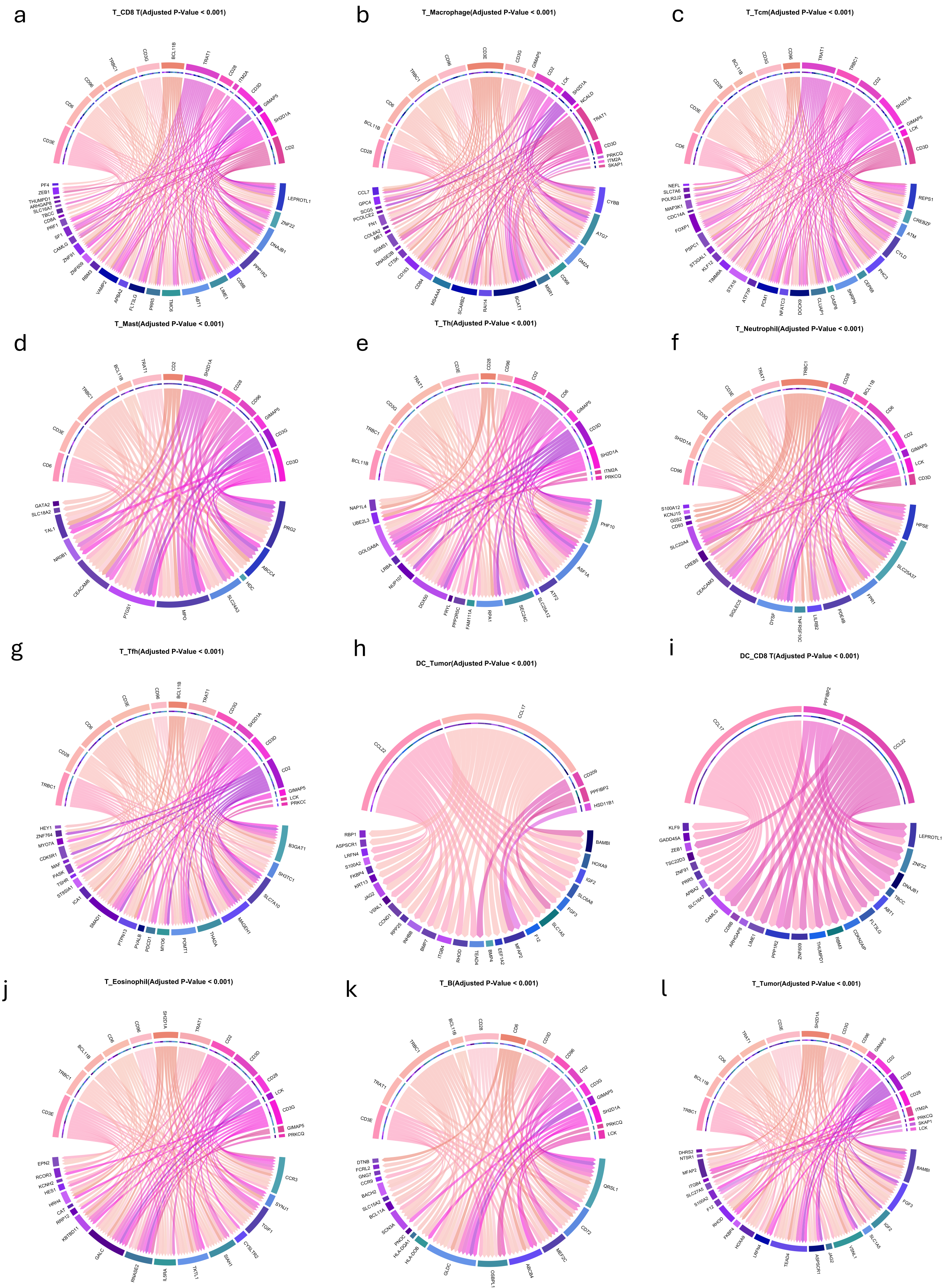


Supplementary Figure 14: Cell-Cell communication in TimiGP analysis in Normal weight patients using Bindea et al.

### Overweight: Bindea

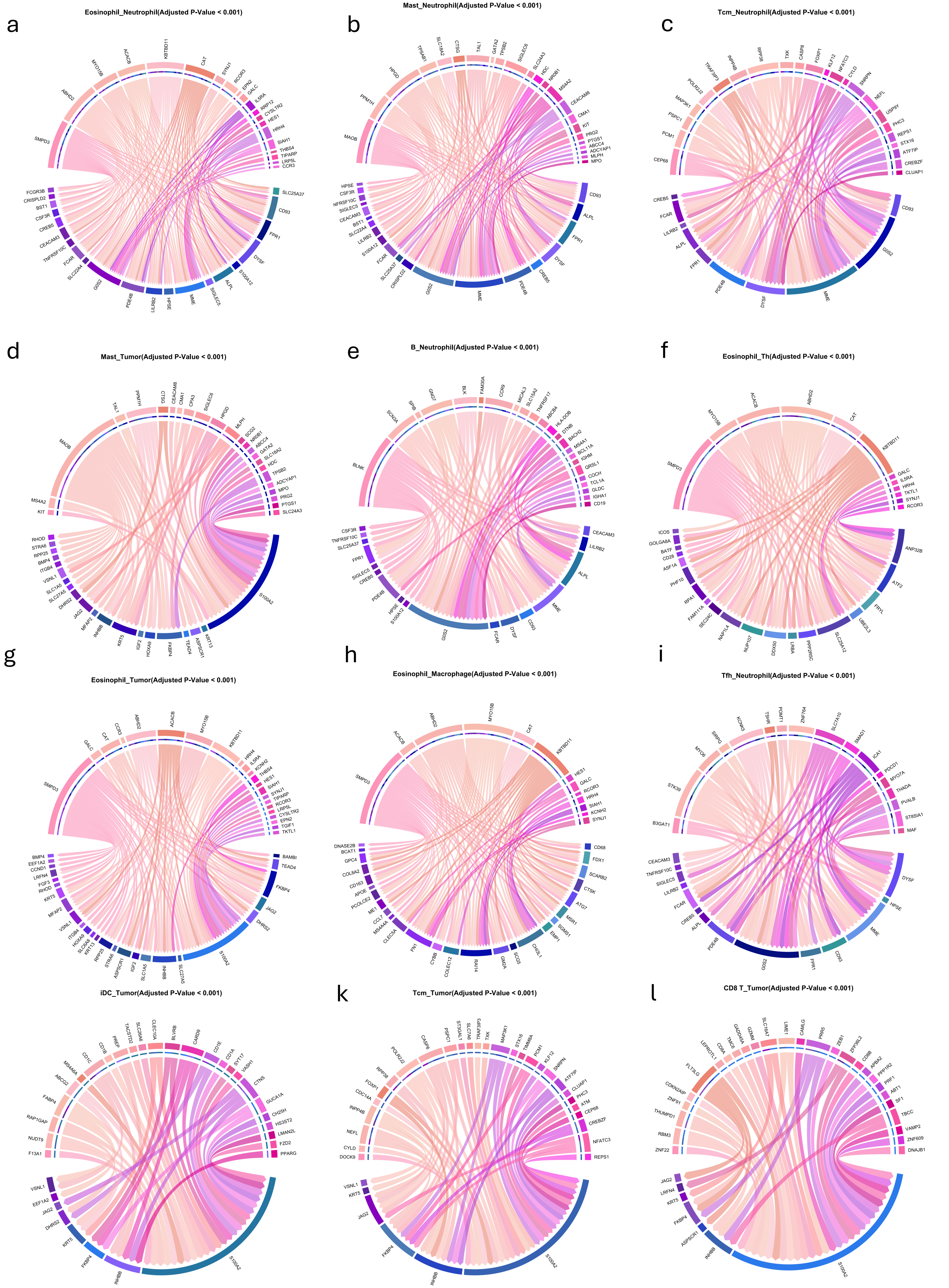


Supplementary Figure 15: Cell-Cell communication in TimiGP analysis in overweight patients using Bindea et al.

### Normal weight: Newman

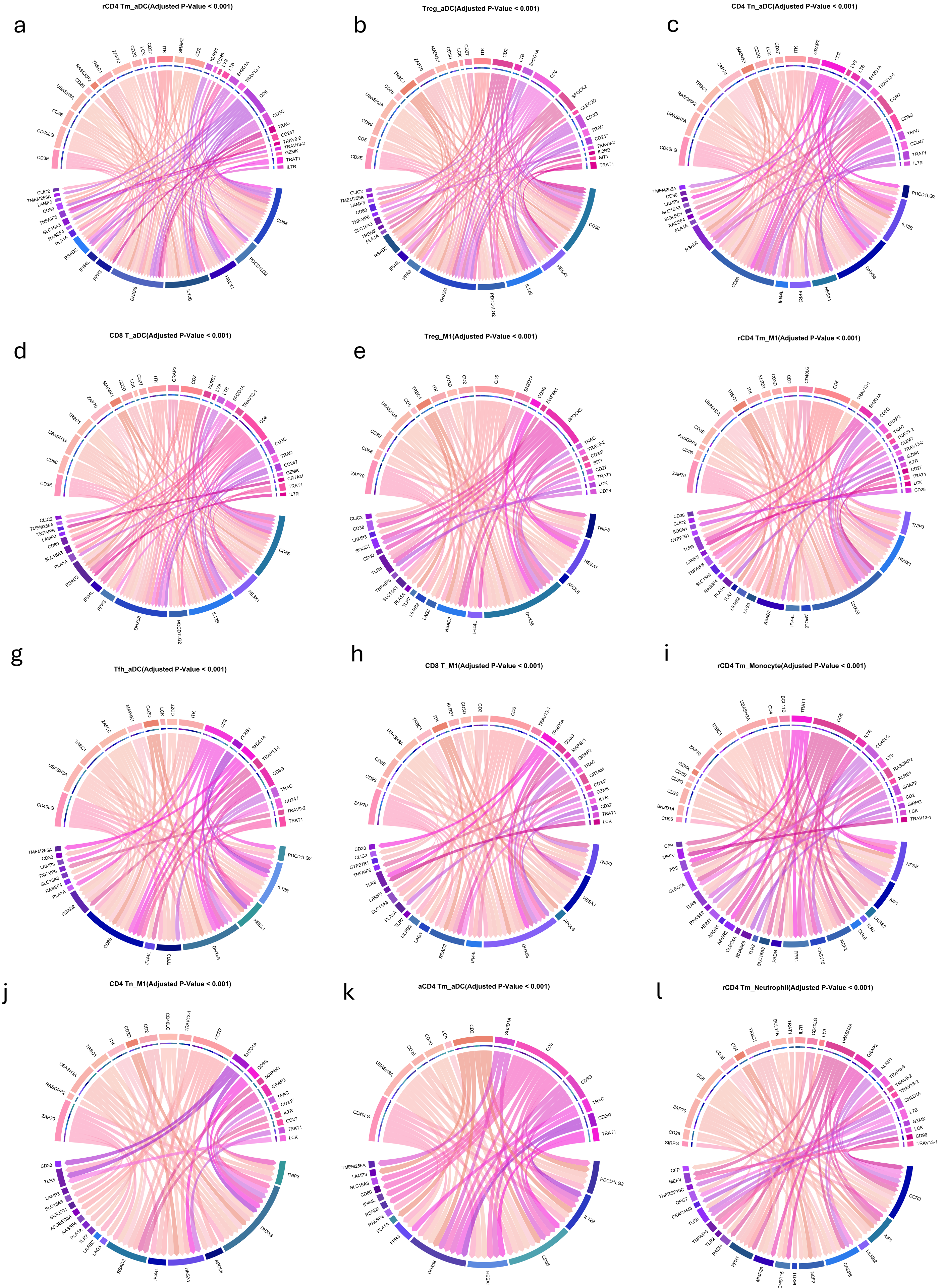


Supplementary Figure 16: Cell-cell communication in TimiGP analysis in normal weight patients using Newman et al.

### overweight: Newman

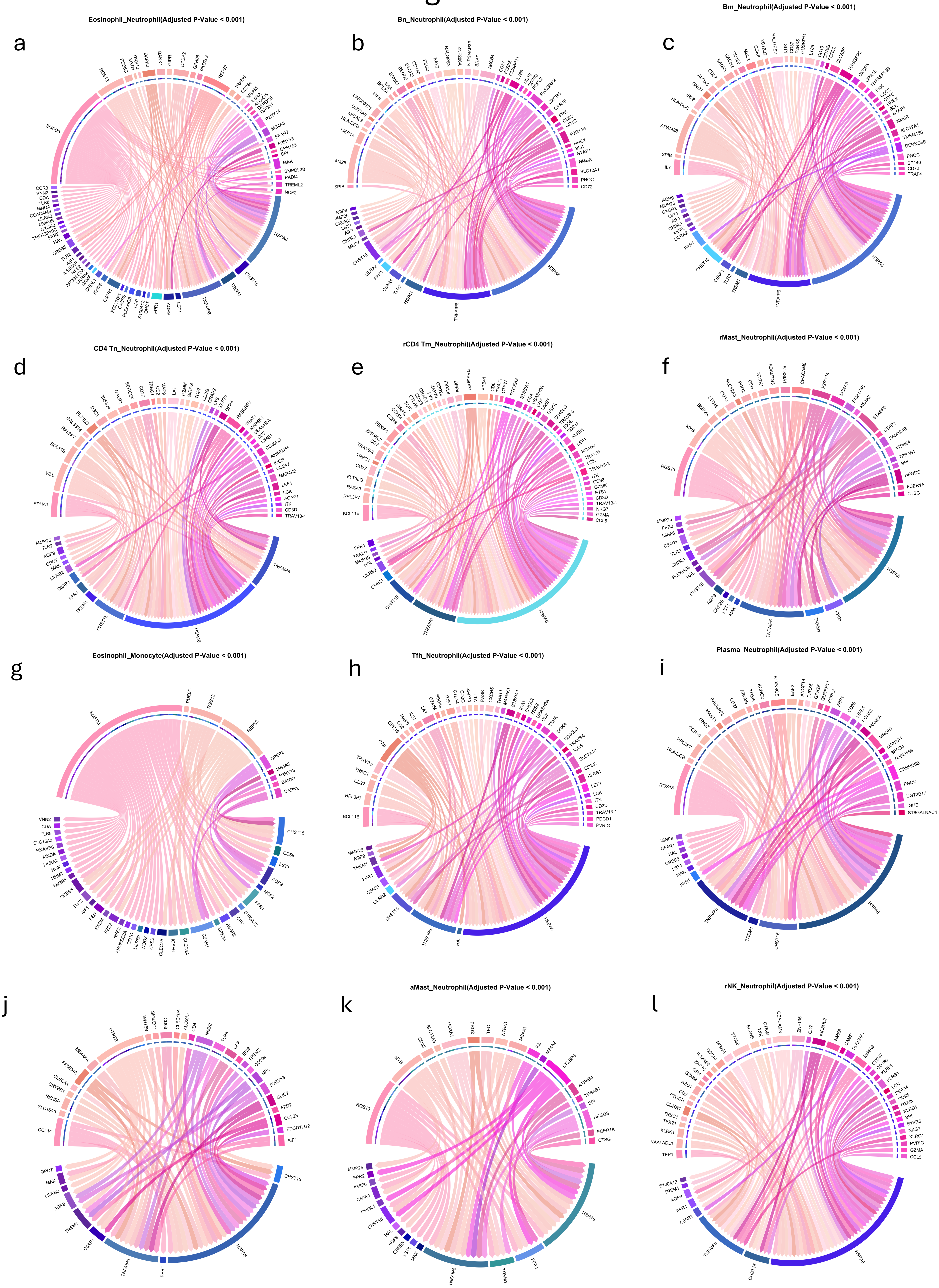


Supplementary Figure 17: Cell-cell communication in TimiGP analysis in overweight patients using Newman et al.

### Normal Weight : Zheng

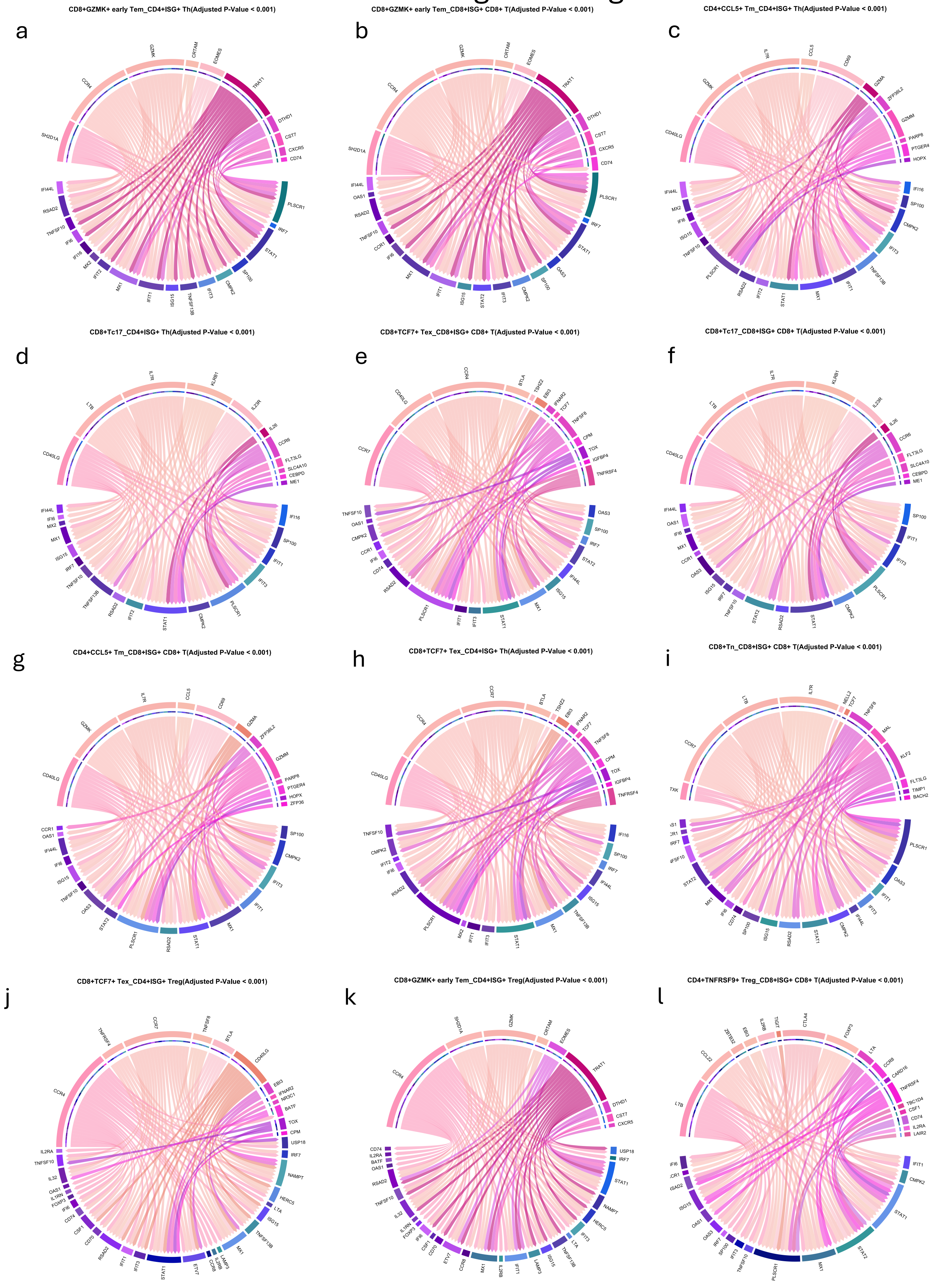


Supplementary Figure 18: Cell-cell communication in TimiGP analysis in normal weight patients using Zheng et al.

### Overweight: Zheng

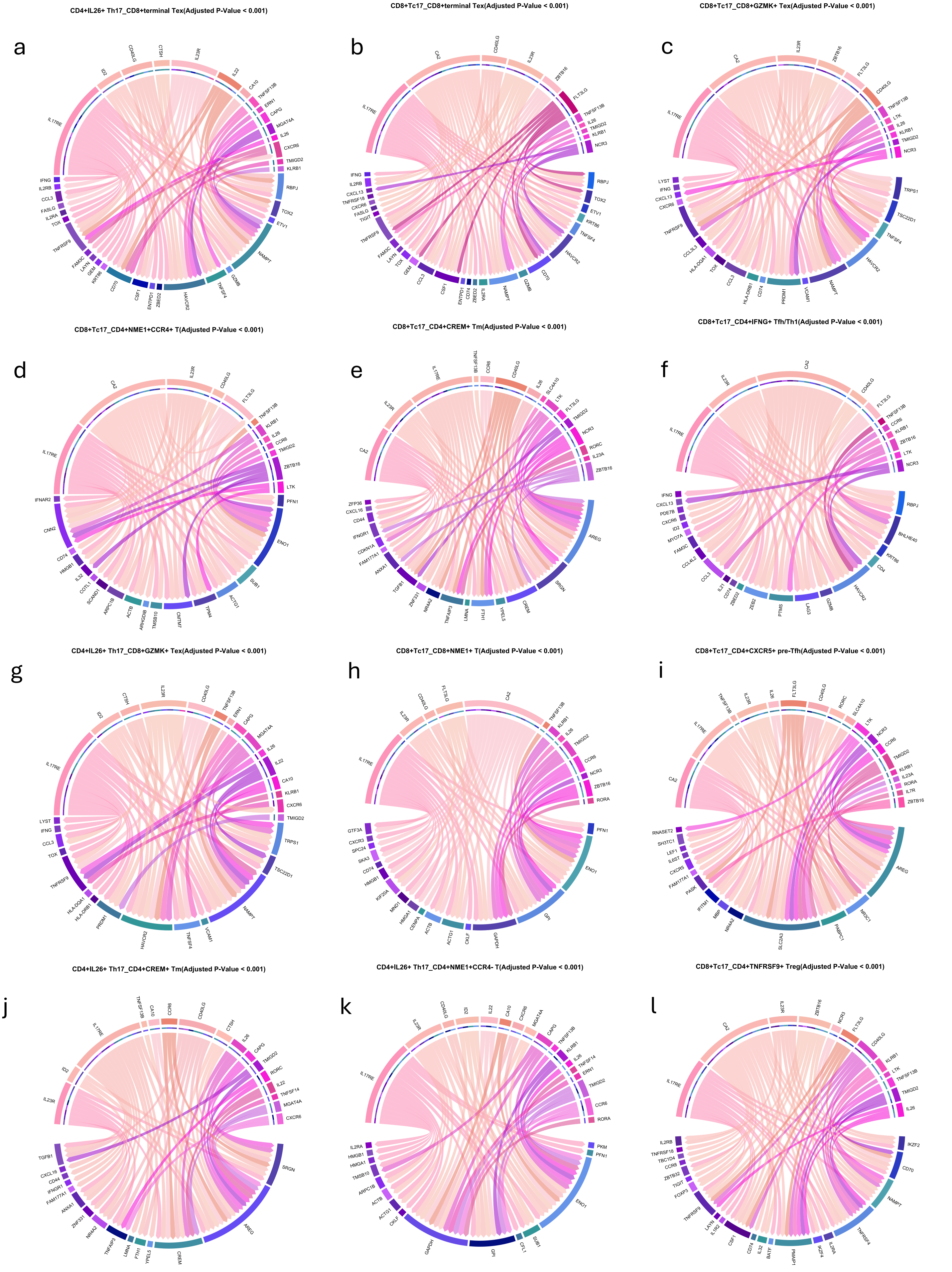


Supplementary Figure 19: Cell-cell communication in TimiGP analysis in overweight patients using Zheng et al.

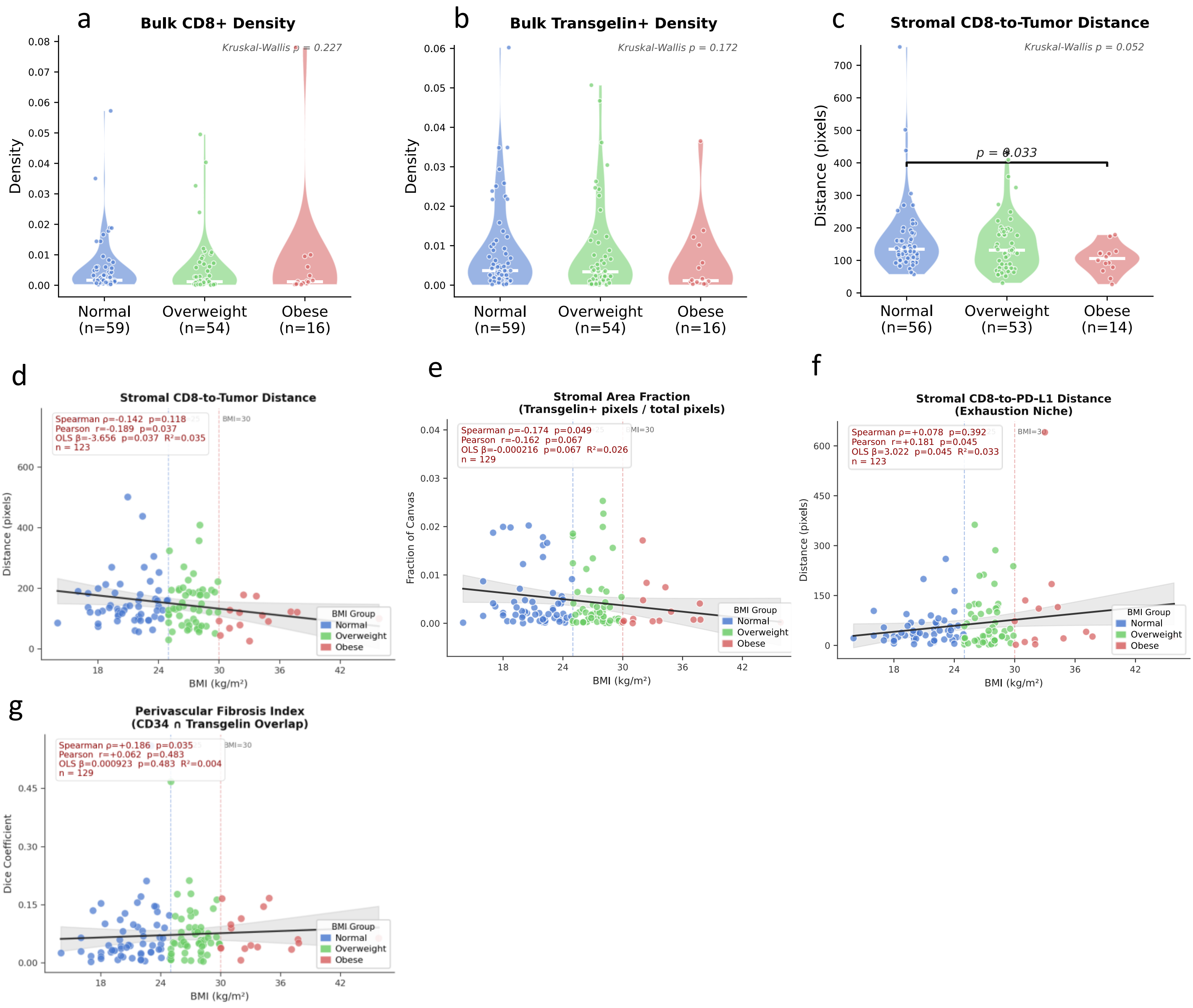


Supplementary Figure 20: Virtual mIF bulk densities and spatial metrics across BMI groups and continuous BMI. Virtual marker densities across BMI groups a CD8+ T cell marker density (Kruskal-Wallis p = 0.227) and b shows Transgelin+ CAF density (Kruskal-Wallis p = 0.172). c shows the stromal CD8-to-tumor boundary distance across BMI groups (Kruskal-Wallis p = 0.052; adj. p = 0.033). d through g show individual patient values plotted against BMI with 95% confidence interval; vertical dashed lines mark the BMI boundaries at 25 and 30. Spearman rho, Pearson r, OLS beta, and sample size are annotated on each panel. d stromal CD8-to-tumour boundary distance (Pearson r = -0.189, p = 0.037, n=123). e stromal area fraction (Spearman rho = -0.174, p = 0.049, n=129). f stromal CD8-to-PD-L1 distance (Pearson r = +0.181, p = 0.045, n=123). g perivascular fibrosis index, quantified as the Dice coefficient between CD34+ and Transgelin+ pixel masks (Spearman rho = +0.186, p = 0.035, n=129).

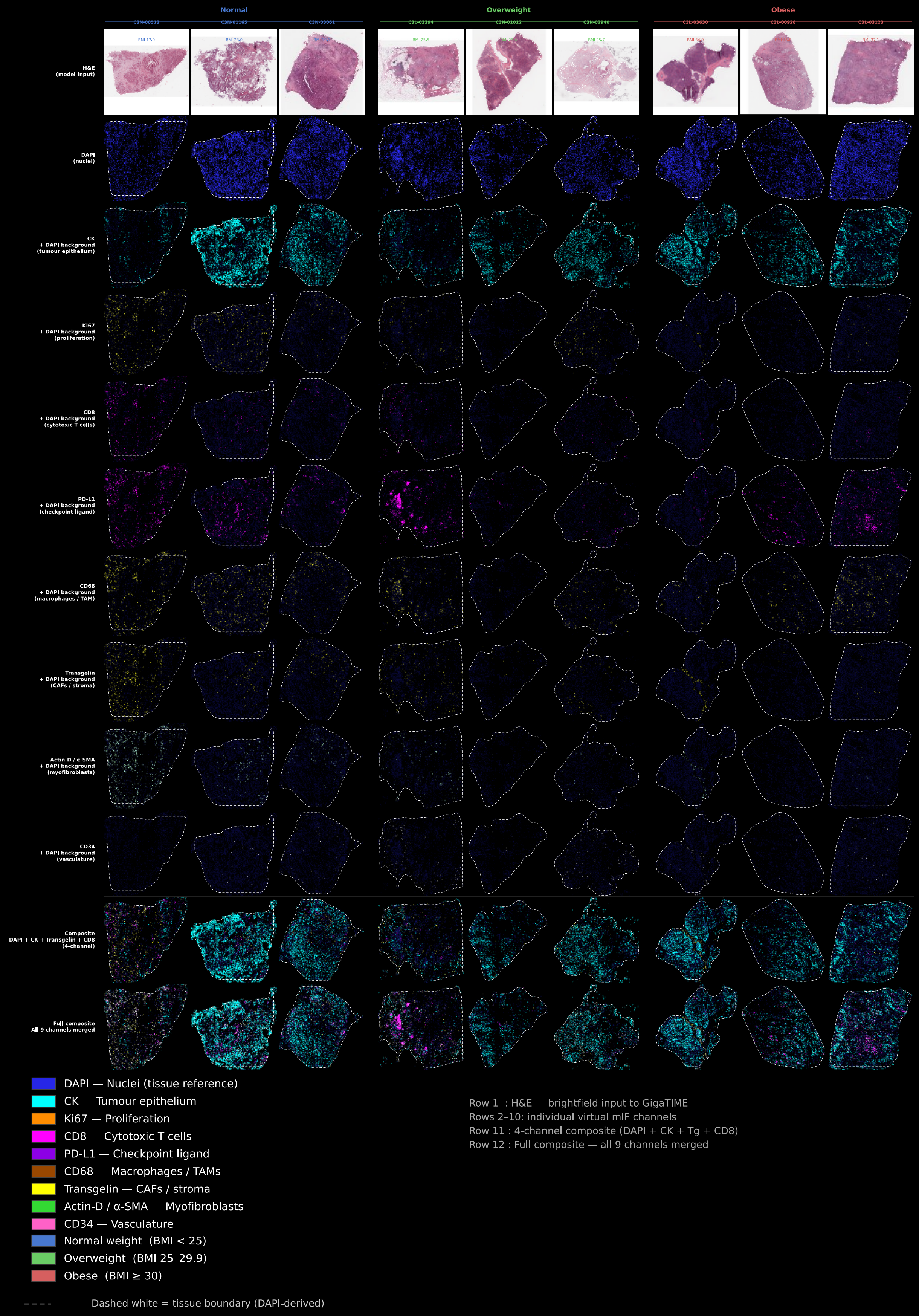


Supplementary Figure 21: GigaTIME virtual mIF output for representative CPTAC-PDA patients.Each column represents one patient. Row 1 shows the H&E brightfield input to the GigaTIME model. Rows 2 to 10 show individual virtual mIF channels with a dim DAPI background included; Row 11 shows the four-channel composite used in the spatial distance analysis (DAPI, CK, Transgelin, CD8). Row 12 shows a full composite merging all nine channels. Dashed white outlines indicate tissue boundaries derived from the DAPI signal. The figure demonstrates that GigaTIME generates spatially coherent signal distributions across tumor epithelial, immune, and stromal compartments in this PDAC cohort.

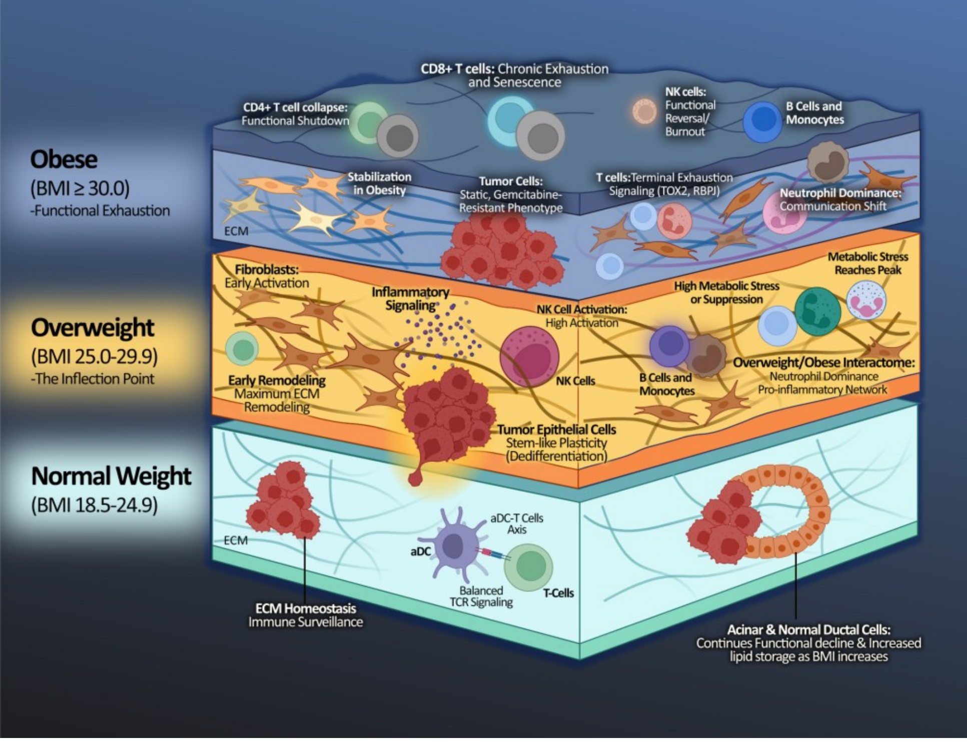


**Functional Reprogramming Across BMI in PDAC**

| Cell Type | Overweight vs. Normal | Obese vs. Overweight | Continues Trend |
|---|---|---|---|
| Fibroblast | High activation | Stabilization | Early remodeling |
| Tumor Cells | Transient dedifferentiation | Absence of plasticity | Therapy resistance |
| B cells | Metabolic stress | Stress plateaus | Early saturation |
| CD4+ T cells (General/Helper) | Functional maintenance | Functional collapse | Total collapse |
| CD4+ Tregs | Metabolic activation | Near complete reversal | Functional collapse |
| CD8+ T cells | Robust activation | No credible increment | Linear progression |
| NK cells | Broad activation | Transcriptional collapse | Functional inversion |
| TAM | Initial suppression | Progressive deepening | Qualitative shift |
| iCAF | Initial activation | Early saturation | Consistent decline |
| Acinar cells | Lipid storage | Effect plateau | Consistent decline |

Supplementary Figure 22: Proposed model of BMI-driven microenvironmental evolution in pancreatic ductal adenocarcinoma (PDAC), highlighting cell type-specific changes across BMI categories (normal weight, overweight, and obese). The accompanying table summarizes cell-type changes identified in both categorical and continuous Bayesian models of obesity-driven PDAC.
