## Supplementary Methods for "Overweight status drives early tumor microenvironment reprogramming in pancreatic ductal adenocarcinoma: a cell-type-resolved Bayesian hierarchical modeling and interactome analysis"

### Supplementary Materials and Methods

#### Bulk Data Differential Expression Analysis

Transcriptomics data from pancreatic ductal adenocarcinoma patients were obtained from the Clinical Proteomic Tumor Analysis Consortium (CPTAC/CPTAC-3) via the Genomic Data Commons (GDC) using the TCGAbiolinks R package. Raw STAR count RNA-seq data from primary tumors of 140 patients with available body mass index (BMI) information were downloaded. Patients were stratified into three groups according to the World Health Organization's BMI classification: normal weight (BMI 18.5-24.9 $kg/m^2$, n = 51), overweight (BMI 25.0-29.9 $kg/m^2$, n = 58), and obese (BMI $\geq$ 30.0$kg/m^2$, n = 18). Raw STAR count matrices were processed using *DESeq2* for differential expression analysis (DEA). Ensembl gene identifiers were converted to HGNC symbols using the *BioMart* tool. Pathway enrichment analysis was performed with *clusterProfiler*, using genes ranked by *DESeq2* test statistics. Gene Set Enrichment Analysis (GSEA) was conducted on Gene Ontology (GO) biological processes, *KEGG* pathways, and *Reactome* pathways. The significance of enrichment was assessed using FDR-adjusted p-values < 0.05, with minimum and maximum gene set sizes of 10 and 500 genes, respectively. KEGG pathway maps were generated using KEGG-Pathview with gene-level log2FC to illustrate alterations in gene expression across selected biological pathways. For *ssGSEA*, gene sets from *ImmPort* were used with VST-normalized counts.

#### Correlation Analysis of BMI and Gene Expression

We conducted a correlation study using the *LinkedOmics* platform to systematically identify genes that vary along with BMI as a continuous variable. The CPTAC-3 pancreatic adenocarcinoma clinical dataset, which includes BMI measures for 140 patients (dataset ID: ID-132290), was analyzed using Pearson correlation with the RNA-seq expression dataset containing 28,057 gene features. The analysis employed RSEM upper-quantile-normalized log2 tumor expression data, ensuring complete sample overlap across both clinical and transcriptomic datasets (n = 140). Genes exhibiting a significant connection with BMI were identified using correlation coefficient thresholds and significance criteria (p-value < 0.05) established by the *LinkedOmics* statistical framework. Functional enrichment analysis of BMI-correlated genes was conducted within the *LinkedOmics* analytical suite, utilizing the integrated *WebGestalt* (Web-based Gene Set Analysis Toolkit) tool. Biological process annotation was performed using GO and KEGG pathway enrichment analyses. The significance Benjamini-Hochberg FDR adjustment, with adj. $p < 0.05$ considered significant. This integrated method facilitated thorough functional characterization of genes whose expression patterns are associated with BMI changes in the PDAC patient group.

#### Deconvolution of CPTAC-PDAC bulk data using Bayes Prism

##### Construction of Single-Cell References

For this paper, two separate single-cell RNA sequencing (scRNA-seq) reference datasets were created from a publicly available PDAC dataset on the Gene Expression Omnibus (GEO). For the whole-tumor reference, we used scRNA-seq data from needle biopsy (n = 25, GSE242230, samples GSM7755911-GSM7755935, total 35,008 cells pre-QC). For immune reference, we used scRNA-seq data from CD45+ enriched tumor-infiltrating leukocytes (n = 3; GSM7502530 from study GSE235452; 6,553 cells post-QC).

##### Quality Control

Dataset-specific adaptive quality control filtering was applied to optimize cell retention while removing low-quality data:

**CD45+ Immune Reference:** Quality gates were set based on data distribution with feature count thresholds at max (200, 1st percentile) for the lower bound and min (6,000, 99th percentile) for the upper bound, total UMI counts <98th percentile, and mitochondrial content <min (20%, 95th percentile). This stringent filtering ensured the construction of high-quality immune cell profiles for deconvolution reference construction.

**Tumor Microenvironment Reference:** To preserve heterogeneous cell populations, including rare stromal and acinar cell types, slightly more permissive thresholds were employed: feature counts between max (200, 0.5th percentile) and min (8,000, 99.5th percentile), total UMI counts <99th percentile, and mitochondrial content < min (25%, 95th percentile). These adaptive gates accommodated the broader transcriptional diversity expected in whole-tumor samples while maintaining data quality.

Cells falling outside these ranges were excluded from downstream analysis in each respective dataset.

##### Malignancy Classification and Tumor Subtyping

To remove possible contamination by CD45+ cells, stromal cells, or epithelial cells in the scRNA-seq data, a primary gating strategy was employed based on PTPRC (CD45) expression ≥25% of non-zero PTPRC values. Additionally, an immune score was calculated using pan-immune markers (PTPRC, CD3E, CD19, CD14, FCGR3A, CD68, MS4A1). Cells are considered immune if PTPRC expression exceeded the threshold or if ≥25% of pan-immune markers were expressed. Stomal and epithelial contamination were excluded based on the stomal (*str_score*) and epithelial score (*epi_score*). *Epi_score* were calculated using six markers (EPCAM, KRT8, KRT18, KRT19, KRT7, TACSTD2) and *str_score* were calculated using six markers (COL1A1, COL1A2, COL3A1, DCN, BGN, SPARC). Within each sample, epithelial scores were calculated as the sum of these marker counts, and cells scoring above the 70th percentile (i.e., the top 30% per sample) were classified as epithelial (n = 7,822 cells analyzed).

Copy number variation analysis was performed using *CopyKat* to distinguish malignant from normal epithelial cells. Prior to *CopyKat* analysis, mitochondrial and ribosomal genes were removed, and genes expressed in <1% of epithelial cells were filtered. *CopyKat* inferred chromosomal copy number profiles using parameters *ngene.chr=5*, *win.size=25*, and *KS.cut=0.1*, classifying cells as aneuploid (malignant) or diploid (normal). Malignant epithelial cells (aneuploid by *CopyKat*) were further subtyped into basal-like and classical PDAC subtypes using signature-based module scoring. Scores were calculated for basal-like (S100A2, KRT6A, KRT17, SERPINB4, LY6D, HMGA2, TP63, KRT5, DSC3, PKP1) and classical (TFF1, LGALS4, TSPAN8, REG4, ST6GALNAC1, CTSE, GATA6, HNF1A, HNF4A, FOXA2, FOXA3, PDX1) signatures. Malignant cells with basal scores exceeding both classical scores and a threshold of 0.20 were classified as basal-like tumors, while those with dominant classical signatures (>0.20) were classified as classical tumors. Remaining malignant cells were designated as unclassified tumor cells. Normal ductal epithelial cells were defined as *CopyKat*-diploid cells or cells with elevated normal ductal marker expression (CFTR, BICC1, SLC4A4, GLIS3, SCTR, AMBP, CA2; module score >0.40).

**Stromal and Immune Cell Classification**

Non-epithelial cells were classified using a marker-based module scoring implemented via Seurat's *AddModuleScore* function. Cancer-associated fibroblasts (CAFs) were identified by quiescent fibroblast marker expression (DPT, DCN, LUM, COL1A1, COL1A2, COL3A1, BGN; score >0.15) and subdivided into functional subtypes: myofibroblastic CAFs (myCAF:

ACTA2, TAGLN, MYL9, POSTN, MMP11, HOPX, MYH11, CNN1, MYLK; score >0.30), inflammatory CAFs (iCAF: IL6, CXCL12, CXCL14, CCL2, PDGFRA, CFD, PLA2G2A, CD34; score >0.30), and antigen-presenting CAFs (apCAF: HLA-DRA, HLA-DPB1, CD74, CIITA, SLPI, HLA-DPA1; score >0.30).

Vascular cells were classified as blood endothelial (PECAM1, VWF, CDH5, KDR, FLT1, FLI1, ERG; score >0.35), lymphatic endothelial (PROX1, LYVE1, FLT4, PDPN, CCL21; score >0.35), or pericytes/smooth muscle cells (RGS5, PDGFRB, CSPG4, MCAM, NOTCH3, ABCC9, ACTA2, MYH11, TAGLN, CNN1, MYLK, CASQ2, KCNAB1; score >0.35). Pancreatic stellate cells were identified using combined markers (RGS5, PDGFRB, CSPG4, MCAM, NOTCH3, ABCC9, GFAP, SPARC, VIM, ACTA2, TIMP1, COL1A1, DES; score >0.30), and acinar cells by acinar-specific markers (PRSS1, CPA1, CTRB1, CELA3A, AMY2A, PNLIP, CTRB2, CTRC; score >0.30).

#### CD45+ Immune Cell Deconvolution

##### Immune Cell Type Annotation

Following quality control and normalization, cells underwent dimensionality reduction and clustering. Initial filtering was done using PTPRC (expression >0), followed by more robust immune gating (PTPRC, CD3E, CD19, CD14, FCGR3A, CD68, MS4A1). Epithelial and stromal cells were filtered out using the corresponding marker expression as described earlier (*epi_score* > 0.5 and *str_score* > 0.5). Cell type annotation employed a multi-reference consensus strategy using *SingleR* with four immune reference databases accessed via the celldex R package: Monaco Immune Database, DICE (Database of Immune Cell Expression), Blueprint, and Novershtern hematopoietic references. *SingleR* was executed at both main and fine annotation levels with optimized parameters: differential expression method="wilcox", top genes de.n=50 (main level) or 20 (fine level), fine-tuning enabled (fine.tune=TRUE, tune.thresh=0.05 for main, 0.1 for fine), standard deviation thresholds (sd.thresh=1 for main, 1.5 for fine), and low-confidence label pruning enabled (prune=TRUE). Using Monaco an *immune coarse* CD45+ scRNA reference were established with B cells (n = 341), Basophils (n=24), CD4+ T cells (n = 876), CD8+ T cells (n =1092), Dentritic cells (n = 369), Monocytes (n = 1970), NK cells (n = 478), Progenitors (n = 14), and T cells (n = 1389).

Monaco fine-level Consensus cell type labels were generated by integrating Monaco and DICE fine-level annotations with lineage-specific conflict resolution rules. T cell subsets were classified by combining CD3 positivity with CD4/CD8 marker expression patterns from Monaco annotations. Myeloid cells were distinguished from dendritic cells using DC-specific markers (FLT3, IRF8, CLEC9A). CD4+ regulatory T cells were identified based on DICE annotations containing "TREG" patterns. CD8+ exhausted T cells were defined by multi-marker criteria: (1) assignment to CD8+ T cell subsets, (2) mean exhaustion marker expression (PDCD1, HAVCR2, LAG3, TIGIT, TOX) above the 80th percentile among CD8+ T cells, and (3) exhaustion module score >0.5.

**Tumor-Associated Macrophage Identification**

Tumor-associated macrophages (TAMs) were distinguished from circulating monocytes using multi-criteria signature-based scoring. Module scores were computed for monocyte markers (LYZ, S100A8, S100A9, FCN1, MS4A7, CTSS, VCAN, TYROBP, CCR2, LST1, FCGR3A, CX3CR1) and macrophage markers (CD68, MSR1, CD163, MRC1, C1QA, C1QB, C1QC, APOE, MARCO, SIGLEC1, TREM2, SPP1) using Seurat's AddModuleScore function. The macrophage-to-monocyte score difference (Delta_MM = MacroLin1 - MonoLin1) was calculated for cells within the monocyte lineage (Monocytes, Monocytes classical, Monocytes non-classical, Macrophage) and z-score normalized globally.

TAM candidates were selected using adaptive thresholding with three mandatory criteria: (1) Delta_MM z-score ≥1.0-1.8 (adaptively reduced from 1.8 in 0.1 increments until ≥30 TAM cells were identified), (2) expression of ≥3 TAM hallmark genes (C1QA, C1QB, C1QC, APOE, TREM2, SPP1, SIGLEC1, MARCO, MRC1, CD163) above gene-specific medians, and (3) exclusion of cells scoring in the top 10% for dendritic cell (FLT3, IRF8, CLEC9A, XCR1, CD1C, FCER1A, LAMP3, CCR7) or neutrophil (CXCR2, FCGR3B, MMP8, CSF3R, ELANE, MPO) signatures. Monaco-annotated dendritic cells were explicitly excluded from TAM consideration. This yielded 211 TAM cells.

The final immune reference comprised 21 well-represented cell types (≥20 cells): classical monocytes (n=1,419), non-classical monocytes (n=79), TAMs (n=211), myeloid DCs (n=557), plasmacytoid DCs (n=35), CD4+ T cells general (n=89), CD4+ Th1 (n=555), CD4+ Treg (n=453), CD4+ Tfh (n=61), CD4+ naive (n=43), CD8+ T cells general (n=540), CD8+ effector (n=859), CD8+ memory (n=149), CD8+ exhausted (n=295), CD8+ naive (n=31), NK cells (n=478), MAIT cells (n=208), gamma-delta T cells (n=78), naive B cells (n=332), dendritic cells (n=36), and basophils (n=23). Two minor populations (Progenitors: 14 cells, B cells general: 8 cells) had limited representation.

**Three-Phase BayesPrism Deconvolution**

BayesPrism was conducted with Gibbs sampling for 2000 iterations, a burn-in period of 500, and a thinning factor of 2; outliers were defined at 0.05 and cut at 0.01 for all phases.

**Development of Custom Signature Database**

Initial Curation and Scope: Signatures specific to cell types were established based on existing literature and functional pathway databases. Signatures encompassed several aspects of immunological failure, including fatigue states, metabolic reprogramming, senescence indicators, stromal exclusion mechanisms, and functional impairment. Comprehensive signature panels were developed for each principal immune cell type (CD8+ T cells, CD4+ T cells, B cells, macrophages, and dendritic cells), and non-immune cell type encompassing essential functional states, including cytolytic activity, functional signaling, oxidative stress response, mitochondrial function, autophagy regulation, and tissue residency programs.

**Removal of redundant signature and LLM-assisted refinement**

To remove redundant genes in signatures derived from different literature sources, we performed pairwise overlap analysis within each cell type. We utilized the Overlap Coefficient ($O = |A \cap B| / \min(|A|, |B|)$) to measure similarity between gene sets. Signatures exhibiting an Overlap Coefficient $> 0.50$ were flagged for manual review. For each flagged pair, we examined the shared and unique genes to determine whether the signature is a subset, an overlap, or distinct. A Python framework integrating the Google Gemini API (gemini-2.5-flash) to assist the evaluation, with low temperature settings (temperature = 0.2) to ensure a consistent response. The model was prompted to assess the biological context "obesity and pancreatic cancer" and suggest whether to retain, remove, or merge the signature. All final decisions were made manually after reviewing gene overlap and model suggestion.

To discover new signatures, the model was asked to propose new signatures relevant to obesity-driven dysfunction in pancreatic cancer. Each signature required 8-12 genes and had fewer than 20% overlaps with any existing signature within the cell type. Suggestions failing this criterion were removed. Remaining suggestions were manually reviewed using literature evidence before being included in the genset.

A total of 2143 signatures across 65 cell types, with 10 average genes per signature across three compartments: non-immune, immune fine, and immune coarse. This signature database is available in an in-house Streamlit app at https://obese-pdac-model.streamlit.app/ under the 'Signature Explorer' section.

**Calculation of Signature Score from Deconvoluted Data**

Signature score is calculated using a standardized Z-score approach. Briefly, gene expression values were mean centered without log transformation. To prevent mean Z-score inflation, scores are capped at ±3 winsorization. Signature scores were computed as the mean Z-score of all detected signature genes. Any missing values were imputed with 0 to ensure neutral contribution, and only signatures with at least 4 detectable genes were retained for analysis.

**Stabl, Machine Learning-Based Signature Selection**

To prioritize gene signatures associated with BMI in a high-dimensional setting, we employed a machine learning algorithm called Stabl (v1.0.0) (2) (https://github.com/gregbellan/Stabl). Before model fitting, samples with missing BMI metadata were excluded from the analysis. The remaining sample data were split into a training set (80%) and a test set (20%). Before Stable Selection, features with >50% missing values were excluded, and remaining missing features were imputed using median imputation. Low-variance features were removed. In categorical mode, logistic regression was used to identify signatures capable of distinguishing between the BMI groups, while in continuous mode, LASSO was used to identify signatures that change with the continuous BMI score. Stability selection over 500 bootstrap iterations. Finally, signatures identified in either mode were aggregated to define a consensus signature for downstream analysis. This essentially allows us to reduce the hypothesis-testing space, which would otherwise have been an insurmountable hurdle. Importantly, STABL was performed as a step in supervised signature prioritization rather than for statistical inference. Subsequent Bayesian hierarchical modeling was performed within the selected feature space.

**Bayesian Hierarchical Modeling Using Cellular Features**

A Bayesian hierarchical modeling framework in PyMC was used to quantify the magnitude and uncertainty of BMI-associated changes in signatures. The model estimates effect sizes while leveraging partial pooling to share information across biologically related features within each cell type. We built two model a categorical model to capture non-linear associations across

BMI groups (Supplementary Figure 1A and B), and we modeled the expected signature change $\mu_{ij}$ as:

$$\mu_{ij} = \alpha_j + \gamma_{p[i]} + \beta_{k[j]}^{\text{Overweight}} \cdot I(BMI_i \in \text{Overweight}) + \beta_{k[j]}^{\text{Obese}} \cdot I(BMI_i \in \text{Obese})$$

Where, $I$ is the indicator function, $\alpha_j$ is the baseline abundance for normal weight, $\beta_{k[j]}^{\text{Overweight}}$ is the effect of being overweight relative to normal and $\beta_{k[j]}^{\text{Obese}}$ is the effect of being obese relative to normal. The continuous model assesses linear changes in signature with increasing BMI.

$$\mu_{ij} = \alpha_j + \gamma_{p[i]} + \beta_{k[j]}^{\text{Slope}} \cdot BMI_{std,i}$$

Where $\beta_{k[j]}^{\text{Slope}}$ means the change in signature with respect to the change in BMI, $BMI_{std,i}$ is the standardized BMI data, $\alpha_j$ is the expected signature expression of a patient with average BMI. Where $\gamma_{p[i]}$is the patient-level random intercept for the patient $p$corresponding to observation $i$, modeled as:

$$\gamma_p \sim \text{Normal}(0, \sigma_{\text{patient}})$$
$$\sigma_{\text{patient}} \sim \text{HalfNormal}(0.50)$$

The hierarchical structure consisted of two levels: (i) cell type level effect representing changes at the cellular level and (ii) signature/feature level changes capturing signature-specific variations. A non-centered parameterization was used to improve sampling efficiency and to avoid sampler divergences. To account for the repeated-measures structure of the data, wherein each patient contributes one observation per signature across multiple cell types, a patient-level random intercept $\gamma_{p[i]}$was incorporated into both models using non-centered parameterization:

$$\gamma_p = \tilde{\gamma}_p \cdot \sigma_{\text{patient}}, \tilde{\gamma}_p \sim \text{Normal}(0,1), \sigma_{\text{patient}} \sim \text{HalfNormal}(0.50)$$

This term partitions systematic between-patient baseline variation in deconvolved Z-scores from BMI-associated effects, preventing pseudoreplication from inflating posterior certainty. The prior scale of 0.50 was applied uniformly across all three compartments and was chosen to allow meaningful patient-level variation while maintaining regularization. The posterior standard deviation of patient intercepts was examined as a diagnostic to confirm that the model meaningfully captured patient-level variation. Regularising priors were applied to reduce overfitting and stabilize inference. Effect sizes were modeled using zero-centered normal priors, while feature-level variability was constrained using half-normal priors. Prior scales were specified for each compartment to reflect differences in signal magnitude and variability.

*Supplementary Table 01: Regularization priors in each compartment:*

| ***Compartment*** | ***celltype_sigma*** | ***feature_sigma*** | ***patient_sigma*** | ***baseline_sigma*** | ***obs_sigma*** |
|---|---|---|---|---|---|
| *Non-Immune* | 0.20 | 0.30 | 0.50 | 1.5 | 1.0 |
| *Immune Coarse* | 0.25 | 0.40 | 0.50 | 1.5 | 1.0 |
| *Immune Fine* | 0.18 | 0.28 | 0.50 | 1.5 | 1.0 |

Posterior distributions were approximated using No-U-Turn Sampler (NUTS) implemented in PyMC. Each of four distinct Markov chains had 2,000 tuning steps and 2,000 sampling draws (Target Acceptance $=$ 0.99). Convergence was assessed using the Gelman–Rubin statistic ($\hat{R} < 1.01$) and Effective sample sizes (ESS) were evaluated for key hierarchical parameters, with values >400 considered acceptable and >1000 indicating strong sampling efficiency. Convergence diagnostics, including trace plots, $\hat{R}$, and ESS, were computed using ArviZ.

Convergence diagnostics for all compartments and comparisons are in Supplementary Table X. E-BFMI values ranged from 0.787 to 0.981 across all chains and compartments, all above the 0.3 threshold for adequate HMC exploration. We treat any comparison with Gelman–Rubin statistic above 1.01 as exploratory; none of these informed primary biological conclusions.

Three comparisons failed this criterion. The OW vs N comparison in the non-immune categorical compartment reached R-hat = 1.012; we use the OB vs N comparison from that compartment as the primary finding instead. The OW vs N comparison in the immune coarse categorical compartment was more problematic ($\hat{R}$ = 1.029 and ESS of 163–193), and we treat it as exploratory throughout. The BMI slope in the continuous immune fine compartment did not fully converge ($\hat{R}$ = 1.017), and we do not draw primary conclusions from it. All other comparisons met convergence criteria ($\hat{R}$ < 1.01, ESS > 400).

We used a combination of Highest Density Intervals (HDI) and a Region of Practical Equivalence (ROPE) to evaluate statistical credibility and practical significance. Statistical credibility was defined by the 95% Highest Density Interval (HDI) excluding zero. Practical significance was determined using multiple thresholds $Categorical\ ROPE = [0.1, 0.2, 0.3, 0.5]$; $Continues\ ROPE = [0.05, 0.1, 0.15, 0.2, 0.3]$ on the standardized effect size scale. For the continuous model, this corresponds to the slope of change of signature with BMI, and for the categorical model, it represents group-level changes under comparison. Effects were categorized as HDI and ROPE credible features (P(|effect| > 0.2) > 0.95, HDI≠0), denoted as “★”, and only HDI credible (HDI≠0) denoted as “ ∘”. Convergence diagnostics for all compartments and comparisons are summarised in the table below:

*Supplementary Table 02: Convergence diagnostic per compartment:*

| **Model & Compartment** | **Comparison** | **R-hat max** | **R-hat range** | **ESS min** | **ESS max** | **E-BFMI range** | **Mean SNR** | **Converged** |
|---|---|---|---|---|---|---|---|---|
| Categorical Non-Immune | OW vs N | 1.012 | 1.000–1.012 | 736 | 6,966 | 0.787–0.882 | 2.066 | Partial† |
| Categorical Non-Immune | OB vs N | 1.004 | 1.000–1.004 | 1,133 | 6,259 | 0.787–0.882 | 1.242 | Yes |
| Categorical Immune Coarse | OW vs N | 1.018 | 1.015–1.018 | 259 | 288 | 0.896–0.981 | 1.900 | No‡ |
| Categorical Immune Coarse | OB vs N | 1.002 | 1.001–1.002 | 1,049 | 1,221 | 0.896–0.981 | 2.102 | Yes |
| Categorical Immune Fine | OW vs N | 1.005 | 1.000–1.005 | 668 | 10,070 | 0.865–0.941 | 1.634 | Yes |
| Categorical Immune Fine | OB vs N | 1.006 | 1.000–1.006 | 940 | 13,147 | 0.865–0.941 | 1.103 | Yes |
| Continuous Non-Immune | BMI slope | 1.002 | 1.000–1.002 | 1,197 | 6,763 | 0.820–0.841 | 1.514 | Yes |
| Continuous Immune Coarse | BMI slope | 1.005 | 1.005–1.005 | 386 | 407 | 0.828–0.900 | 1.847 | Partial§ |
| Continuous Immune Fine | BMI slope | 1.017 | 1.000–1.017 | 625 | 9,479 | 0.836–0.921 | 1.075 | No¶ |

† Nine of 14 cell types reached Primary status; fibroblasts, pericytes/SMC, islet endocrine, Schwann, and tumor classical were Marginal (R-hat > 1.01). Results from Marginal cell types in this comparison are interpreted with caution.

‡ All eight cell types were Marginal (R-hat 1.015–1.018, ESS 259–288). No feature-level credibility claims are drawn from this comparison; OW vs N cell-type posterior means are reported as directional signals only.

§ Three of eight cell types reached Primary status (CD8 T General, T Cells, Basophils); the remaining five were Marginal. Credible feature-level results are reported only from Primary cell types.

¶ Ten of 20 cell types reached Primary status; the remaining ten were Marginal. Credible feature-level results are reported only from Primary cell types. The max R-hat of 1.017 reflects poor mixing in Marginal cell types and does not affect inference from Primary cell types.

#### Interactome analysis

##### Implementation of the TimiGP Framework

An investigation of cell gene pair interactions was conducted utilizing the TimiGP framework (https://github.com/CSkylarL/TimiGP). It describes cell-cell communication networks and functional states based on bulk RNA-seq expression data. Analysis was confined to normal-weight and overweight patient groups, as the obese cohort lacked sufficient sample size for reliable statistical modeling (n = 18).

TimiGP offers three unique collections of immune signatures: (1) Bindea et al. 2013 signatures, encompassing 28 immune cell types from colorectal cancer immune landscape analysis, (2) Newman's LM22 signature with 22 distinct immune cell types (3) Zheng et al. 2021 pan-cancer T cell signatures obtained from single-cell RNA sequencing across 21 cancer types. These signature sets provide comprehensive insights into immune cell interaction dynamics and the prognostic privileges they offer. The analysis is done as per the protocol previously published (3).

##### Cox Regression and Selection of Gene Pairs

Bulk RNA-seq expression data were log-transformed and normalized by geometric mean normalization with the TimiPreProcess tool. Gene pairs were created from signature-derived marker genes and analyzed using Cox proportional hazards regression with survival and clinical outcome data. The first Cox regression selection of gene pairs relied on raw p-values ($p<0.01$) to determine prognostic gene pairs having survival relevance. The top 5% of gene pairs, ranked by p-value < 0.01, were selected for subsequent network analysis and cell-cell interactions.

##### Enrichment of Cell-Cell Interactions

Chosen gene pairings were subjected to enrichment analysis utilizing the TimiEnrich function with Benjamini-Hochberg FDR correction (adjusted p-value <0.05) to discern important cell-cell interaction modules. Background gene pair distributions were determined using the TimiBG function, whereas cell pair definitions were generated using TimiCellPair with multi-core processing. Permutation-based false discovery rate estimation (n=100 iterations) was performed using TimiPermFDR to assess interaction significance and mitigate multiple-testing artifacts.

**Network Development and Assessment**

Cell interaction networks were established using the TimiCellNetwork tools, based on annotations from the signature dataset. Favorability ratings were computed using TimiFS to rank cell types by their prognostic relevance and interaction centrality within the immune ecosystem. Network visualization utilized chord diagrams and dot plots to depict interaction intensities and prognostic correlations among BMI groupings. Using the network data, we constructed an interaction network using Cytoscape V3.10.4.

**Virtual Multiplex Immunofluorescence using GigaTIME**

GigaTIME is a multimodal image-to-image translator (https://github.com/prov-gigatime/GigaTIME) with a NestedUNet (UNet++) architecture trained on a pan-cancer Providence dataset comprising 14,256 patients across 24 cancer types, including pancreatic cancer, generating 23-channel mIF images from standard H&E input. The model is available via Hugging Face (https://huggingface.co/prov-gigatime/GigaTIME).

*Data acquisition:* Whole-slide images for CPTAC-PDAC patients were obtained from DICOM files from the NCI Imaging Data Commons (IDC) using the idc-index (V23) python package. For each patient, the full resolution DICOM series was identified and opened using WsiDicom.open() from the wsidicom library.

*Tissue detection:* Each slide was scanned with non-overlapping 512×512-pixelk tiles extracted at base resolution. Tiles with a mean pixel density below 220 were retained as tissue positive; tiles above this threshold were classified as background and excluded.

*Section detection and selection:* Many CPTAC-PDAC slides contained multiple sections and disconnected tissues. Disconnected sections were isolated by a connected component analysis on the tile grid. Tiles separated by two grid steps (approximately 1024 pixels) were considered part of the same section, allowing occasional missed background tiles within a real tissue block. Sections below a minimum size of 50 tiles were excluded as edge artifacts, and sections with tissue density below 0.30 (defined as the number of tissue tiles divided by the total grid area) were excluded as parse debris. Among the retained patient sections, the most tumor-representative sections were selected by running GigaTIME inference on 30 randomly sampled 256×256-pixel patches per section and computing the mean CK channel density as a proxy for tumor epithelial content. Sections below a minimum CK channel density (minimum threshold < 0.01) were discarded. The section with the highest CK density score was selected for full inference.

*Inference:* Full gigaTIME inference was performed on the sectioned section using 256×256-pixel windows with a stride of 128 pixels, resulting in 50% overlap between adjacent windows to ensure maximum spatial resolution. Each path was upscaled from 256 to 512 pixels using albumentations Resize with ImageNet normalization before model input, and the resulting 512×512-pixel logic output was downscaled back to 256×256 pixels using bilinear interpolation so that one output pixel corresponds to one WSI pixel at base resolution. Logits from overlapping windows were averaged at each pixel position before applying a sigmoid threshold of 0.5 to produce binary activation values. The final canvas was saved as a compressed array (23 channels × H × W pixels), with tissue bounding box coordinates recorded in WSI pixel space.

*Channels used and normalization.* The 23-channel output follows the standard GigaTIME panel. Channels TRITC and Cy5 were used to represent the fluorescence background and were excluded. DAPI normalization (marker density divided by DAPI density) was applied to correct for variable tissue area across patients.

*Spatial Metric Computation:* Stromal CD9-to-tumor boundary distance: Individual CD8+ blobs were identified by connected component analysis (*scipy.ndimage.label*), and blobs less than 4 pixels were excluded as noise. Centroids were computed (using *scipy.ndimage.center_of_mass*) and filtered to retain only those falling strictly within Transgelin+ pixels (stromal-trapped CD8+ T cells). The Euclidean distance from each retained centroid to the nearest CK+ pixels was computed using a k-d tree (*scipy.spatial.cKDTree*). The per-patient metric is the mean distance across all stromal-trapped CD8 centroids. Pixel sets were subsampled to a maximum of 5,000 each using a fixed random seed.

*Stromal area fraction:* Computes as the count of Transgelin+ pixels divided by the tota;l canvas pixel count.

*Spatial co-localization:* The Sørensen–Dice coefficient between marker pairs was computed as 2× the intersection count divided by the sum of the individual pixel counts, applied to CD34 and Transgelin masks, and to PD-L1 with CK and CD68 masks, respectively.

*Stromal CD8-to-PD-L1 distance.* Identical pipeline to the primary metric with the target changed from CK+ to PD-L1+ pixels.

*Statistical Analysis:* Associations with continuous BMI were assessed using Spearman's rank correlation as the primary test and Pearson's correlation as a secondary test. A metric was considered significant if either test reached $p < 0.05$; trends were noted at $p < 0.10$. Categorical group comparisons used the Kruskal-Wallis test with the Benjamini-Hochberg correction across three pairwise comparisons per metric. Pixel-level quantitative accuracy against paired multiplex immunofluorescence ground truth in this CPTAC-PDA cohort has not been established.

#### Code repository

All code, including single-cell reference construction, Bulk analysis, BayesPrism deconvolution, Bayesian Hierarchical model building, and TimiGP analysis, is available at the GitHub repository (https://github.com/arunviswanathan91/obese-model). All data generated by the model is available at https://obese-pdac-model.streamlit.app/. All analyses were executed in Google Colab using a CPU for normal analysis and an A100 GPU for gigaTIME.
